# A mouse model of PTEN Hamartoma Tumour Syndrome reveals that loss of the nuclear function of PTEN drives macrocephaly, lymphoid overgrowth, and late-onset cancer

**DOI:** 10.1101/2025.06.20.660297

**Authors:** Priyanka Tibarewal, Victoria Rathbone, Sarah E Conduit, Gala Anastasia Electra Classen, Fiona Black, Mohammad Amin Danesh, Georgia Constantinou, Zeinab Asgarian, Koujiro Tohyama, Nisha Kriplani, Virginia Alvarez Garcia, Elizabeth Foxall, Djenat Belarbi, Marie Leverve, Wayne Pearce, Mahreen Adil, Zofia Varyova, Lucia Conde, Adriana Alves, Glenn R Masson, Roger L Williams, Adrienne M Flanagan, Javier Herrero, Isra Ahmed Mohamed, Katerina Stroud, Marc Tischkowitz, Katherine Lachlan, Cheryl L Scudamore, Mark G H Scott, Nicholas R Leslie, Nicoletta Kessaris, Bart Vanhaesebroeck

**Affiliations:** Cancer Institute, University College London, UK; Wolfson Institute of Biomedical Research, University College London, UK; Department of Physiology, School of Dentistry, Iwate Medical University, Yahaba, Japan; Institute of Biological Chemistry, Biophysics and Bioengineering, Heriot Watt University, Edinburgh, UK; Université de Paris, 75014 Paris, France; Institut Cochin, INSERM U1016, 75014 Paris, France; CNRS, UMR8104, 75014 Paris, France; MRC Laboratory of Molecular Biology, Cambridge, UK; Research Department of Pathology, University College London, UK; Cellular and Molecular Pathology, Royal National Orthopaedic Hospital, Stanmore, UK; Cambridge University Hospitals NHS Foundation Trust, Cambridge, UK; Department of Genomic Medicine, National Institute for Health Research Cambridge Biomedical Research Centre, University of Cambridge, Cambridge, UK; Wessex Clinical Genetics Service, University Hospital Southampton, Princess Anne Hospital, Southampton, UK; Human Development and Health, Faculty of Medicine, University of Southampton, Southampton, UK; Exepathology, Exmouth, UK

**Keywords:** PTEN, PI3K, PHTS, overgrowth, mouse model, cancer, macrocephaly, germline, ASD

## Abstract

PTEN Hamartoma Tumour Syndrome (PHTS) is a rare disorder characterized by germline heterozygous mutations in the PTEN tumour suppressor gene, leading to multi-organ/tissue overgrowth, autism spectrum disorder and increased cancer risk. PHTS individuals display heterogeneity in phenotypes, which has been linked in part to the diverse genetic alterations in the *PTEN* gene and the multifaceted functions of this protein. Indeed, while PTEN primarily functions as a PIP_3_ lipid phosphatase in the cytosol, regulating PI3K/AKT signalling, a pathway commonly deregulated in cancer, it also plays crucial roles in maintaining chromosomal stability through nuclear activities such as double strand (ds) DNA damage repair. Recent studies have identified a subset of missense PHTS variants that cause nuclear exclusion of PTEN, impairing its nuclear functions. Here, we present our findings from one such pathogenic variant, *PTEN-R173C*, frequently found in PHTS and somatic cancers. Using cell biological and mouse modelling approaches, we show that PTEN-R173C has higher PIP_3_ phosphatase activity than wild-type PTEN, resulting in effective regulation of canonical PI3K/AKT signalling. However, PTEN-R173C is unstable and excluded from the nucleus. Aligning with their near normal PI3K/AKT signalling, *Pten^+/R173C^*mice display a low incidence of solid tumours compared to *Pten^+/-^*mice. *Pten^+/R173C^* mice also exhibit lymphoid hyperplasia and macrocephaly which correlates with compromised nuclear functions of PTEN-R173C. That nuclear functions are compromised is demonstrated by reduced dsDNA damage repair in *Pten^+/R173C^*mice. Integrating PHTS patient data with findings from our mouse model, our study indicates that nuclear dysfunction of pathogenic *PTEN* variants is a key factor in predicting the onset of the different PHTS-associated phenotypes. We speculate that late-onset cancer in individuals with nuclear-excluded PTEN results from genetic alterations unrelated to PTEN itself, facilitated by impaired PTEN-mediated dsDNA damage repair.

## INTRODUCTION

PTEN is a ubiquitously expressed tumour suppressor^1–3^ that shows complete loss of expression in some cancers (e.g. prostate cancer and glioblastoma) and overrepresentation of missense variants in other tumours (e.g. endometrial cancer)^1,4^. Heterozygous germline variants in PTEN cause PTEN Hamartoma Tumour Syndrome (PHTS), a rare autosomal dominant disorder. A recent study puts the prevalence of PHTS as 1:8000-13,000^5^ which is 10-20 fold higher than previous estimates^6^. Clinical features of PHTS include macrocephaly, developmental delay (DD), autism spectrum disorder (ASD), dermatological pathologies, gastrointestinal (GI) polyps, immune dysfunction, vascular anomalies. Individuals with PHTS also face a significantly increased risk of developing cancers particularly of the breast, endometrium and thyroid^7,8^, with emerging evidence for ovarian cancer^9^.

PTEN acts in both the cytosol and nucleus. Cytosolic functions of PTEN predominantly include lipid phosphatase activity on phosphatidylinositol(3,4,5)trisphosphate (PIP_3_) and phosphatidylinositol(3,4)bisphosphate (PI(3,4)P_2_), thereby antagonizing PI3K/AKT signalling^10^. PTEN also has protein-phosphatase activity, reducing phosphorylation of Ser/Thr/Tyr on proteins such as PTK6, DREBRIN, DVL2, FAK and SHC^11^, in addition to auto-dephosphorylating its T366 site^12^. Nuclear PTEN has been implicated in cell cycle control, dsDNA repair, maintenance of chromatin structure, genome integrity and transcriptional control^13,14^.

The functional core of the PTEN protein consists of a catalytic (amino acids (AA) 16-185) and a C2 (AA 186-350) domain^15^, with mutations in different regions differentially impacting PTEN expression and function. 65% of PHTS patients express *PTEN* variants such as deletions, DNA insertions or non-sense mutation leading to STOP codons (such as R130X and R233X) that result in PTEN truncation and are therefore predicted to cause heterozygous loss-of-expression. The remaining PHTS patients carry heterozygous missense mutations in the phosphatase or C2 domain, leading to expression of PTEN proteins with altered function or subcellular distribution^16,17^. These include mutations in the catalytic pocket resulting in loss of all catalytic activity [e.g. C124 (C124S/R), R130 (R130G/Q/L) and G129 (G129R)]^15^ or selective loss of PIP_3_ phosphatase activity, while retaining protein phosphatase activity (e.g. G129E)^10^. Missense mutations leading to selective loss of protein phosphatase activity (Y138L/C) have also been described, although these have not been reported in PHTS^12,18^. Recent reports have shown that a subset of PTEN variants in PHTS (including K289E, D252G, F241S, I101T and others) are excluded from the nucleus and therefore lack all PTEN nuclear functions^14^.

The PIP_3_ phosphatase activity of PTEN has long been regarded as the main and critical factor in PHTS pathogenesis, yet other lipid phosphatase-independent functions of PTEN are now also emerging as important^14^. Various studies have attempted to establish genotype-phenotype associations in PHTS. Limited patient data make it challenging to ascertain the impact of individual variants in driving patient phenotypes. This challenge has been partially overcome by classifying the variants based on their position in the *PTEN* gene and their impact on PTEN expression and function^19–23^. Recently, a large paediatric and adult patient cohort study revealed that missense variants are associated with early disease onset, with DD and macrocephaly being the reason for early diagnosis, whereas *PTEN* truncation variants are associated with late onset disease, with cancer being the reason for PHTS diagnosis at later stages in life^16^. However, patient cohort studies have been limited by the variability in age and sex of the individuals, one of the key challenges in rare disease research.

Genetically modified mice have helped overcome some of the challenges in PTEN research by allowing assessment of PTEN functions in an endogenous tissue context^24^. They also serve as a platform for testing potential treatments, such as PI3K pathway inhibitors, aimed at mitigating the effects of PTEN loss. An example are mice with heterozygous loss of PTEN expression (*Pten^+/-^*), representing *PTEN* variants that result in loss of expression and function. This mouse model recapitulates many aspects of human PHTS remarkably well in terms of cancer development and lymphoid and brain overgrowth^25,26^.

Here we report the creation and characterisation of a novel mouse model for PHTS, carrying a PTEN-R173C (NM_000314.8(PTEN):c.517C>T (p.Arg173Cys)) missense variant in the germline. R173 is located in the TI (Thr-Ile) loop in the catalytic domain of PTEN important for PTEN catalytic activity^15^. Mutation of R173 to Cys or His in similar ratios are observed in PHTS and sporadic cancers, with conflicting reports on catalytic activity impact, from having been reported active to inactive^27–29^. R173 variants are frequently found in multiple patient cohort studies with 34 PHTS patients been reported in literature to the best of our knowledge, and an additional four new patients identified with this variant in the UK (**Supplementary Table 1**). Most patients with R173 variants received PHTS diagnosis at young age due to presentation of macrocephaly, DD and cutaneous pathology. Adult patients additionally presented with breast cancer, benign and malignant tumours of the skin, thyroid and ovary, endometrial polyps and fibroids, GI polyps, with malignant disease onset much later in life (over the age of 50 years). Vascular malformations, mainly haemangiomas, were also seen in adult patients (**Table 1** and **Supplementary Table 1**).

**Table 1.**
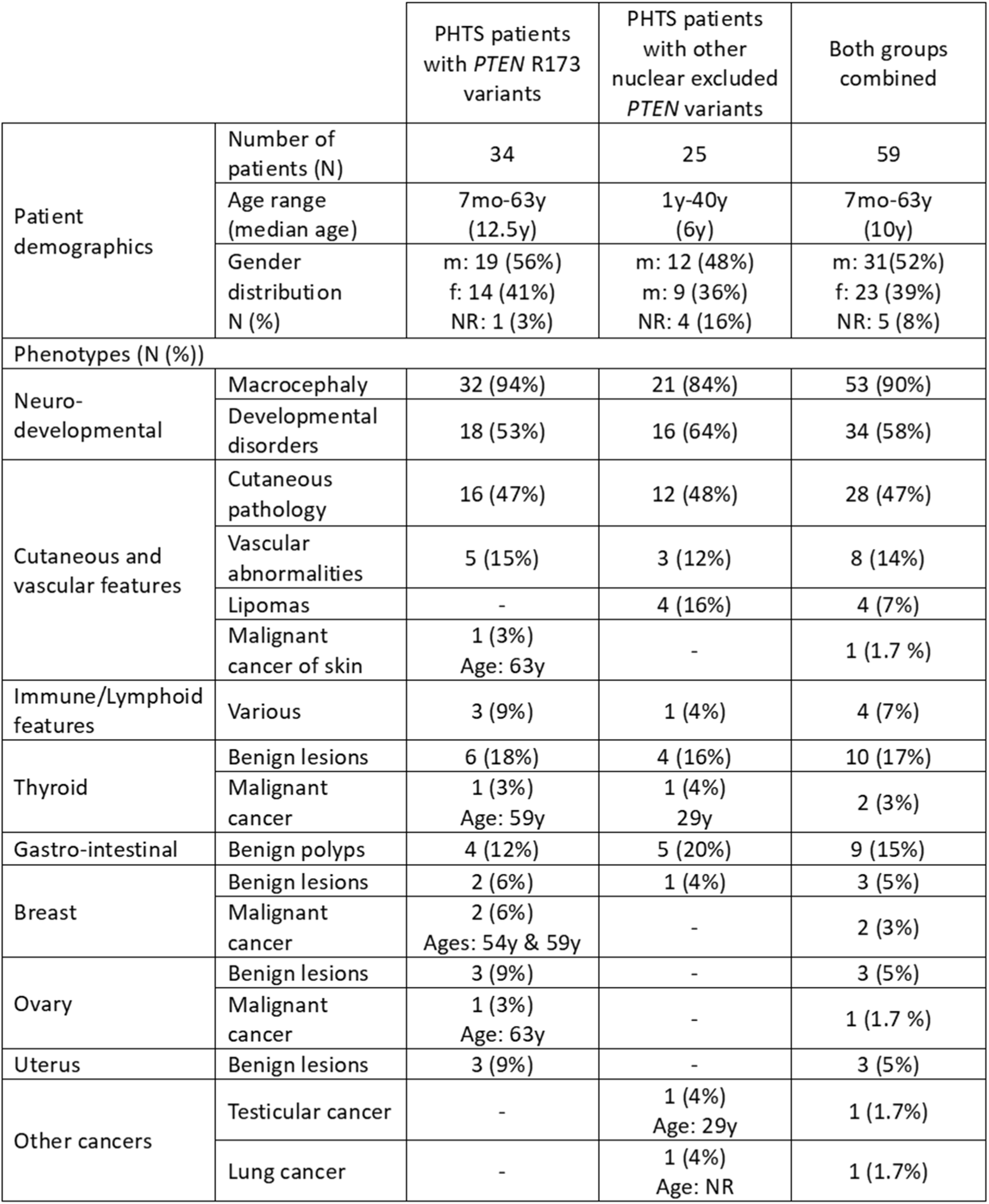
Clinical phenotypes of PHTS patients with *Pten-R173* variants and other nuclear excluded *Pten* variants. mo, age in months; y, age in years; m, males; f, females, NR, not recorded. Phenotypic categories include the following: <u>Developmental disorders</u>; global developmental delay, delay in speech development, mental retardation, autism spectrum disorder, special educational needs. <u>Cutaneous</u> <u>pathology</u>; hamartomas, tricholemmoma, mucocutaneous lesions, acral keratoses, palmoplantar keratoses, oral papillomas, penile freckling, facial papules, café-au-lait spots, skin tags. <u>Vascular abnormalities</u>; haemangiomas, venous malformations, arterio-venous malformations. <u>Immune/</u> <u>lymphoid features</u>; recurrent upper respiratory tract infections, hypogammaglobulinemia, enlarged tonsils and adenoids, lymphadenopathy, splenomegaly. <u>Thyroid benign lesions;</u> cyst, nodules, adenomas. <u>Thyroid malignant cancer</u>; follicular cancer, papillary-follicular thyroid cancer. <u>Breast</u> <u>benign lesions</u>; fibrocystic disease of the breast, fibroadenomas, benign breast disease. <u>Breast</u> <u>malignant cancer</u>; ductal breast carcinoma, invasive ductal carcinoma and ductal carcinoma in-situ of the breast. <u>Ovary benign lesions</u>; ovarian cysts, benign neoplasm. <u>Ovary malignant cancer</u>; malignant neoplasm of the ovary. <u>Uterus benign lesions</u>; uterine fibroids, endometrial polyp. The phenotypes of individual patients are listed in Supplementary Tables 1 and 2.

Here, we first revisited the impact of the R173 mutation on PTEN expression and function using biochemical and cell-based models, with a focus on PTEN-R173C. We show that PTEN-R173C has reduced protein stability but retains its ability to dephosphorylate PIP_3_ and therefore regulate the canonical PI3K/AKT pathway in the cytosol. However, we find that this PTEN mutant is largely excluded from the nucleus.

Several mouse models have been developed to distinguish the relative importance of the cytosolic *versus* nuclear functions of PTEN (reviewed in^14^). For instance, *Pten^m3m^*^4^ mice, engineered by Eng *et al*., feature non-naturally occurring PTEN mutations that result in a predominantly cytosolic PTEN distribution without affecting its PIP_3_ phosphatase activity. These mice have been extensively studied for the impact of the nuclear-cytoplasmic partitioning of PTEN on neuronal and immune phenotypes, although their tumour spectrum has not been reported^30–32^. Other models, expressing PTEN missense variants found in PHTS such as F341V^33^, and non-pathogenic mutations such as K13R, Y240F and D384V^34,35^, that do not affect the catalytic functions of PTEN but impact its nuclear/sub-cellular localisation, have also been developed, but there is limited or no patient data for these individual variants (**Table 1** and **Supplementary Table 2**).

We created heterozygous germline *Pten*^R173C^ knock-in mice (*Pten*^+/R173C^) as a unique model of a frequent pathogenic PHTS PTEN variant representing nuclear-excluded PTEN missense variants. We examined the consequences of this mutation on embryonic development, glucose metabolism, development of PHTS-relevant cancers as well as immune, neurological and behavioural phenotypes. By integrating our findings with phenotypic data from patients for this variant, and by comparison of reported data on the *Pten^m3m4^*mice, we delineate the role of the nuclear functions of PTEN in the manifestation of PHTS phenotypes.

## RESULTS

### Increased PIP_3_ phosphatase activity but reduced protein stability of PTEN-R173 mutant proteins upon transient expression in mammalian cells

Conflicting data on the catalytic activity of PTEN-R173 variants have been published. PTEN-R173 mutant proteins, expressed in *E. coli*, have been reported to be catalytically inactive^27^ however, mutant PTEN proteins expressed in non-mammalian systems often fail to fold properly, resulting in lack of activity^17^. Consistent with this, in our hands, PTEN-R173C protein expressed in *E. coli* lacked *in vitro* PIP_3_ lipid phosphatase activity (**Fig. S1**) and when expressed in Sf9 insect cells precipitated upon purification ^36^ (data not shown), rendering it unsuitable for use in *in vitro* assays.

A commonly used assay to determine the ability of PTEN and its mutants to regulate PI3K/AKT signalling involves transfection into the PTEN-null U-87 MG glioblastoma cell line (further referred to as U87) and assessment of basal AKT phosphorylation and cell proliferation^18^. Transient lentiviral expression of untagged PTEN-R173C in U87 revealed consistently lower expression levels compared to PTEN-WT (**Fig. 1A**, left panel), despite similar mRNA expression levels (**Fig. 1A**, right panel). Similar results were seen with PTEN-R173H, suggesting that other PTEN-R173 mutant proteins are less stable than PTEN-WT.

**Figure 1.**
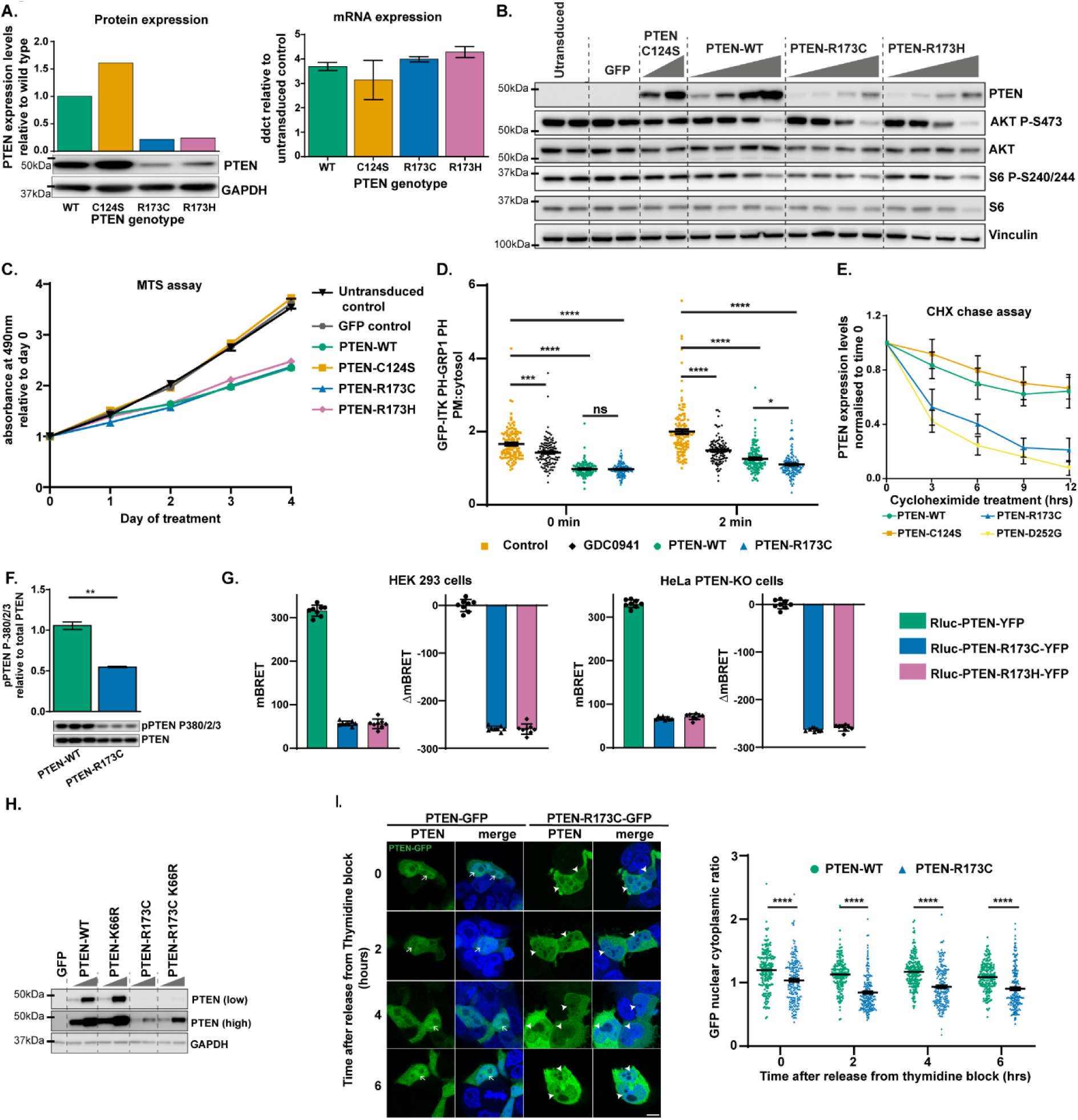
Characterisation of PTEN-R173C/H. (**A-C**) Transient lentiviral expression in U87 cells of PTEN-WT or the indicated PTEN mutants at fixed (A and C) or increasing concentrations of lentiviral particles (B). 48 h after transduction (A and B), cells were either lysed and PTEN expression and (B) AKT P-S473 and S6 P-S240/244 assessed by immunoblotting, or (A) qPCR analysis was performed to determine PTEN mRNA levels (mean ± SEM, n=3, right panel). The PTEN-C124S mutant was used as a control. (**C**) Cell proliferation was determined by MTS assay over 4 days. Graph shows the mean ± SEM of absorbance at 490 nm relative to day 0 from 10 replicate wells of a representative experiment from n=2. Immunoblots of PTEN protein and AKT P-S473 from the cells in (C) are shown in **Fig. S2A**. (**D**) U87 cells transduced with lentivirus expressing a PIP_3_ biosensor, with or without PTEN-WT or PTEN-R173C, were pre-treated ± GDC-0941 (1 µM) for 1 h, followed ± insulin treatment (100 nM) for 2 min. The cells were stained with antibodies to PTEN and imaged by confocal microscopy (images in **Fig. S2B**). Graph shows mean fluorescence intensity (MFI) of the PIP_3_ biosensor at the plasma membrane normalized to the MFI of the PIP_3_ biosensor in the cytosol. Data shown as mean ± SEM from 3 independent experiments, statistical analysis performed using 2-way ANOVA. Representative immunoblots for PTEN protein and AKT P-S473 from the cells are shown in **Fig. S2C**. (**E-F**) Transient lentiviral expression of PTEN wild-type (WT) or the indicated PTEN mutants in U87 cells. (E) The cells were treated with cycloheximide and PTEN protein expression levels determined by immunoblotting at the indicated time points. Graph shows PTEN protein levels normalized to time 0, data shown as mean ± SEM, n=6. Representative blots shown in **Fig. S3A**. (F) Phosphorylation of PTEN on S380/T382/T383 was determined by immunoblotting. Quantification shows mean ± SEM, n=3, statistical analysis was performed using Unpaired Student’s t-test. (**G**) mBRET and ΔmBRET values, obtained in live HEK-293T and HeLa PTEN KO cells, transfected with Rluc-PTEN-WT-YFP, Rluc-PTEN-R173C-YFP or Rluc-PTEN-R173H-YFP. Graphs represent mean ± SD of n=8 points from a representative experiment from 3 independent experiments. (**H**) Transient lentiviral expression of PTEN-WT or the indicated PTEN mutants in U87 cells. Protein extracts from these cells were used for immunoblotting to assess PTEN protein levels. (**I**) Lenti-X™ 293T cells expressing GFP-tagged PTEN-WT or PTEN-R173C were arrested in the G1/S phase of cell cycle by double-thymidine block and imaged by confocal microscopy at the indicated time points after release from thymidine block, for expression of PTEN-GFP. Left panel shows representative images, arrows indicate nuclear PTEN-WT or reduced nuclear PTEN-R173C. Scale bar: 10 µm. The graph shows the nuclear:cytoplasmic ratio of the mean fluorescence intensity (MFI) of the PTEN-GFP signal (mean ± SEM; n=3, statistical analysis performed using 2-way ANOVA). p-values for all statistical tests: *, p< 0.05; **, p<0.01; ***, p<0.001; ****, p<0.0001.

Expression of a dose range of PTEN proteins in U87 revealed that, compared to PTEN-WT, lower levels of PTEN-R173C or PTEN-R173H effectively downregulated AKT P-S473 and S6 P-S240/244 phosphorylation (**Fig. 1B**) and suppressed U87 proliferation (**Fig. 1C and S2A**). We also assessed the impact of PTEN-R173C expression in U87 cells on recruitment of a PIP_3_-biosensor protein to the plasma membrane. In both serum-starved or insulin-stimulated cells, the biosensor plasma membrane to cytosol ratio (PM:cytosol) was significantly reduced upon treatment with the pan-class I PI3K inhibitor GDC-0941 or upon expression of PTEN-WT or PTEN-R173C, with the strongest decrease upon PTEN-R173C expression (**Fig. 1D and S2B**), despite its lower expression levels compared to PTEN-WT (**Fig. S2C**).

Taken together, these data suggest that in mammalian cells, PTEN-R173C is less stable but catalytically more active than PTEN-WT.

### Evidence for a more open, proteolytically-sensitive protein conformation of PTEN-R173 mutants

We next investigated possible biochemical mechanisms underlying the reduced expression of PTEN-R173C mutants in U87 cells. Cycloheximide chase experiments of U87 cells transiently transfected with PTEN expression constructs indicated a reduced protein half-life of PTEN-R173C compared to PTEN-WT (**Fig. 1E, S3A** and Ref.^17^). A recent study has shown similar findings with PTEN-R173H^37^.

PTEN protein stability and activity is regulated, in part, by phosphorylation on multiple C-terminal sites (S380/T382/T383/S385) by kinases such as CK2^38^. Such phosphorylation results in a more closed, less active and more stable protein conformation^38–42^. Upon transient expression of similar levels of untagged proteins in U87 cells, PTEN-R173C showed a ∼50% lower level of S380/T382/T383 phosphorylation compared to PTEN-WT (**Fig. 1F**). Similar data were observed using YFP-tagged Rluc-PTEN-R173C and PTEN-R173H in HEK-293T cells (**Fig. S3B**), suggesting that reduced phosphorylation may contribute to the observed instability of PTEN-R173 mutants.

Dephosphorylation of S380/T382/T383/S385 on PTEN releases the inhibitory effect of these phosphorylated residues on the C2 domain of PTEN, leading to a more open conformation with increased membrane binding and therefore enhanced activity^38–41^. It has also been shown that, once at the membrane, the active form of PTEN becomes degraded^43^. A study using computational modelling predicted increased protein flexibility of PTEN-R173C/H mutants, leading to impaired protein stability^29^. In order to gain insight into these phenomena, we used an intramolecular bioluminescent resonance energy transfer (BRET)-based biosensor Rluc-PTEN-YFP in live HEK-293T and PTEN KO HeLa cells. This biosensor reveals dynamic changes in PTEN conformational rearrangement and function due to shifts in energy transfer between the donor/acceptor couple^44^ (**Fig. S4A-B**). Using this approach, PTEN-R173C and PTEN-R173H were found to be in a more open/flexible conformation compared to PTEN-WT (**Fig. 1G**).

We previously reported that PTEN protein dephosphorylated on S380/T382/T383/S385 is more prone to ubiquitination and degradation, and that this involves the K66 ubiquitination site in PTEN^43,45^. In line with this, when transiently expressed in U87 cells, the PTEN-R173C/K66R double mutant which cannot be ubiquitinylated, showed increased protein expression levels compared to PTEN-R173C (**Fig. 1H**).

Altogether these data indicate that the PTEN-R173C protein has a more open conformation that allows ubiquitination and degradation, leading to reduced protein levels.

### PTEN-R173 mutants are predominantly cytoplasmic and largely excluded from the nucleus

The tumour suppressor functions of PTEN have been mostly attributed to its ability to regulate the PI3K/AKT pathway, with the role of other PTEN functions such as protein phosphatase activity and nuclear functions less well-studied in this context^13^. Given that our results so far show that PTEN-R173C/H mutants are effective at regulating the PI3K/AKT pathway, we turned our attention to other PTEN functions that might explain the high mutation frequency of this site in somatic cancers and PHTS.

Analysis of the subcellular distribution of PTEN-R173C by transiently expressing C-terminally GFP-tagged PTEN-R173C or PTEN-WT in Lenti-X™ 293T cells revealed a significantly lower nuclear:cytoplasmic ratio of the GFP signal in cells expressing PTEN-R173C-GFP than in cells with PTEN-WT-GFP, suggesting that PTEN-R173C was largely excluded from the nucleus (**Fig. 1I**). Nuclear exclusion of the R173 mutants was also revealed upon expression of YFP-tagged Rluc-PTEN-WT, -R173C or -R173H in HEK-293T or PTEN KO HeLa cells (**Fig. S5**).

### Homozygous germline PTEN-R173C expression is embryonically lethal in mice

In order to study PTEN-R173C in its endogenous tissue context and to better understand its role in PHTS, we created heterozygous germline *Pten*^R173C^ knock-in mice (*Pten*^+/R173C^) by CRISPR-Cas9 technology (**Fig. S6**). Intercrosses of *Pten*^+/R173C^ mice did not yield live-born homozygous *Pten*^R173C/R173C^ mice (**Fig. 2A**). *Pten*^R173C/R173C^ embryos were almost all reabsorbed at embryonic day (E)11.5, with under-representation compared to Mendelian ratios at earlier time points (**Fig. 2A**). Intact *Pten*^R173C/R173C^ embryos were found up to E9.5 at which stage, 5 out of 9 embryos were smaller and had not undergone turning, with the remaining 4 having been reabsorbed, suggesting that *Pten*^R173C/R173C^ embryos die around E9.5 (**Fig. 2A**).

**Figure 2.**
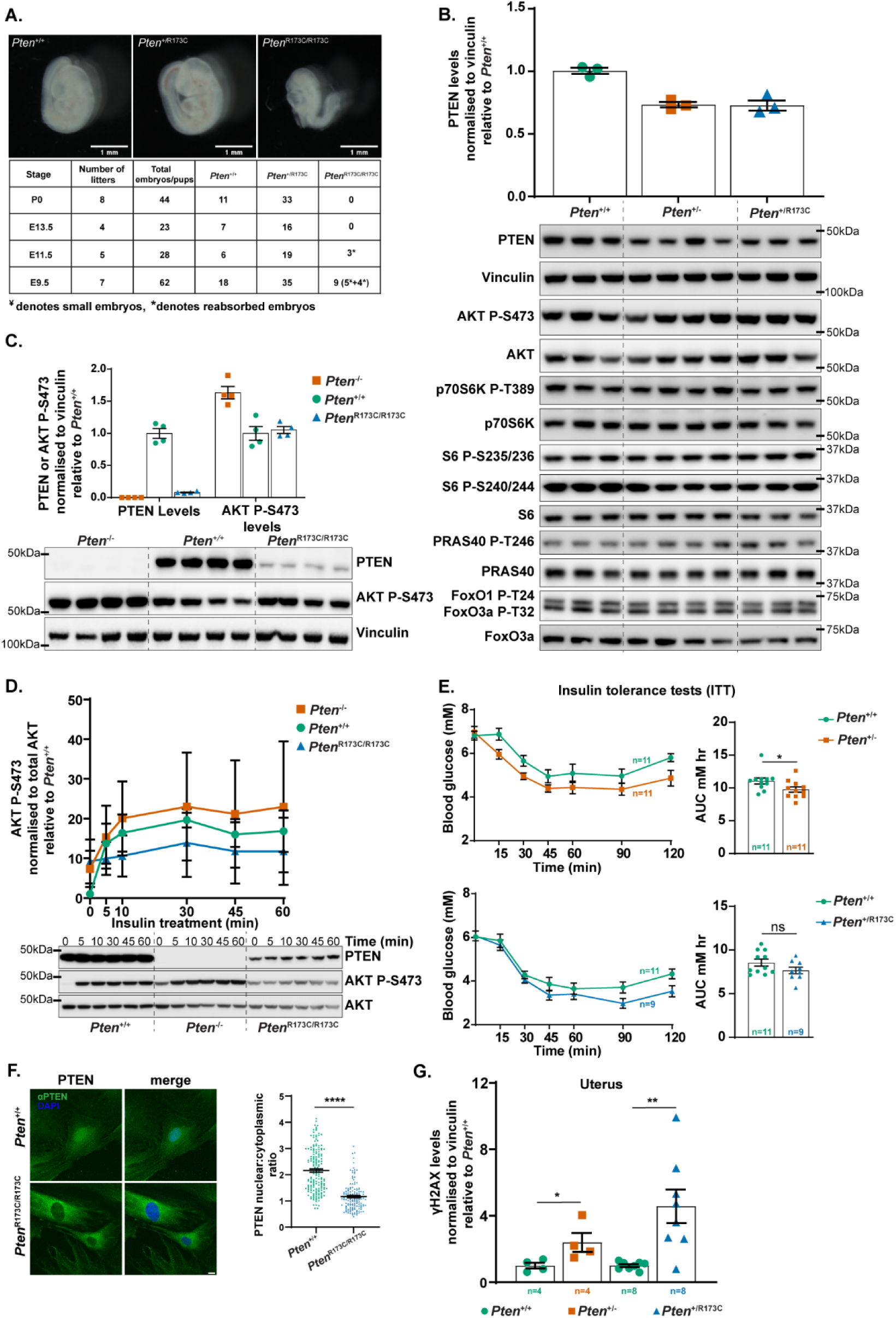
Characterisation of *Pten*^+/R173C^ mice. (**A**) Homozygous *Pten*^R173C/R173C^ mice die *in utero* after embryonic stage E9.5. Images show morphology of E9.5 *Pten*^+/+^, *Pten*^+/R173C^ and *Pten*^R173C/R173C^ embryos. Scale bar: 1 mm. The table shows the number of embryos/pups of each genotype obtained at the indicated time points. ¥ indicates embryos smaller in size and * indicates embryos that had been reabsorbed. (**B, C**) Protein extracts from MEFs of the indicated genotype were used for immunoblotting with the antibodies shown. Data shown as mean ± SEM. At least 3 independent MEF lines of each were used for (B) and 2 independent MEF lines were used for (C). (**D**) MEFs of the indicated genotype were treated with insulin (100 nM) for the indicated times and protein extracts immunoblotted with the antibodies shown. Data shown as mean ± SEM from 2 independent MEF lines for each genotype. (**E**) ITT assays on 3-month-old male mice of the indicated genotypes. XY curves show blood glucose levels post insulin injection over time. Bar graphs show mean ± SEM of measurements of Area Under the Curve (AUC). Statistical analysis performed using Unpaired Student’s t-test. (**F**) MEFs of the indicated genotype were immunostained with antibodies to PTEN (green) and DAPI (blue) and imaged by confocal microscopy. Scale bar: 10 µm. Graphs show nuclear:cytoplasmic ratio of the GFP MFI (mean ± SEM from 3 independent experiments). Statistical analysis was performed using Unpaired Student’s t-test. (**G**) 6-week-old mice of the indicated genotypes were treated with 7Gy γ-radiation and protein extracts from uterus were used for immunoblotting. Graph shows levels of γH2AX relative to littermate *Pten*^+/+^ controls. Data shown as mean ± SEM. Statistical analysis was performed using Mann–Whitney U tests. Accompanying immunoblots are shown in **Fig. S9A**. p-values for all statistical analysis: *, p< 0.05; **, p<0.01; ***, p<0.001; ****, p<0.0001. Mice used for generating MEF lines in (A-D and F) and for experiments in (E and G) were on a C57BL/6J background.

Given the embryonic lethality of homozygous *Pten ^R173C/R173C^*mice, and the notion that PTEN mutations in PHTS individuals are heterozygous, in the next series of experiments we used heterozygous *Pten^+/R173C^* mice to further investigate the impact of the PTEN R173C mutation. For these studies, we used littermate wild-type (*Pten^+/+^*) mice as controls as well as *Pten^+/-^*mice, a well-characterised model for PHTS^25,46–50^.

### PTEN-R173C expressed from its endogenous promoter is unstable but retains PIP_3_ phosphatase activity

As a cell-based model to study endogenous PTEN-R173C protein expression and function, we derived mouse embryonic fibroblasts (MEFs) of the following genotypes: *Pten*^+/R173C^, *Pten*^R173C/R173C^, *Pten*^-/R173C^ and compared these to *Pten^+/-^*, *Pten^-/-^* and *Pten^+/+^*MEFs.

PTEN protein expression levels in *Pten*^+/R173C^ and *Pten^+/-^* MEFs were 60-70% of those in *Pten*^+/+^ MEFs (**Fig. 2B**). The PTEN levels observed in *Pten*^+/-^ MEFs agree with previous studies where the *Pten*^WT^ allele was shown to express higher than 50% level of PTEN protein^51^. In *Pten*^R173C/R173C^ and *Pten*^-/R173C^ MEFs, PTEN protein levels were ∼13% and ∼2%, respectively, of those in *Pten*^+/+^ MEFs (**Fig. 2C; Fig. S7**), indicative of low protein stability of the PTEN-R173C protein, in line with our exogenous expression studies above.

We next used immunoblotting to assess PI3K pathway activation, under standard cell culture conditions and upon insulin-stimulation. *Pten*^+/R173C^*, Pten*^+/-^ and *Pten*^+/+^ MEFs growing in complete media showed similar basal levels of phosphorylation of AKT, S6 and other downstream effectors (**Fig. 2B**). Under these conditions, *Pten*^-/-^ MEFs had increased levels of AKT P-S473, which was not observed in *Pten*^R173C/R173C^ MEFs (**Fig. 2C**). In *Pten^-/R173C^* MEFs, the levels of AKT P-S473 were elevated compared to *Pten^+/+^* and *Pten*^+/-^ MEFs, but still substantially lower compared to *Pten^-/-^* MEFs (**Fig. S7**).

Upon insulin stimulation, the increase in AKT P-S473 was dampened more effectively in *Pten*^R173C/R173C^ MEFs than in *Pten*^+/+^ and *Pten*^-/-^ MEFs, despite the very low levels of PTEN-R173C expression (**Fig. 2D**).

Taken together, these data indicate that, even at low levels of expression, PTEN-R173C is still capable of effectively downregulating unstimulated and insulin-stimulated AKT P-S473 phosphorylation and thus retains overall PIP_3_ phosphatase activity.

### PTEN-R173C regulates glucose metabolism *in vivo*

PTEN plays an important role in glucose metabolism by dampening insulin signalling, with *Pten*^+/-^ mice showing enhanced acute insulin signalling and increased glucose metabolism^52–54^. PHTS patients also often show constitutive insulin sensitization^54^. In line with these reports, insulin tolerance tests on 3-month-old mice revealed enhanced insulin sensitivity in *Pten*^+/-^ mice, evidenced by lower levels of blood glucose post insulin injection compared to WT littermates (**Fig. 2E**). In contrast, WT and *Pten*^+/R173C^ mice showed a similar insulin response up to 60 min after insulin injection (**Fig. 2E**). This indicates that, at the organismal level, PTEN-R173C maintains the ability to downregulate acute insulin/PI3K/AKT signalling and therefore glucose homeostasis (**Fig. 2E**).

### PTEN-R173C expressed from an endogenous promoter is predominantly nuclear-excluded with reduced ability to regulate dsDNA damage repair

To determine the subcellular distribution of PTEN-R173C upon expression from its endogenous locus, we used immunocytochemistry and quantification of the PTEN nuclear/cytoplasmic ratio in *Pten*^R173C/R173C^ and *Pten*^-/R173C^ MEFs. These data showed that PTEN-R173C expressed from its endogenous locus was predominantly nuclear-excluded, in contrast to PTEN-WT protein which was present in both the nucleus and cytoplasm (**Fig. 2F and S8A**). Given that R173 is not located in a known PTEN nuclear localisation signal, one reason for the apparent nuclear exclusion of PTEN-R173C could be sequestration at the plasma membrane, with little protein available for nuclear import. To investigate this, we treated *Pten*^R173C/R173C^ MEFs with the pan-class I PI3K inhibitor GDC-0941, with the aim to reduce the PIP_3_ substrate levels for PTEN at the plasma membrane and to potentially restore the presence of PTEN-R173C in the nucleus. However, GDC-0941 treatment did not affect the subcellular distribution of PTEN-R173C (**Fig. S8B**), suggesting that plasma membrane PIP_3_ levels do not significantly contribute to the nuclear exclusion of PTEN-R173C.

Nuclear PTEN has numerous functions including regulation of DNA repair, transcription, chromatin structure, chromosomal stability and genome integrity by various mechanisms^13,14^. We investigated one of these nuclear functions in *Pten*^+/R173C^ mice, namely dsDNA repair. Nuclear PTEN deficiency has been shown to cause a homologous recombination (HR) defect in cells, leading to increased γH2AX expression and decreased Rad51 foci formation upon genotoxic stress such as ionizing radiation^55–57^. Ionizing radiation (IR) of *Pten*^+/R173C^ and *Pten*^+/-^ mice, followed by analysis 5 h post-IR treatment, led to a similar increase in γH2AX levels in protein extracts from uteri, kidney and liver, compared to wild-type mice (**Fig. 2G and S9)**, indicative of a similarly reduced DNA repair capacity upon heterozygous PTEN loss or heterozygous PTEN-R173C expression. This suggests that PTEN-R173C has a reduced ability to regulate DNA damage repair compared to PTEN-WT.

### Reduced cancer predisposition of *Pten*^+/R173C^ mice compared to *Pten*^+/-^ mice

Individuals with PHTS have an increased risk of cancers of multiple tissues including breast, endometrium and thyroid^4,8,58^. To gain insight into the cancer predisposition of germline *Pten*^R173C^ expression, we aged heterozygous *Pten*^+/R173C^ mice alongside *Pten*^+/-^ and *Pten*^+/+^ littermate controls. We used mice on a mixed C57Bl/6JxSv129 background, as on this background *Pten^+/-^* mice develop PHTS-relevant phenotypes, with an overlapping tumour spectrum including hyperplasia/tumours of the thyroid, endometrium, lymphoid tissue, small intestine and adenomyoepithelioma and malignant mammary tumours in the females^26^. *Pten*^+/-^ mice also develop lesions not seen in PHTS patients including, pheochromocytoma and intraepithelial neoplasia (PIN) and adenocarcinoma of the prostate^25,46–50^.

Gross morphology and development of *Pten*^+/R173C^ mice was found to be similar to *Pten*^+/+^ littermates, with a tendency for a slight increase in body weight at young age in both *Pten*^+/R173C^ and *Pten*^+/-^mice (**Fig. S10**).

Mice were aged and sacrificed at first appearance of signs of ill health or palpable masses. The reasons for humane culling of mice are listed in **Supplementary Table 3**. Under these experimental conditions, the median survival age of *Pten*^+/R173C^ mice was 585 and 302 days for male and female mice, respectively. This is significantly longer than the median survival age of 310 and 179 days for male and female *Pten*^+/-^ mice, respectively (**Fig.3A**). However, *Pten*^+/R173C^ mice had a shorter lifespan compared to wild-type mice (**Fig.3A**). This was especially the case for female *Pten*^+/R173C^ mice, which were mainly culled due to palpable masses or ill health.

**Figure 3.**
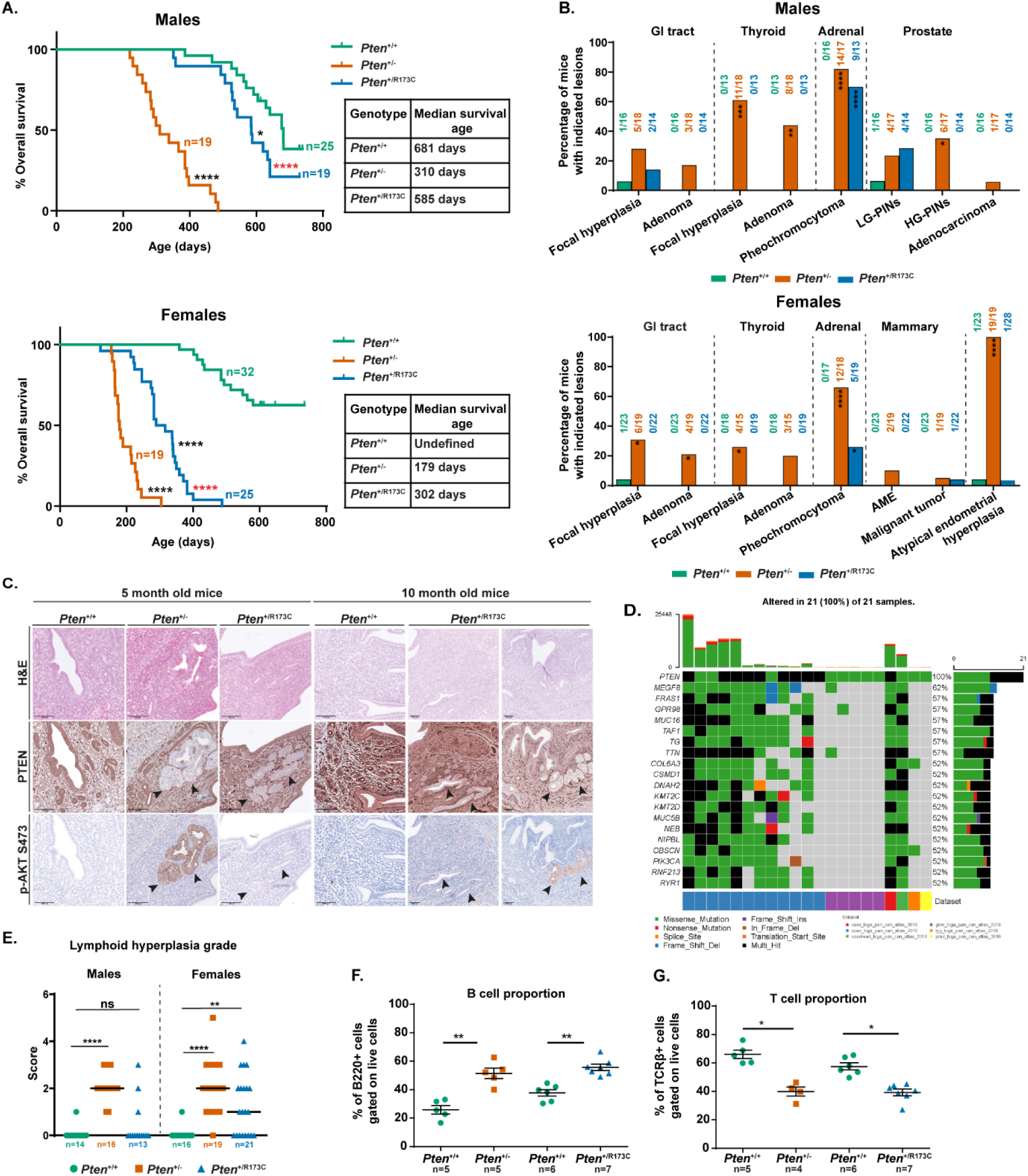
Survival and histopathological analysis of *Pten*^+/R173C^ mice. *Pten*^+/R173C^ and *Pten*^+/-^ mice on a mixed C57BL/6J x Sv129 background were allowed to age. Mice were euthanized for welfare reasons (ill health or palpable masses with a combined size of ≥ 1.4 cm^2^ surface area) or at a specified age. (**A**) Kaplan–Meier survival curves for male and female mice. Statistical analysis was performed using Log-rank (Mantel–Cox) test and Gehan–Breslow–Wilcoxon test. Comparisons were made to *Pten*^+/+^ mice (p-values shown as black stars) or *Pten*^+/-^ mice (p-values shown as red stars). Table show median survival age of the mice. (**B**) Incidence of specific tumour types in male and female mice, as assessed by histopathological analysis on all mice. Statistical analysis was performed using Fisher’s exact test. (**C**) Representative IHC images of uteri from female *Pten*^+/R173C^ and *Pten*^+/-^ and littermate *Pten*^+/+^ controls at the indicated timepoints, black arrowheads indicate loss of PTEN immunoreactivity and the corresponding AKT P-S473 immunoreactivity. (**D**) Oncoplot showing co-occurrence of mutations in the indicated genes in tumor samples harbouring PTEN-R173C mutation (cBioportal). (**E**) Grade of lymphoid hyperplasia in mice at the end of study. Graph shows scatter plot of scores from individual mice with line at median score. Statistical analysis was performed using Fisher’s exact test. (**F-G**) Flow cytometry analysis was performed on the lymph nodes of female *Pten*^+/R173C^, *Pten*^+/-^ and littermate *Pten*^+/+^ controls at 5 and 10 months of age. Graphs show the proportion of (F) B220+ B-cells and (G) TCRβ+ T-cells. Statistical analysis was performed using Mann–Whitney U test. p-values for all statistical analysis: *, p< 0.05; **, p<0.01; ***, p<0.001; ****, p<0.0001.

Histopathological analysis of selected tissues from *Pten*^+/-^ and *Pten*^+/R173C^ mice revealed very similar gross morphologies of all tissues analysed, with overall histopathology data summarized in **Fig. 3B** and **Supplementary Tables 4-7**.

*Pten*^+/R173C^ mice developed far fewer tumours than *Pten*^+/-^ mice, with occasional hyperplasia of the small intestine (2/14 mice) and low-grade PINs of prostate (4/14 mice), one case of mammary adenocarcinoma (1/22 mice) and one case of endometrial hyperplasia (1/28) (**Fig. 3B; Fig. S11-S12; Supplementary Tables 4 and 5**). Similar to *Pten*^+/-^ mice, *Pten*^+/R173C^ mice had a high incidence of pheochromocytoma (**Fig. 3B, Fig. S11-S12; Supplementary Tables 4 and 5**).

One out of 28 *Pten*^+/R173C^ mice displayed a single focus of endometrial hyperplasia, which correlated with loss of immunostaining for PTEN and increased AKT P-S473 (**Fig. 3C**). Interestingly, loss of PTEN immunoreactivity was also observed in microscopically *normal* endometrial glands from 5-month-old and 10-month-old *Pten*^+/R173C^ mice, but this was not accompanied by a concomitant increase in AKT P-S473 immunoreactivity in these regions (**Fig. 3C**). In the absence of reagents which allow us to specifically stain for WT or mutant PTEN protein and, based on the low expression level of endogenous PTEN-R173C protein when expressed alone (**Fig. S7**), it is most likely that expression of the wild-type *Pten* allele is lost in these morphologically normal glands. We speculate that, while the very low expression level of PTEN-R173C from the endogenous allele may not be detectable by IHC, it is still sufficient to partially downregulate PI3K signalling and AKT P-S473 and thereby prevent the development of endometrial tumours. Although this observation is based on a single mouse, we speculate that the hyperplastic gland found in the endometrium of this *Pten^+/R173C^* mouse may have resulted either from loss-of-expression of the remaining *Pten^R173C^* allele, or from acquisition of PI3K pathway-activating mutations. These events likely led to PI3K pathway hyperactivation, ultimately contributing to the development of endometrial hyperplasia. This hypothesis is supported by the fact that somatic cancers with the PTEN-R173 mutation, along with mutation of the wild-type allele of *PTEN*, also contain mutations in other cancer drivers such as *PIK3CA* (**Fig. 3D**).

### Prominent but delayed lymphoid hyperplasia in *Pten*^+/R173C^ mice compared to *Pten*^+/-^ mice

Previous studies have reported a range of lymphoid and immune phenotypes in PHTS patients including peripheral lymphoid hyperplasia, enlargement of tonsils and adenoids, hypogammaglobulinemia, recurrent respiratory tract infection and autoimmune disorders.^59,60^. Some of these phenotypes are found in a few PHTS patients with *PTEN-*R173 variants and other nuclear-excluded variants (**Table 1** and **Supplementary Tables 1&2).**

In *Pten* mutant mice, lymphoid hyperplasia leads to enlarged cervical, brachial, inguinal and axillary lymph nodes^25,51^. While this does not cause discomfort to the mice, in our study it was a main reason for sacrificing *Pten*^+/-^ mice (75% males and 80% females) and many *Pten*^+/R173C^ mice (21% males and 50% females), due to animal welfare regulations on the allowed tissue overgrowth size (**Supplementary Table 3**).

Histological analysis revealed a similar severity grade of hyperplasia (assessed by semi-quantitative histopathological scoring) in female *Pten*^+/R173C^ and *Pten*^+/-^ mice but of significantly lower grade in *Pten*^+/R173C^ males than *Pten*^+/-^ males (**Fig. 3E**). Lymphoid hyperplasia has also been reported in heterozygous *Pten^+/m3m4^*mice^32^.

Flow cytometry analysis of lymph nodes of 5-month-old female *Pten*^+/-^ and 10-month-old female *Pten*^+/R173C^ mice, the age at which they have visibly enlarged lymph nodes, compared to littermate *Pten*^+/+^ mice, revealed a significant increase in the proportion of B-cells and a corresponding decrease in T-cell proportions, a phenotype also reported in *Pten^+/m3m4^*mice (**Fig. 3F-G**).

Although we did not assess dsDNA damage repair in the lymphoid tissues of *Pten^+/R173C^*mice, the lymphoid phenotypes correlate with loss of nuclear PTEN as this phenotype was also seen in *Pten^+/m3m4^* mice^32^.

### Macrocephaly, increased neuronal complexity and white matter disturbances in *Pten*^+/R173C^ mice

PHTS patients, including the majority of patients with PTEN-R173 mutations (**Supplementary Table 1**), commonly present with macrocephaly, DD and ASD. We therefore explored neurological and behavioural phenotypes in *Pten*^+/R173C^ mice.

We examined the brains and neurons from *Pten*^+/+^, *Pten*^+/R173C^ and *Pten*^+/-^ mice and, where possible, conditional knock-out mice that selectively lack PTEN expression in cortical pyramidal neurons (referred to here as cKO)^61,62^. These cKO mice were further crossed with *Pten*^+/R173C^ mice to obtain conditional knock-in mice (referred to here as cKI) which express PTEN-R173C in cortical pyramidal neurons without a copy of the PTEN-WT allele and are overall hemizygous for the *Pten*^R173C^ allele (*Pten*^-/R173C^). The observed phenotypes are summarized in **Table 2**, which also shows a comparison to phenotypes published for *Pten^m3m4/m3m4^* mice. The latter model germline expression of non-naturally occurring PTEN mutations that leave PTEN PIP_3_ phosphatase activity intact but lead to its nuclear exclusion, and whose neuronal phenotypes have been extensively characterised^30–32^.

**Table 2.** Neuronal and behavioural phenotypes in mice with *Pten* mutations: Table shows data from *Pten^+/R173C^*, *Pten^+/-^*, and conditional *Pten^-/-^* mice from this study, and *Pten^+/m3m4^ and* Pten^m3m4/m3m4^ mice as reported in ^30,31^. ND; not determined. The behavioural assays (sociability, motor coordination and locomotion) are relative to wild type littermate mice.

| Phenotypes | Mouse genotype |  |  |  |  |
| --- | --- | --- | --- | --- | --- |
|  | *Pten*<sup>+R173C</sup> | *Pten*<sup>+/-</sup> | *Pten*<sup>-/-</sup> | *Pten*<sup>+/*m3m4*</sup> | *Pten*<sup>*m3m4/m3m4*</sup> |
| Increased brain size | Yes | Yes | ND | Yes | Yes |
| Bigger corpus callosum | Yes | Yes | ND | ND | ND |
| Increased numbers of oligodendroglia cells | Yes | No | ND | No | Yes |
| Astrogliosis | No | No | ND | No | Yes |
| Increased cell body size | Yes | No | Yes | No | Yes |
| Increased neuronal complexity | Yes | No | Yes | No | No |
| Enlarged axons | Yes | No | ND | ND | ND |
| thinner myelin | Yes | Yes | ND | ND | ND |
| Sociability | No significant difference | ND | ND | No significant difference | Too social with revised criteria |
| Motor coordination | ND | ND | ND | ND | reduced |
| Locomotion | reduced | ND | ND | ND | ND |

We first assessed the levels of PTEN protein and downstream pathways in protein lysates from cortical neurons. In the absence of a PTEN-WT allele, the level of PTEN protein in cKI was minimal and similar to that seen in protein extracts of cKO mice (**Fig. S13A-B**). However, the level of AKT P-S473 in cKI brains was significantly lower compared to the cKO mice (**Fig. S13A and C**). These observations mirror our data in MEFs (**Fig. S7**), indicating that PTEN-R173C is also highly active in the brain despite its low level of expression.

At 3 months of age, *Pten*^+/R173C^ and *Pten*^+/-^ mice of both sexes had enlarged brains with increased mass compared to *Pten*^+/+^ littermates (**Fig. 4A-B, left panel**). Whole-body mass was also increased in both *Pten* mutant genotypes (**Fig. 4B, middle panel**), however, the brain was disproportionally enlarged (**Fig. 4B, left panel**), indicating a macrocephalic phenotype. Also, at 3- and 6-months of age, and independent of the genetic background, *Pten*^+/R173C^ and *Pten*^+/-^ mice showed a similar increase in brain mass (**Fig. S14**). Given the apparently normal PI3K/AKT regulatory capacity of PTEN-R173C, and the neurological phenotypes and macrocephaly previously observed in nuclear-excluded PTEN in heterozygous *Pten^+/m3m4^* and homozygous *Pten^m3m4/m3m4^* mice^30,31^, these observations allow us to postulate that the nuclear function of PTEN is key to the development of macrocephaly.

**Figure 4.**
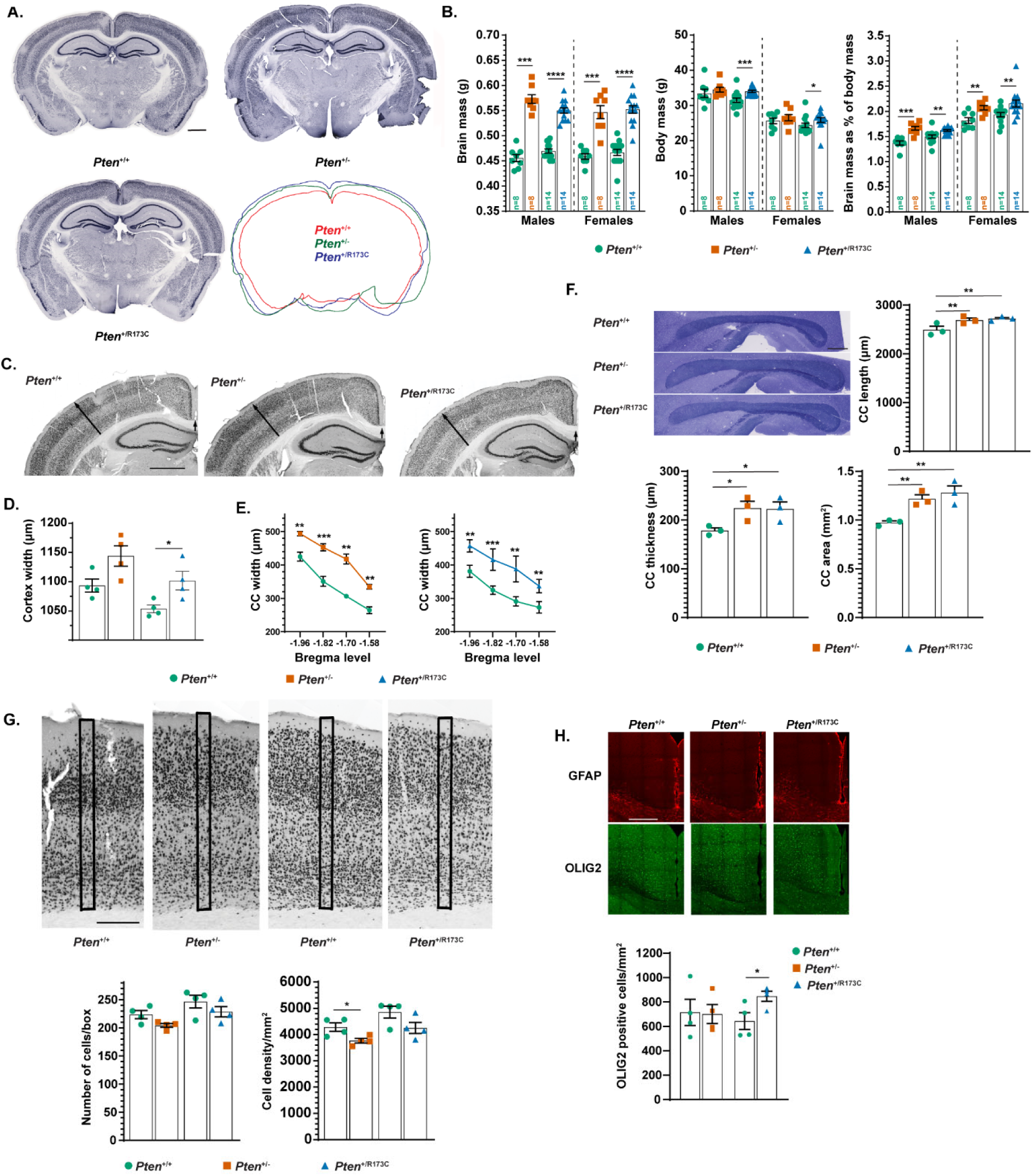
Enlarged brain phenotype with associated increases in grey and white matter areas in *Pten* mutant mice. (**A**) Representative NeuN immunostaining of coronal sections from *Pten*^+/+^ and mutant animals at 3 months. The overlay of the contours of the depicted slices is also shown. Scale bar: 1000 µm. (**B**) Brain mass, body mass and brain mass as a % of body mass for male and female *Pten*^+/-^, *Pten*^+/R173C^ and WT littermate controls at 3 months. Data show mean ± SEM, statistical analysis performed using Two-tailed Mann-Whitney U test. (**C**) Representative NeuN immunostaining from coronal sections of WT and mutant animals at 3 months. Scale bar: 1000 µm. (**D**) Quantification of cortical thickness in the primary somatosensory cortex barrel field (area indicated with long arrows in C). n = 4 mice of each sex and genotype. Data show mean ± SEM, statistical analysis performed using Two-tailed Unpaired Student’s t-test. (**E**) Quantification of the thickness of the corpus callosum at the indicated Bregma levels (area indicated with short arrows in C). n=4 mice of each sex and genotype. Data show mean ± SEM, statistical analysis performed using Two-way RM ANOVA with uncorrected Fisher’s LSD test. (**F**) Representative sagittal sections from control and mutant mouse brains stained with Toluidine Blue and quantification of the corpus callosum length, thickness, and area. Scale bar: 500 µm. n=3 mice per genotype. Data show mean ± SEM, statistical analysis performed using One-way RM ANOVA with uncorrected Fisher’s LSD test. (**G**) Top: Representative images of coronal sections from the somatosensory cortex of *Pten*^+/+^ and mutant animals at 3 months stained for NeuN. Scale bar: 250 µm. Bottom: Quantification of the number and density of neurons in the indicated boxed areas in (G). n=4 mice of each sex each genotype. Data show mean ± SEM; statistical analysis performed using Two-tailed Unpaired Student’s t-test. (**H**) Top: Representative images of sections from *Pten*^+/+^ and *Pten* mutant animals at 3 months immunostained with OLIG2 and GFAP, Scale bar: 100 µm, and bottom: quantification of OLIG2 positive cells in the motor cortex. n=4 mice per genotype. Data show mean ± SEM and statistical analysis performed using Two-tailed Unpaired Student’s t-test. p-values for all statistical tests: *, p<0.05; **, p<0.01; ***, p<0.001; ****, p<0.0001. Mice used were on a mixed C57BL/6J x Sv129 background.

We next quantified several brain parameters to characterize the macrocephalic phenotype in more detail. *Pten*^+/R173C^ mice had increased cortical thickness as measured at the level of the somatosensory barrel field (**Fig. 4C-D**). A significant increase was also observed in the thickness of the corpus callosum at four different Bregma levels (**Fig. 4E**) in both *Pten*^+/R173C^ and *Pten*^+/-^ mice. Consistent with macrocephaly, a lengthwise enlargement of the corpus callosum as well as an increase in the total area of this white matter tract at Bregma level were also observed (**Fig. 4F**). The increased thickness of the cortex was not caused by increased NEUN+ neuronal cell numbers or neuronal densities (**Fig. 4G**). In contrast, immunohistochemistry for OLIG2 revealed a small but significant increase in OLIG2-expressing oligodendrocyte lineage cell numbers in the cortex of *Pten*^+/R173C^ mice (**Fig. 4H**). Interestingly, this increase was not seen in our *Pten^+/-^* mice and has not been reported in *Pten^+/m3m4^*mice but was observed in *Pten^m3m4/m3m4^* mice. Astrocyte overabundance and astrogliosis, as detected by GFAP immunostaining, were not observed (**Fig. 4H**). Interestingly, astrogliosis has been reported in *Pten^m3m4/m3m4^* mice, suggesting that the remaining wild type *Pten* allele protects the brain from this immunoinflammatory phenotype.

To gain further insight into the causes of the macrocephaly, we quantified neuronal and glial parameters *in vitro*. Morphologies of cortical pyramidal neurons *in vitro* were assessed following transfection with a GFP expression vector to allow visualisation of the cells (**Fig. 5A**). *Pten*^+/R173C^ neurons had enlarged cell body size and increased process complexity compared to WT controls (**Fig. 5B,C**). While cKO neurons demonstrated a more severe phenotype, this phenotype was not observed in *Pten*^+/-^ neurons and not reported in *Pten^+/m3m4^* mice but was observed in *Pten^m3m4/m3m4^* mice (**Fig. 5B,C**). To determine whether glial and/or axonal defects contribute to the enlargement of the white matter, we examined the corpus callosum of *Pten*^+/+^, *Pten*^+/R173C^ and *Pten*^+/-^ mice using transmission electron microscopy (**Fig. 5D**). We found a small increase in average axonal diameter in *Pten*^+/R173C^ mice, with a reduction in the number of small diameter axons and a gain of larger diameter axons (**Fig. 5E**). This phenotype was not observed in *Pten^+/-^* mice. Quantification of the G-ratio in the corpus callosum showed an increase in average myelin thickness in both *Pten*^+/R173C^ and *Pten*^+/-^ mice (**Fig. 5F**). However, scatter plot analysis of the g-ratio as a function of axonal diameter did not show significant changes in either *Pten*^+/R173C^ and *Pten*^+/-^ mice compared to controls (**Fig. 5G**).

**Figure 5.**
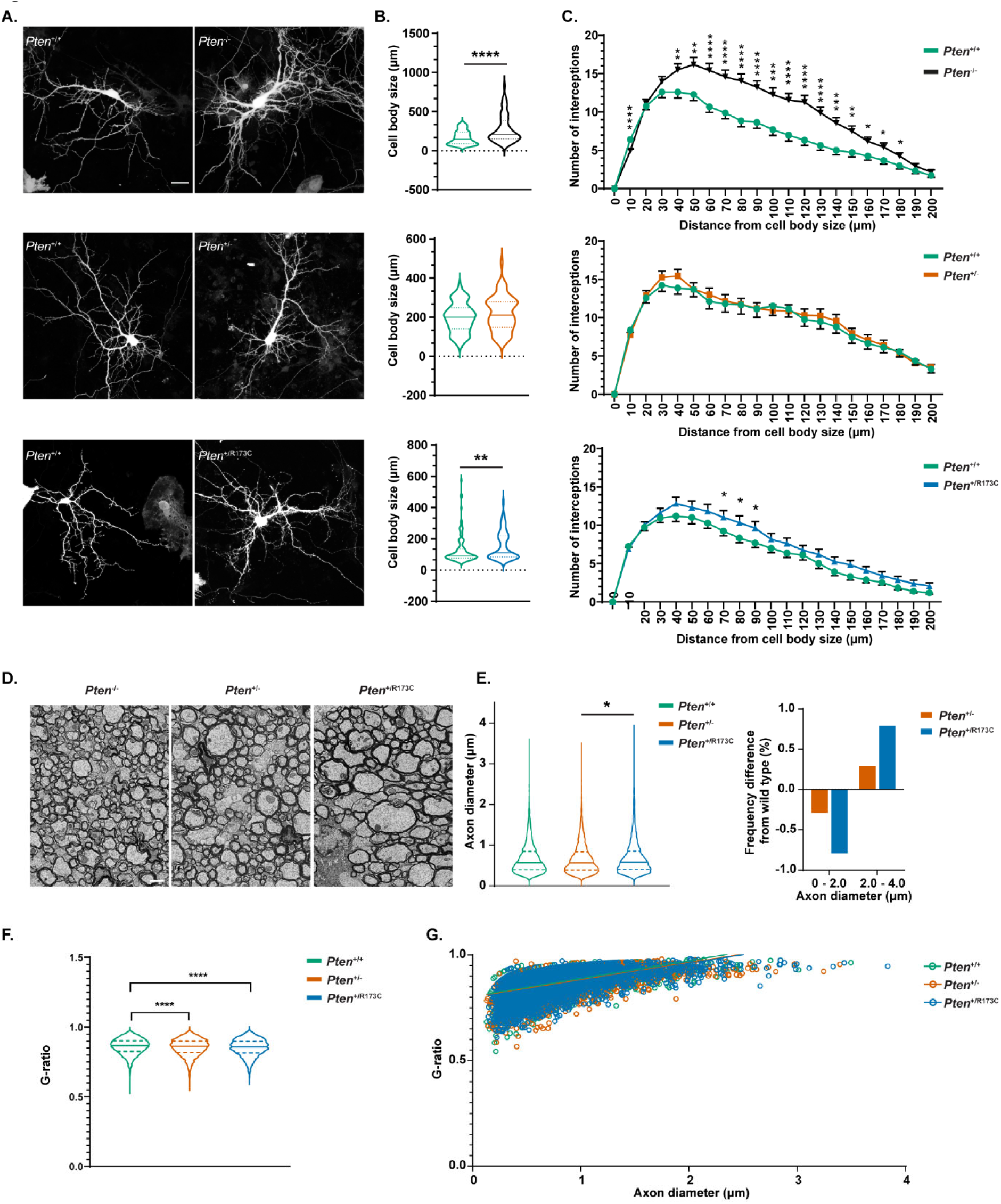
Neuronal and glial deficits in *Pten*^+/R173C^ and *Pten*^+/-^ mice. (**A**) Representative images from GFP-transfected cortico-hippocampal neurons prepared from the indicated *Pten* mutants and their respective *Pten*^+/+^ littermate controls at DIV12/13. Neurons were stained for GFP. Scale bar: 20 µm. (**B**) Quantification of cell body size of GFP-transfected cultured cortico-hippocampal neurons shown in (A). from n>60 cells for each genotype from 3-6 independent experiments. Violin plots show mean ± SEM, statistical analysis performed using Two-tailed Mann-Whitney U test. (**C**) Sholl analysis of GFP-transfected cortico-hippocampal neurons shown in (A) from n=63–122 cells from 3–5 independent experiments. Data show mean ± SEM, statistical analysis performed using Two-way RM ANOVA with uncorrected Fisher’s LSD test. (**D**) Representative electron micrographs of the corpus callosum in cross-sections from 3-month-old *Pten*^+/+^, *Pten*^+/-^ and *Pten*^+/R173C^ mice. Scale bar: 1 µm. (**E**) Quantification of axonal dimeters and their frequency difference in mutant mice compared to *Pten*^+/+^ littermate controls. >4500 axons measured in each of three mice per genotype. Violin plots indicate mean ± SEM, statistical analysis performed using Two-tailed Mann-Whitney U test. (**F**) Quantification of g-ratios for corpus callosum axons. >4500 axons measured in each of three mice per genotype. Statistical analysis performed using Two-tailed Mann-Whitney U test. (**G**) Scatter plot of g-ratios as a function of axon diameter in 3-month-old *Pten*^+/+^, *Pten*^+/-^ and *Pten*^+/R173C^ mice. >4500 axons measured in each of three mice per genotype. p-values for all statistical tests: *, p< 0.05; **, p<0.01; ***, p<0.001; ****, p<0.0001. Mice used in (A-C) were on a C57BL/6J background, and in (D-G) were on a mixed C57BL/6J x Sv129 background.

Altogether, our experiments show that the macrocephalic *Pten*^+/R173C^ phenotype is caused by alterations in multiple parameters, including increases in neuronal cell body size, axonal thickness, axonal complexity and oligodendrocyte lineage cell numbers.

*Pten^+/R173C^* mice have a similar degree of macrocephaly as *Pten^+/m3m4^* heterozygous mice and have multiple neuronal phenotypes overlapping with *Pten^m3m4/m3m4^* homozygous mice and *Pten^+/-^* mice (**Table 2**). Importantly, the apparently intact or even increased PI3K/AKT regulatory capacity of PTEN-R173C, and the phenotypic overlap with *Pten^m3m4^* mice, suggests that the nuclear function of PTEN is key to the development of macrocephaly and several neuronal phenotypes.

Defective dsDNA damage repair is increasingly being associated with macrocephaly and ASD with studies showing increased dsDNA breaks in cells isolated from patients with these phenotypes^63,64^. To explore the possible involvement of impaired dsDNA damage repair in neurons in the macrocephalic phenotype, we treated *Pten^+/-^* and *Pten^+/R173C^* mice with IR and examined the levels of γH2AX in the brain 5 h post-IR treatment. While there was an increase in the γH2AX levels in protein extracts from the brains of both *Pten^+/-^* and *Pten^+/R173C^* mice, compared to that seen in littermate wild-type mice, the difference observed was not statistically significant (**Fig. S15**). This might be due to the IR dosage and/or time of analysis, and warrants further investigation.

### *Pten*^+/R173C^ mice display mild behavioural aberrations

ASD is common in PHTS ^4^. Behavioural abnormalities are found in mice with germline expression of heterozygous loss-of-function of *Pten*^65^ or homozygous expression of the nuclear-excluded *Pten*^m3m4^ mutations^31^. To examine such type of aberrations in *Pten*^+/R173C^ mice, we used a range of tests associated with ASD-like mouse behaviours.

In Crawley’s sociability test, *Pten*^+/R173C^ mice were indistinguishable from *Pten*^+/+^ littermate controls, indicating normal social interactions (**Fig. S16A-D**). In the open field test, male mice were not significantly different from their *Pten*^+/+^ littermates with regards to distance travelled in the open field arena and time spent moving during the test (**Fig. S17B,C**). However, female *Pten*^+/R173C^ mice travelled less and spent less time moving than WT controls (**Fig. S17F,G**), suggesting decreased exploratory behaviour and/or decreased locomotion. This was not caused by increased anxiety since female (and male) mice spent equal length of time exploring the different compartments of the open field test as their WT controls (**Fig. S17D,H**), with defecation during the test being comparable between the two groups (**Fig. S17E,I**). Reduced exploratory locomotion in females was also observed in a hole board test (**Fig. S18A**). Changes in repetitive behaviours, as assayed in a marble burying test were absent in both males and females (**Fig. S18B**).

## DISCUSSION

Since the discovery of PTEN as a tumour suppressor in somatic cancer, multiple genetic mouse models targeting *Pten* have been generated, highlighting the importance of PIP_3_ phosphatase activity of PTEN in tumour suppression, embryonic development, cellular homeostasis, neuronal development and the development of autism spectrum traits^24^.

PTEN mouse models have also improved our understanding of PHTS, a rare genetic condition caused due to heterozygous germline *PTEN* mutations. These include *Pten^+/-^* mice which represent the heterozygous genetic constellation of *PTEN* in PHTS and on a mixed C57BL6xSv129 background develop multiple phenotypes overlapping with the clinical presentation of PHTS, including brain and lymphoid overgrowth and the development of specific cancers^26^. While the PIP_3_ phosphatase activity of PTEN has been the focus of research over the last three decades, there is an increasing body of evidence for other PTEN functions in disease pathogenesis, in particular its nuclear function^14^. Recently, the *Pten^m3m4^* mouse model with germline expression of nuclear-excluded, engineered non-naturally-occurring *Pten* mutations, has indicated a role for nuclear PTEN defects in driving neuronal and immune phenotypes in PHTS^31,32^.

In this study, we developed a new mouse model featuring the pathogenic PTEN-R173C variant commonly found in PHTS and somatic cancer. We report that this PTEN variant retains its ability to regulate PI3K/AKT signalling but is impaired in dsDNA damage repair through HR, likely due to nuclear exclusion. Contrary to previous reports^27–29^, we found that PTEN-R173 mutant proteins (R173C and R173H) were catalytically more active PIP_3_ lipid phosphatases than PTEN-WT, both in cells and tissues. However, these mutants displayed reduced protein stability, possibly resulting from a more open protein configuration that is more susceptible to ubiquitination-mediated degradation. Upon heterozygous germline expression in *Pten*^+/R173C^ mice, this biochemical feature results in PTEN protein expression levels comparable to those in *Pten*^+/-^ mice but an overall lipid phosphatase activity similar to wild-type mice, as demonstrated by effective downregulation of insulin-stimulated PI3K/AKT signalling.

*Pten*^+/-^ mice with heterozygous PTEN loss are predisposed to a range of tumours similar to those observed in PHTS patients^26^. To assess the importance of nuclear functions of PTEN in tumourigenesis, we aged *Pten*^+/R173C^ mice and found only a modest predisposition to solid tumour development compared to *Pten*^+/-^ mice. This is reminiscent of mouse models of genes involved in dsDNA damage repair via HR such as *Brca*, *Rad51* and *Palb*^66–71^. Indeed, while homozygous inactivation of these genes frequently leads to embryonic lethality, as also observed in *Pten*^R173C/R173C^ mice, their heterozygous inactivation only leads to a very low to absent tumour burden, most likely because the lifespan of mice is insufficiently long to accumulate the additional mutations required for tumourigenesis^72^. That additional genetic defects may be required to allow phenotypic output of PTEN-R173 defects is indicated by the observation that somatic cancers with PTEN-R173 mutation have a range of co-occurring mutations, including but not limited to oncogenic mutation in *PIK3CA*or loss of second allele of *PTEN* (**Fig. 3D**). Also in mouse models, combination of *Pten* mutations with other oncogenic drivers such as mutant *Pik3ca* and *ErbB2* have accelerated development of mammary tumours^73,74^. Combining the *Pten-R173C* mutation in mouse models carrying additional cancer drivers (such as *PIK3CA* mutation) or defects in DNA repair genes (such as *Brca* or *Rad51*) could provide evidence for this speculation, but such experiments are outside the scope of the current study.

Our data further show that, in a germline configuration, deficient nuclear localisation of PTEN, and consequently its nuclear function, correlates with brain and lymphoid overgrowth. While our study does not establish a direct correlation between these overgrowth phenotypes in our mice and dsDNA damage repair, it is possible that such a link exists. Indeed, one study reported increased IR-induced DNA strand breaks in lymphocytes from autistic children compared to non-autistic controls supporting such a functional link^64^. On the other hand, lymphoid cells isolated from PHTS patients with PTEN-R173C mutation, who all present with ASD, did not exhibit increased DNA damage following IR treatment compared to wild-type controls^75^, suggesting that nuclear functions of PTEN other than dsDNA repair may underlie the observed lymphoid overgrowth and macrocephaly in *Pten^+/^*^R173C^ mice. This is further supported by the absence of lymphoid overgrowth and macrocephaly in patients and mouse models with germline mutations in other genes regulating dsDNA damage repair by HR, such as the *Brca*, *Rad51* and *PalB2* genes.

We find that the macrocephalic *Pten*^+/R173C^ phenotype correlates with alterations in multiple parameters, including increases in neuronal cell body size, axonal thickness, axonal complexity and oligodendrocyte lineage cell numbers. While this brain overgrowth phenotype in *Pten*^+/R173C^ mice is very clear, its impact on animal behaviour is less prominent. It has to be said that while mice with heterozygous germline expression of PTEN accurately reflect the genetic makeup of patients with PHTS, the behavioural abnormalities observed in heterozygous mutant mice are often subtle and highly variable^65^. This necessitates large mouse numbers to achieve quantifiable significant differences between genotypes. Consequently, the field of behavioural analysis has mostly relied on the use homozygous mouse models. This approach is particularly true for PHTS, with numerous studies conducted on homozygous brain-specific PTEN knockout models^76,77^ and homozygous *Pten^m3m4^*models^31^. Similar homozygous models are also utilized in behavioural research on other rare diseases such as Tuberous Sclerosis^78,79^. Although these models do not share the exact genetic composition of patients, they serve as platforms for behavioural and neurological studies and testing therapeutics.

Our study suggests that the status of nuclear function of PTEN alongside its PIP_3_ phosphatase activity, allows refinement of genotype/phenotype analysis in PHTS. Based on our findings and assessment of patient phenotypes (**Table 1** and **Supplementary Tables 1 and 2**), our data indicate that individuals carrying *PTEN* variants such as *PTEN^R173C^* that retain their PIP_3_ phosphatase function but are nuclear-excluded tend to follow a characteristic progression of known PHTS disease phenotypes: early-signs such as neurological abnormalities – including macrocephaly and global developmental delay - alongside cutaneous features which together lead to a PHTS diagnosis during childhood. As they enter early adulthood, they may be prone to developing vascular abnormalities and benign tumours in the thyroid and GI tract, with an increased likelihood of developing malignant cancers later in life.

Our study also highlights the possibility of new therapeutic treatments for cancer in PHTS patients. A previous study has shown that glioma cells expressing PTEN-Y204F, a mutation that leads to lack of PTEN nuclear localisation and functions such as dsDNA damage repair are more sensitive to radiation therapy^35^. It is therefore tempting to speculate that radiation therapy and dsDNA damage repair antagonist drugs, such as *PARP* inhibitors, might be additional therapeutic options beyond PI3K pathway inhibitors for cancer in PHTS patients with R173 mutant-like PTEN variants.

## Methods

### Plasmids

DNA constructs are detailed in **Supplementary Table 8**. Point mutations were introduced in PTEN WT cDNA in different vectors using primers listed in **Supplementary Table 9** and KOD hot start DNA polymerase (Sigma-Aldrich) using the manufacturer’s instructions.

### Mice

All mice were maintained at University College London in accordance with the UK Animals (Scientific Procedures) Act 1986 and following UK Home Office guidance. All procedures were authorized by a UK Home Office Project Licence subject to local ethical review. *Pten*^+/-^ mice ^25^, *Pten*^flox/flox^ mice ^62^, *CAG CreER^T2^* mice ^80^ and *Emx1-Cre* mice ^61^ have been described elsewhere. *Pten*^+/R173C^ mice were generated by Taconic Biosciences using a CRISPR/Cas9 approach. gRNA was designed to contain the R173C mutation (c.517C>T) and a silent mutation containing the *Eco*NI restriction site upstream of the R173 site (to allow discrimination of wild-type and knock-in allele following PCR). **Fig. S6** shows the gRNA sequence and the targeting strategy. gRNA was microinjected into C57BL/6J mouse zygotes along with recombinant Cas9. The zygotes were then transferred to pseudo-pregnant mothers. The litters born (founder animals) were sequenced to check for the incorporation of the R173C mutation and percentage of mosaicism. They were then used to as a breeder to generate F1 animals. F1 males positive for the R173C mutation and negative for any off-target mutations were used to generate further animals by IVF using C57BL/6J females. All mouse lines were maintained on a C57BL/6J background. Sv129 mice were purchased from Charles River or Envigo. *Pten*^+/-^ and *Pten*^+/R173C^ mice were crossed with Sv129 mice to obtain mixed background F1 mice that were used for some experiments as described. For neuronal studies, conditional knock-out mice that selectively lack PTEN expression in cortical pyramidal neurons *were generated by crossing Pten*^flox/flox^ mice with *Emx1-Cre* mice to give *Emx1-Cre*;*Pten*^fl/fl^; referred to here as cKO. The cKO mice were crossed with *Pten*^+/R173C^ mice to obtain *Emx1-Cre*;*Pten*^fl/R173C^,referred to here as cKI, which express one only copy of *Pten*-R173C in the cortical pyramidal neurons.

For genotyping, mouse tissue samples (from ear biopsy or from the embryonic yolk sac) were lysed in 25 mM NaOH/0.2 mM disodium EDTA for 1 h at 95°C. The samples were neutralized by adding an equal volume of a buffer containing 40 mM Tris-HCl, pH 4.5. Two µl of the DNA extract was used for PCR using and Titanium polymerase PCR kit (Takara) and primers listed in **Supplementary Table 10**. For genotyping of *Pten*^+/R173C^ mice, an additional restriction digestion step was performed on the PCR products using *Eco*NI (New England Biolabs).

### Mouse survival studies

For sample size calculations for long term survival study, we use ClinCalc (www.clincalc.com). The mean survival age of *Pten*^+/-^ on a mixed C57BL6J x Sv129 background has been reported to be 12±1.8 months ^51^. The sample size required for *Pten*^+/R173C^ mice needed to see a 15% improvement in mean survival age compared to *Pten*^+/-^ mice was calculated as 16 (probability of Type-I error was set to 0.05 and Power was set to 80%). We used at least 16 mice for each genotype and each sex. At least 10 Littermate wild type mice were used for both *Pten*^+/-^ and *Pten*^+/R173C^ mice. For survival analysis, mice were weighed twice a week. They were monitored and euthanized upon showing a more than 20% reduction in body weight, any signs of ill health such as hunched appearance, laboured breathing, sluggish behaviour, or palpable masses with a total surface area of 1.4 cm^2^ surface area. Animals showing no clinical signs were euthanized at the end of the study (600 days for females and 730 days for males).

### Cell lines

U-87 MG (U87), HEK-293T and Phoenix cells were purchased from ATCC, Lenti-X™ 293T cells were purchased from Clontech. Human PTEN knockout HeLa cell line and control HeLa cells was obtained from Abcam (ab255419 and ab255928); PTEN knockout was validated using western blotting (**Fig. S4C**). All cell lines were maintained in DMEM (Sigma-Aldrich), supplemented with 10% Filtered Bovine Serum (FBS) (Pan-Biotech) and Pen-Strep (Sigma-Aldrich) using standard cell culture methods. **Preparation of MEFs.** For preparation of E13.5 MEFs, timed matings were set up between *Pten*^+/-^ mice and *Pten*^+/R173C^ mice, *Pten*^flox/+^*CreER^T2^* and *Pten*^+/-^ or *Pten*^+/R173C^ mice. Pregnant females were euthanized on day 13.5 and embryos harvested from the uterus. Following removal of the head and visceral organs, the rest of the embryo was minced in 2x Trypsin/EDTA using a sterile scalpel and incubated at 37°C for 40 min. The cell suspension was plated on 10 cm dishes in DMEM (Sigma-Aldrich) medium supplemented with 10% FBS (Pan-Biotech) and Pen-Strep (Sigma-Aldrich), and cells maintained using standard cell culture procedures. For preparation of E9.5 MEFs, timed intercrosses were set up between *Pten*^+/R173C^ mice. Pregnant females were euthanized on day 9.5, embryos removed from the uterus and gently dissociated by pipetting up and down using a p1000 pipette and 2 ml of media containing DMEM (Sigma-Aldrich), supplemented with 10% FBS (Pan-Biotech) and Pen-Strep (Sigma-Aldrich). The cell suspension was plated in 48-well plates and cells maintained using standard cell culture procedures. To induce Cre-mediated recombination of the floxed *Pten* allele in *Pten*^flox/+^*CreER^T2^*MEFs, 4-hydroxy tamoxifen (Sigma Aldrich) was added to the cells at a final concentration of 1 µM and refreshed every 24 h for 3 days.

### Primary neuron culture

Primary cortico-hippocampal cultures were prepared from P1 or P2 pups after genotyping. Cortex and hippocampus were dissected, digested with Trypsin, dissociated using glass pipettes and 40,000 cells were plated per 12 mm coverslip. Neurons were cultured in serum-free medium (neurobasal, NBA) medium supplemented with B27 supplement, L-glutamine, penicillin/streptomycin and 10 mM HEPES pH 7.4. Cells were cultured at 5% CO_2_ at 37°C with replacement of half of the medium every 2-3 days. Cells were transfected on DIV6 with a GFP expression vector using Lipofectamine 3000 according to the manufacturer’s instructions. On DIV 12/13 cells were fixed with 4% PFA/4% sucrose in PBS for 10 min at room temperature (RT) and washed 3x in PBS. Cells were blocked and permeabilized in immunofluorescence (IF) buffer (PBS supplemented with 0.3% Triton X and 3% BSA) for 20 min and incubated with primary antibodies to NeuN (Millipore MAB377 and GFP, Abcam 1:1000) for 2 h at RT in IF buffer in a humidity chamber. After 3 washes with PBS, cells were incubated with cross-absorbed Alexa Fluor labelled secondary antibodies (Invitrogen) in IF buffer for 1 h at RT in a humidity chamber in the dark, washed 3x with PBS and mounted with Immunomount. Neurons were imaged using a LSM 880 microscope (Leica) acquiring Z-stacks at 40 x magnification. Dendritic complexity was determined using the Sholl macro in FIJI. Interceptions were analysed in 10 µm radii within a 200 µm distance from the centre of the soma. To measure cell body size in FIJI, stacks were processed to z-projections, ROIS were drawn manually around the cell bodies and the ROI area was measured in FIJI.

### PTEN activity assays

For purification of bacterially-expressed GST-tagged PTEN proteins, pGEX6P1 vectors with cDNA for WT or mutant PTEN were introduced in *E. coli* and recombinant PTEN was purified by glutathione-affinity chromatography followed by cleavage of the GST tag as described ^18^. PIP_3_ assays using purified recombinant PTEN were performed with diC8 PIP_3_ (Cell Signals Inc) and malachite green phosphate assay kit (Sigma Aldrich) and have been described elsewhere ^81^. Purification of recombinant PTEN protein from insect cells was as described in Ref.^82^.

### Lentiviruses and retroviruses

Lentiviruses for PTEN and PIP_3_ biosensor were generated by co-transfecting Lenti-X™ 293T cells or HEK-293T cells with plasmids expressing the cDNA of interest along with packaging vectors using TransIT-LT1 following the manufacturer’s instructions. For preparation of retroviruses, Phoenix cells were transfected with p53 shRNA expressing vector ^83^ using TransIT-LT1 following the manufacturer’s protocol. 24 h post transfection, sodium butyrate was added to the cells to a final concentration of 12.5 mM for 6 h, followed by washing the cells with warm PBS and addition of fresh media. The supernatant containing lentiviral particles was collected after 20 h, passed through a 0.45 µm filter and stored at -80°C. Target-cells for transductions were plated at 40-50% confluency. Once the cells had attached, lentiviral/retroviral particles were added to the cells along with polybrene (Sigma-Aldrich) at 20 µg/µl. The media was changed 24 h post transduction.

### Immunoblotting analysis

Protein extracts from cell cultures were prepared by scraping cells into lysis buffer containing 25 mM Tris-HCl pH 7.4, 150 mM NaCl, 1% Triton X-100, 10% glycerol, 1 mM EGTA, 1 mM EDTA, 5 mM sodium pyrophosphate, 10 mM β-glycerophosphate, 50 mM sodium fluoride, 1 mM sodium orthovanadate, 1 mM DTT and Protease inhibitor cocktail (Millipore). Mouse tissues harvested from mice upon euthanasia were homogenize using Lysing Matrix M tubes (MP Biomedicals) in double the volume (v/w) of lysis buffer (25 mM Tris-HCl pH 7.4, 150 mM NaCl, 1% Triton X-100, 0.1% SDS, 10% glycerol, 1 mM EGTA, 1 mM EDTA, 10 mM sodium pyrophosphate, 20 mM β-glycerophosphate, 100 mM sodium fluoride, 2 mM sodium orthovanadate, 1 mM DTT and protease inhibitors) on a FastPrep 24 homogeniser (MP Biomedicals) at 4 m/sec for 20 sec. Cell or tissue lysates were pre-cleared by centrifugation at 20,000xg for 10 min and protein extracts analysed by immunoblotting. 293T cells expressing Rluc-PTEN-YFP constructs were lysed in buffer containing 50 mM HEPES (pH 7.4), 250 mM NaCl, 2mM EDTA, 0.5% NP-40, 10% glycerol supplemented with protease and phosphatase inhibitors (Roche) and protein A/G PLUS-Agarose (30µl/IP) and GFP antibody (Roche, Cat #11814460001, Mouse, Dilution 1:200) was used to immunoprecipitate YFP-tagged PTEN protein which was used for immunoblotting as described below. Protein gel electrophoresis was conducted with 10 µg of total soluble protein per lane on NuPage Bis-Tris 4-12% gradient polyacrylamide gels (Thermofisher Scientific) following the manufacturer’s protocols. Proteins were transferred onto PVDF membrane (Millipore) and membranes blocked in 5% milk powder/TBST for 1 h at RT. Blocked membranes were incubated overnight with primary antibodies (**Supplementary Table 11**). Antibody complexes were detected by 1 h incubation at RT with HRP-conjugated secondary antibodies (GE healthcare). Blots were developed with Immobilon Forte Western HRP substrate (Millipore) and chemiluminescence imaged using an ImageQuant LAS4000 imaging system (GE healthcare).

### qRT-PCR assays

Total RNA was extracted from U87cells non-transduced or transduced with lentiviruses for PTEN-WT, PTEN-C124S, PTEN-R173C or PTEN-R173H, using RNAEasy kit (Qiagen) following the manufacturer’s instructions. 1 µg of RNA was converted to cDNA using iScript RT kit (BioRad). 50 ng of cDNA was used for QRT-PCR reactions using primers in **Supplementary Table 12**. The reactions were run on a QuantStudio™ 5 Real-Time PCR machine (Applied Biosystems). dCT was calculated by deducting the CT for GAPDH from CT of PTEN. ddCT was calculated by deducting the dCT of untransduced cells from the dCT of the sample of interest.

### Cycloheximide chase studies

U87 cells were transduced with lentiviruses for PTEN-WT, PTEN-C124S, PTEN-R173C or PTEN-D252G. 48 h post transduction, cells were treated with 200 µg/ml cycloheximide, lysed at different time points and protein extracts analysed for PTEN expression by immunoblotting.

### BRET assays

BRET assays using the Rluc-PTEN-YFP biosensor were performed as described in Refs ^44,84^. Briefly, 24 h post-transfection, cells were detached from 12-well plates, using trypsin-EDTA, and resuspended in 1 ml of complete media. The cells were subsequently distributed into poly-L-ornithine-coated (30 µg/ml) white 96-well optiplates (Perkin Elmer). The next day media was replaced with Opti-MEM (Gibco) and Coelenterazine substrate (Interchim) was added to a final concentration of 5 µM and incubated for 3 min at 25°C. BRET readings were then collected using a Multilabel Reader Mithras2 LB 943 (Berthold Technologies). Sequential integration of light output was measured for 1 sec using two filter settings (480+10nm for YFP and 540+20nm for Rluc). The BRET signal represents the ratio of the light emitted by YFP and the light emitted by Rluc (YFP/Rluc). The ratio values were corrected by subtracting background BRET signals obtained with Rluc-PTEN. MilliBRET (mBRET) values were calculated by multiplying these ratios by 1,000. Data are represented as specific mBRET values or ΔmBRET when compared to the signal obtained with WT PTEN set to 0.

### MTS assays

U87 cells were transduced with lentiviruses for GFP, PTEN-WT, PTEN-C124S, PTEN-R173C or PTEN-R173H. 48 h post transduction, 5000 cells/well were plated in 96-well plates. A different plate was prepared for each day. 20 µl of MTS assay reagent (Abcam) was added to the cells, incubated at 37°C for 1 h, followed by measurement of absorbance at 490 nm using a plate reader.

### PIP_3_ biosensor analysis and PTEN localization using fluorescence microscopy

For plasma membrane PIP_3_ quantification, U87-MG cells were transduced with lentiviral particles expressing the PIP_3_ biosensor on its own or with lentiviral particles for PTEN-WT or PTEN-R173C, were pretreated ± GDC-0941 (1 µM) for 1 h and stimulated ± insulin (100 nM) for 2 min. For PTEN localization studies, MEFs were treated ± GDC-0941 (1 µM) 1 h prior to fixation. For both plasma membrane PIP_3_ quantification and localization of untagged or endogenous PTEN, cells were fixed with 4% PFA for 20 min, washed in PBS, permeabilized with 0.1% Triton X-100 in PBS for 90 sec, washed 3 times with PBS and blocked with PBS containing 1 or 3% (w/v) BSA for 30 min. Primary antibodies to PTEN were diluted in 1 or 3% BSA block solution and incubated for 1 h at RT. Following 3 washes with PBS, secondary antibodies and Phalloidin were diluted in 1 or 3% BSA block solution and incubated for 45 min at RT. Cells were then washed 3 times and mounted using ProLong Gold antifade reagent containing DAPI (P36935, Life Technologies). To examine GFP-PTEN localization during different stages of the cell cycle, Lenti-X™ 293T cells were transfected with DNA for PTEN-WT-GFP or PTEN-R173C-GFP. 18 and 36 h post transfection, culture medium was changed to fresh media containing 2 mM thymidine (Sigma). 18 h post the second media change, the cells were released from double thymidine block by changing into fresh, complete media. Cells were fixed with 4% PFA for 15 min, washed in 3 times in PBS and permeabilized with 0.1% Triton X-100 in PBS for 5 min. Cells were then washed 3 times with PBS and once with water before mounting using ProLong Gold antifade reagent containing DAPI. Confocal microscopy was performed using a Zeiss LSM 880 microscope with AiryScan and a 63x PL APO objective and Zen Black acquisition software or Zeiss LSM900 microscope and a 63x or 40x PL APO objective and Zen Black acquisition software. Image processing was performed using Image J software (National Institutes of Health, Rockville, MD, United States) and was limited to alterations of brightness, subjected to the entire image. Quantification of plasma membrane PIP_3_ intensity was performed using Image J as described previously with minor modifications ^85^. The average fluorescence intensity was measured within a box of defined size (20 x 10 pixels) at three random regions of the plasma membrane and three random regions of the cytosol (box size 20 x 20 pixels). The plasma membrane and cytosol fluorescence intensity measurement were each averaged and the ratio of plasma membrane to cytosolic fluorescence intensity determined. The PTEN nuclear to cytoplasmic ratio was measured using Image J. First, the nuclear PTEN fluorescence intensity was measured, followed by the cytoplasmic PTEN fluorescence intensity. The ratio of nuclear to cytoplasmic PTEN fluorescence intensity was then calculated.

### Insulin tolerance test (ITT)

3-month-old male *Pten*^+/-^ and *Pten*^+/R173C^ mice and their WT littermates were starved for 6 h. The tip of the tail was punctured using a 26G needle and time-0 blood glucose levels were recorded using a standard glucometer. The mice were given an IP injection of insulin to a final concentration of 0.75 U/kg. Blood glucose readings were recorded at 15 min, 30 min, 45 min, 60 min, 90 min and 120 min post injection.

### Irradiation of mice

8-12-week-old mice were subjected to whole body irradiation of 7 Gy dose using the Small Animal Radiation Therapy Platform (SARRP). 5 h post irradiation, the mice were euthanised with a lethal injection of pentobarbital. Tissues of interest were harvested and snap frozen in liquid nitrogen for biochemical analysis.

### Flow cytometry

Mouse spleens and lymph nodes were mashed with a syringe plunger in RPMI containing 10% FBS and filtered through a 70 µm filter. Cell suspensions from spleen were treated with RBC lysis buffer (Biolegend). The cell suspensions were washed with PBS and stained with Fixable Viability Dye eFluor™ 780 (ThermoFisher scientific) in PBS for 40 min followed by a few PBS washes. For staining of surface antigens, antibodies were diluted in FACS staining buffer at 1:100 and added to the cells for 30 min followed by two washes with FACS buffer. Details on antibodies and reagents used is shown in **Supplementary Table 11**. The stained cells were analysed on BD FACSymphony™ analyser.

### Tissue processing for histological analysis and immunohistochemical analysis

Mouse tissues were fixed in 10% neutral buffered formalin for 24 h, followed by processing and embedding into paraffin blocks. The tissues were orientated with the biggest surface down, to obtain complete sections. 3 µm serial sections were cut and stained for Haematoxylin and Eosin, followed by immunohistochemistry staining on the Leica BOND Rxm. Tissue sections for immunohistochemistry were incubated for 40 min at 100°C in BOND epitope retrieval solution 2 (Leica, Germany), followed by blocking and staining using the BOND polymer refine detection kit (Leica, DS9800), primary antibody (**Supplementary Table 11**) and DAB enhancer (Leica, Germany).

### Histopathological analysis

Non-neoplastic lesions, e.g. splenic extramedullary haematopoiesis and lymph node hyperplasia, were graded using a standard semi-quantitative grading scheme from 0 to 5, where 0 = lesion not present and 5 = the maximum lesion size/extent ^86^. In general, where present, the presence of other proliferative lesions including hyperplasia and neoplasia was recorded for each individual organ, allowing the overall incidence to be established for each lesion in each group. Prostatic intraepithelial neoplasia (PIN) was graded as low or high grade according to the consensus suggested by Ittmann *et al.* ^87^. Low-grade PIN lesions are focal, with one to two layers of cells and mild nuclear atypia. High-grade lesions tend to be more extensive, filling the prostatic lumen but without stromal invasion; have two or more cell layers often forming papillary or cribriform patterns; increased nuclear atypia and mitoses.

### Perfusion and preparation of brain sections for DAB immunostaining and immunofluorescence

Mice were perfused with saline (0.9 % NaCl) followed by 4% PFA (w/v) in PBS through the left ventricle of the heart. Brains were removed and post-fixed overnight at 4°C in 4% PFA, cryoprotected in 20% sucrose in PBS for 24 h at 4°C and embedded in Tissue-Tek OCT compound (Sakura) in isopentane cooled on dry ice. Frozen brains were stored at -80°C until use. Brains were processed on a Cryostat (30 µm thickness) and sections were collected in PBS for immunohistochemistry. For DAB immunostaining, floating sections were incubated with 0.3 % H_2_O_2_ in PBS for 30 min while shaking at RT, and then blocked using blocking buffer (10% FBS in PBS supplemented with 0.1% Triton X-100) for 1 h at RT. After blocking, sections were incubated overnight at 4°C with primary antibodies diluted in blocking buffer. Primary antibodies were detected with biotinylated secondary antibodies (Donkey Anti-Mouse IgG Biotinylated Antibody, R&D systems) diluted 1:300 in blocking buffer for 1 h at RT. The signal was amplified using the Vectastain ABC Kit HRP kit (Vector, PK-6100), according to the manufacturer’s instructions and developed using a DAB Peroxidase (HRP) Substrate Kit (Vector, SK-4100) until the desired staining intensity was reached. Floating sections were mounted on Superfrost plus slides in 0.2 % pork skin gelatine dissolved in 50 mM Tris-HCl pH 7.4, dried at RT overnight and dehydrated in ethanol and xylene before mounting with DPX mounting medium (Merck). Images were acquired using 20x magnification on a ZEISS Axio Scan.Z1 microscope and processed using ZEISS ZEN lite software. For immunofluorescence, sectioned brains were blocked for 1 h before primary antibodies were applied overnight. AlexaFluor 488 secondary antibody was used to detect OLIG2 (1:1000; Invitrogen, Carlsbad, CA) for 60 min at room temperature together with Hoechst 33258 (1:1000; Sigma) to detect cell nuclei. All secondary antibodies were diluted in block solution. Floating sections were transferred onto Superfrost plus slides (BDH Laboratory Supplies) and air dried before being coverslipped with Dako fluorescent mounting medium.

### Analysis of corpus callosum and cortex thickness and cell counting

Matched sections at specified Bregma levels were measured using the Zeiss Blue software measurement length tool. Counting boxes were positioned onto matched slices in the barrel field of the primary somatosensory cortex perpendicular to the corpus callosum. In each box, cells were counted from the border of the corpus callosum to the pial surface. The length of the box was adjusted to fit the cortical thickness while the width was not changed. All cells positive for NeuN within the box were counted and cell numbers displayed as crude numbers in the box as well as density considering the area of the boxes analysed. Four animals per genotype were used.

### Electron microscopy (EM)

EM was performed as described in Ref.^88^. Briefly, mice were prepared for EM by PBS perfusion followed by 2.5% (v/v) glutaraldehyde and 2% (w/v) paraformaldehyde in 0.1 M sodium cacodylate buffer (pH 7.6). Brains were removed and kept at 4°C for 48 h in the same fixative, before being washed with 0.1 M cacodylate buffer, rinsed briefly with 0.1 M phosphate buffer and shipped to Japan at ambient temperature. Sagitally-sectioned brains were processed for EM. After immersion in a 1% (w/v) osmium tetroxide solution for 2 h at 4°C, the specimens were dehydrated through a graded alcohol series and embedded in Epon 812 (TAAB Laboratories, UK). The required area was trimmed, and ultrathin sections were cut, collected on a platinum-coated glass slide, stained with uranyl acetate and lead citrate and imaged in a scanning electron microscope equipped with a back-scattered electron beam detector (Hitachi SU8010) at 1.5 kV accelerating voltage. EM images of sagittal sections of corpus callosum were taken at 15000x magnification. To estimate g-ratios, the circumference of the axon and the external circumference of the myelin sheath were measured using ImageJ, avoiding paranodal areas. The diameters of axon and myelin sheath were calculated from these circumferential measurements, assuming a circular profile, and the g-ratio was calculated.

### Behavioural analysis

Behaviour testing was carried out as previously described ^89^. 12-week-old mice on mixed C57BL/6 x Sv129 background were used in this study. All animals were kept on a 12 h light-dark cycle with food and water accessible ad libitum. The indicated numbers of WT mice and their *Pten*^+/R173C^ littermates were tested in the open field test, three chambers test, hole board and marble burying. Mice tested in behavioural experiments were habituated to handling and the experimenter for 9 days before the start of behavioural experiments to reduce stress and habituated mice were generally handled by cupping. Mice were analysed in three cohorts and male and female mice were tested on separate days. During experiments white noise was played at around 50 Db to mask external noises and all experiments were video recorded. Before every experiment, mice were brought into the behavioural room 30 min before the first experiment for habituation and animals that had finished a test were transferred into a new cage in order not to influence the behaviour of the remaining experimental animals. Directly before the first experimental animal was tested and after every mouse the set up (open field, sociability chamber, hole board) was cleaned thoroughly with water and 70% ethanol in a consistent manner for every animal. Open field: the experimental animal was placed in an open field box measuring 30 cm x 30 cm. The walls were white and 40 cm high so that the animal could not get any visual clues. The mouse was allowed to roam freely for 30 min to explore the arena without any prior habituation. Distance, time spent in different areas and the time moving were analysed using Ethovision software while faecal boli were counted manually. Sociability: The experimental animal was placed into the central compartment of Crawley’s chamber for 5 min with closed doors for habituation. After 5 min empty cages were placed into the middle of the left and right compartments and a stranger mouse was gently put into one of the cages (counterbalance left and right sides). The stranger mouse matched the subject mouse with regards to the breeding background, age and gender and had no contact with the subject mouse ever before. Both doors were opened at the same time and the experimental mouse was allowed to explore the area for 10 min. A stopwatch was used to determine the time the subject mouse spent closely interacting with either cage (direct contact/sniffing/cage climbing). The time spent with the empty cage versus the first stranger was measured, referred to as sociability. Calculation of discrimination indices: (time spent with mouse – time spent with empty cage)/(sum of the time spent with both). Hole board: since the experimental animals were familiar with the open field arena from the open field experiment, they were only briefly re-exposed to the empty arena for 10 min on day 1 of the hole board test. On the next day a hole board with 16 holes evenly spaced was placed into the arena, the subject mouse was put into the middle of the field and was recorded for 15 min. A manual cell counter was used to determine the total number of head dips into the holes. Marble burying: a cage was filled with 5cm of bedding that was tamped down lightly to produce an even surface. 12 glass marbles were distributed evenly spaced out in the cage and the animal was allowed to roam freely for 15 min. Marbles that were buried to 2/3 of their depth with bedding were counted.

### Statistical analysis

GraphPad Prism (San Diego, CA, United States) was used for statistical analysis. The method used for individual datasets is indicated in the figure legends.

## ACKNOWLEDGEMENTS

Personal funding was from the Jean Shanks Pathological Society (Clinical PhD fellowship; to V.R.), CRUK UCL Centre Non-Clinical Training Awards (C416/A29287 to G.C. and CANTAC721\100022 to F.B.) and the European Commission (H2020-MSCA-IF-2018 GA: 838559; to S.C.). Research in the B.V. laboratory was supported by PTEN Research (UCL-16-001, UCL-20-001), Cancer Research UK (C23338/A25722) and the UK Biotechnology and Biological Sciences Research Council (BB/W007460/1). Research in N.R.L. laboratory was supported by PTEN Research. Research in the M.G.H.S. group was supported by Fondation ARC pour la recherche sur le cancer (ARCPJA2022060005118) and Ligue Contre le Cancer (Comité de Paris). The UCL Cancer Institute Translational Technology Platforms are supported by the CRUK UCL Centre Award (C416/A25145). The N.K. lab is funded by grants from the UK Biotechnology and Biological Sciences Research Council (BB/N009061/1) and the Wellcome Trust (108726/Z/15/Z). M.T. and K.S. are supported by PTEN Research and the NIHR Cambridge Biomedical Research Centre (NIHR203312). The PHTS Patient Registry UK is funded by PTEN Research (UOC-17-001). G.R.M. and R.L.W. were supported by MRC funding (MC-A024-5PF91 to R.L.W.). We thank staff at the Technical Support Centre for Life Science Research in IMU for excellent technical support with EM processing. We also thank the UCL Research Capital Infrastructure Fund (RCIF) and the National Institute for Health Research University College London Hospitals Biomedical Research Centre for upgrade of the UCL Cancer Institute Microscopy facility and Dr Maria Whitehead and Dr Paul Elvin and members from the B.V. group for excellent feedback on the manuscript.

## Author contributions

PT, NRL, NK and BV conceived the study, PT, VR, SEC, GAEC, FB, MAD, GC, ZA,KT, NK, VAG, EF, DB, ML,WP, MA, ZV, SH, LC, AA, GRM and MGHS performed and analysed the experiments, CLS performed the histological analysis, RLW, AMF and JH provided reagents/resources for the study, PT, NK and BV wrote the manuscript.

