## Supplementary Figures for "A mouse model of PTEN Hamartoma Tumour Syndrome reveals that loss of the nuclear function of PTEN drives macrocephaly, lymphoid overgrowth, and late-onset cancer"

*Corresponding authors

#### Figure S1

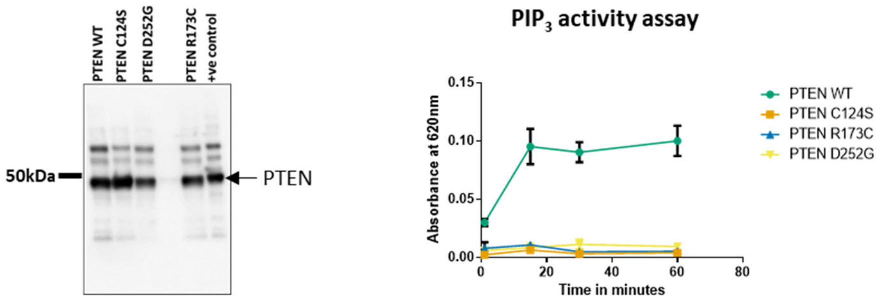


**Figure S1. Characterisation of PTEN-R173C.** Recombinant untagged PTEN wild-type (WT) and mutant proteins were purified from *E. coli* and used in a $PIP_3$ lipid phosphatase assay. *Left*, Coomassie blue-stained SDS-PAGE gel showing the level of purified recombinant protein. A previous batch of purified PTEN protein is used as a positive control. *Right,* Graph showing PTEN activity measured as absorbance at 620 nm in a malachite green assay. Data shown as mean ± SEM from n=3.

### Figure S2

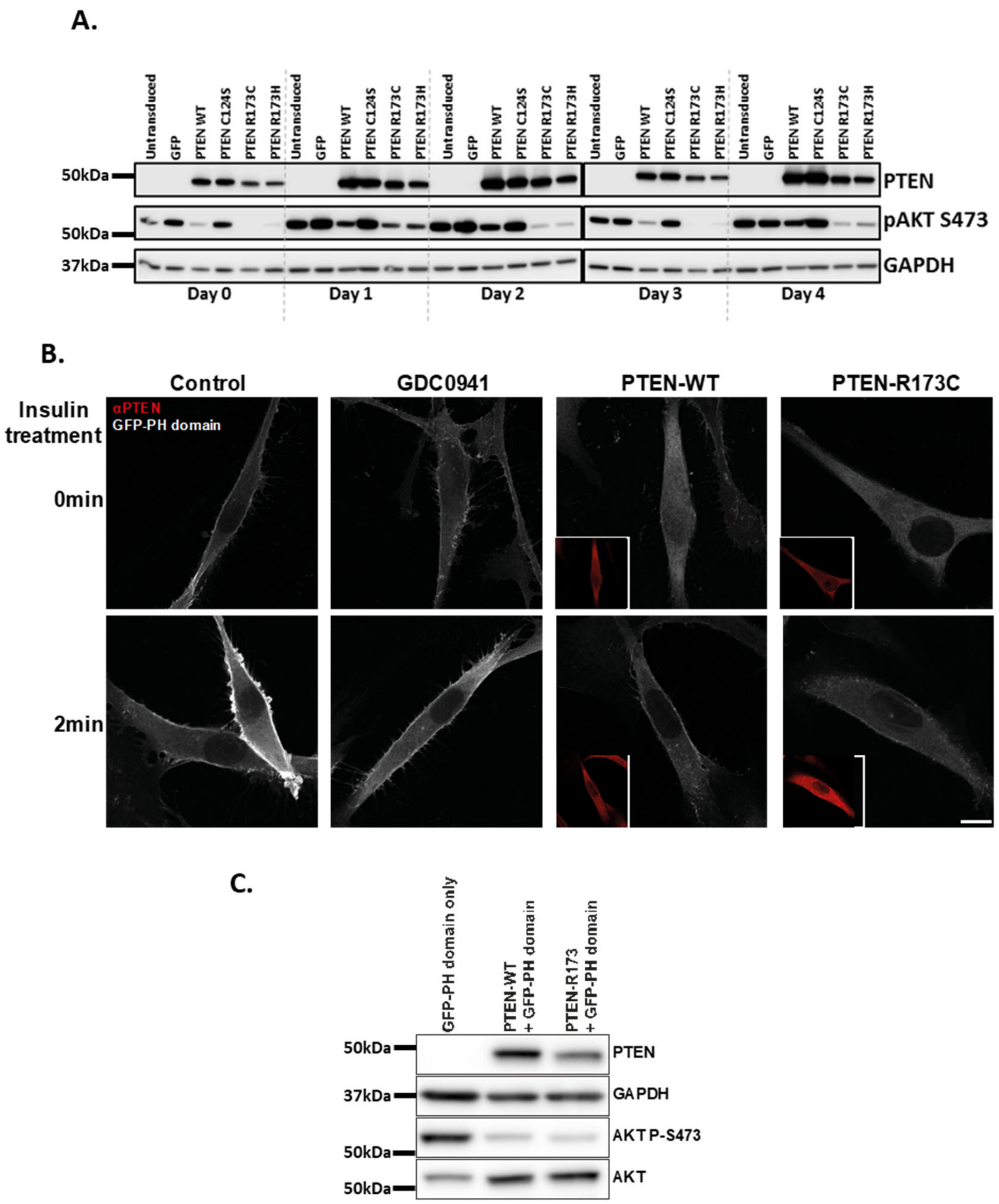


**Figure S2. Characterisation of PTEN-R173C. (A)** Immunoblots of PTEN protein and pAKT-473 from the cells in Fig. 1C. **(B)** U87 cells, transiently transduced with lentivirus encoding a $PIP_3$ biosensor, with or without PTEN-WT or PTEN-R173C, were pre-treated $\pm$ GDC-0941 (1 µM) for 1 h, followed $\pm$ insulin treatment (100 nM) for 2 min. The cells were stained with antibodies to PTEN (red inset) and imaged by confocal microscopy. Quantification of the mean fluorescence intensity (MFI) of the $PIP_3$ biosensor is shown in Fig. 1D. **(C)** Immunoblots of PTEN, AKT P-S473, AKT and GAPDH from the cells in (B) and Fig. 1D.

### Figure S3

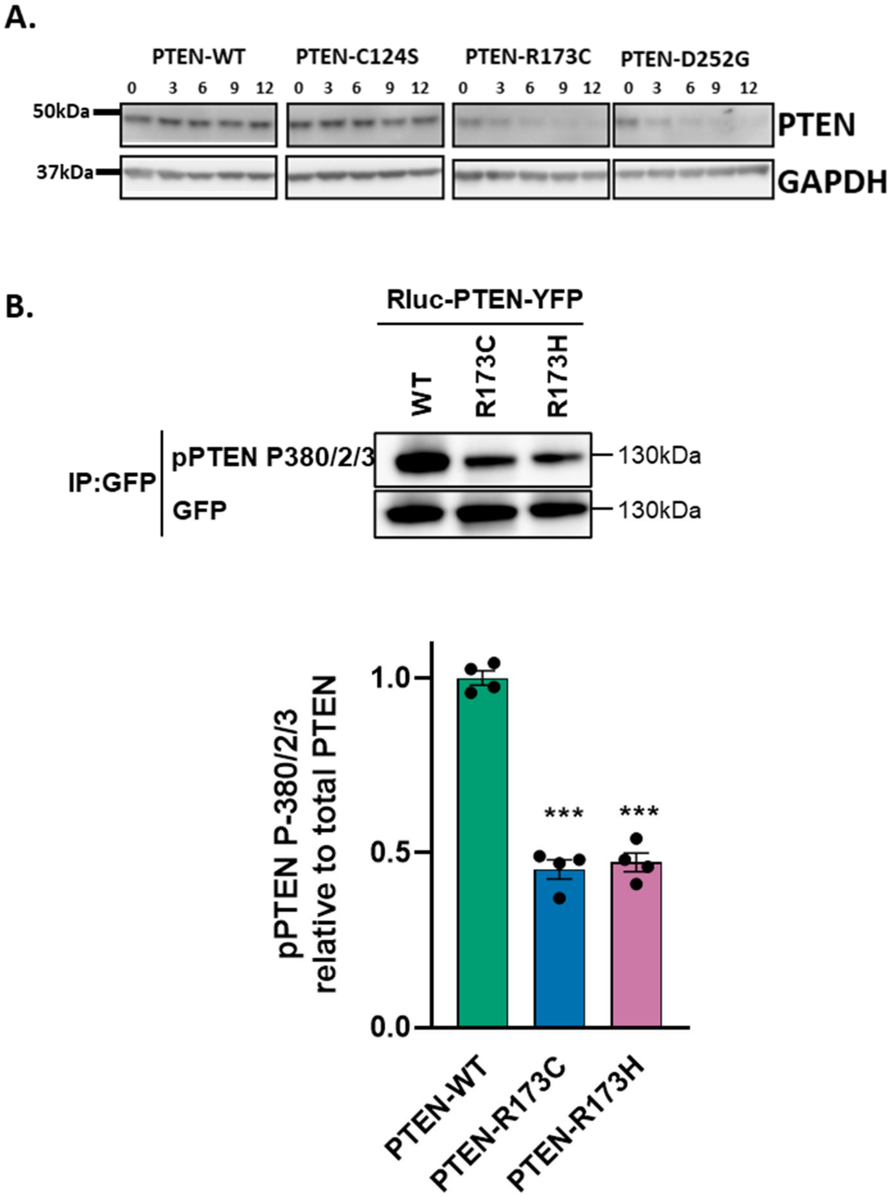


**Figure S3. Characterisation of PTEN-R173C.** (**A**) Transient lentiviral expression of PTEN wild-type (WT) or the indicated PTEN mutants in U87 cells. The cells were treated with cycloheximide, and PTEN protein expression levels were determined by immunoblotting at the indicated time points. Representative blots are shown. (**B**) Lysates of HEK-293T cells expressing Rluc-PTEN-WT-YFP, Rluc-PTEN-R173C-YFP or Rluc-PTEN-R173H-YFP were immunoprecipitated using anti-GFP antibodies and immunoblotted for PTEN-P-S380/T382/T383. Representative immunoblots from n=4 showing phosphorylated and total PTEN levels. Graph shows quantification of 4 experiments shown as mean ± SEM. Statistical analysis was performed using one-way ANOVA. p-values: *** $p<0.01$.

### Figure S4

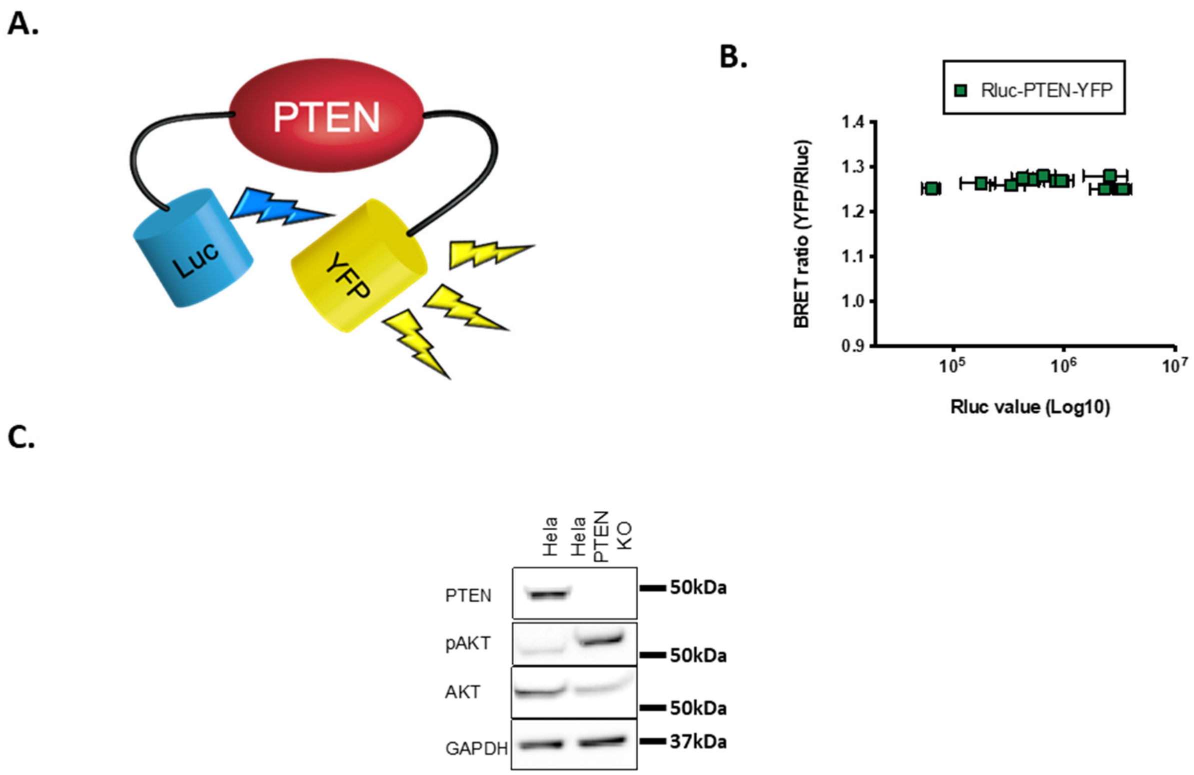


**Figure S4. BRET analysis of PTEN.** (**A**) Cartoon showing the intramolecular bioluminescent resonance energy transfer (BRET)-based biosensor (Rluc-PTEN-YFP). The biosensor can reveal dynamic changes in PTEN conformational rearrangement and function, due to shifts in energy transfer between the donor/acceptor couple. (**B**) The conformational readout (measured as BRET ratio) is independent from expression levels of the biosensor, as seen by a stable BRET signal when the biosensor (Rluc-PTEN-YFP) is expressed over 2 log differences (measured as Rluc value). (**C**) Validation of PTEN knock out by immunoblotting in the PTEN-KO HeLa cell line.

### Figure S5

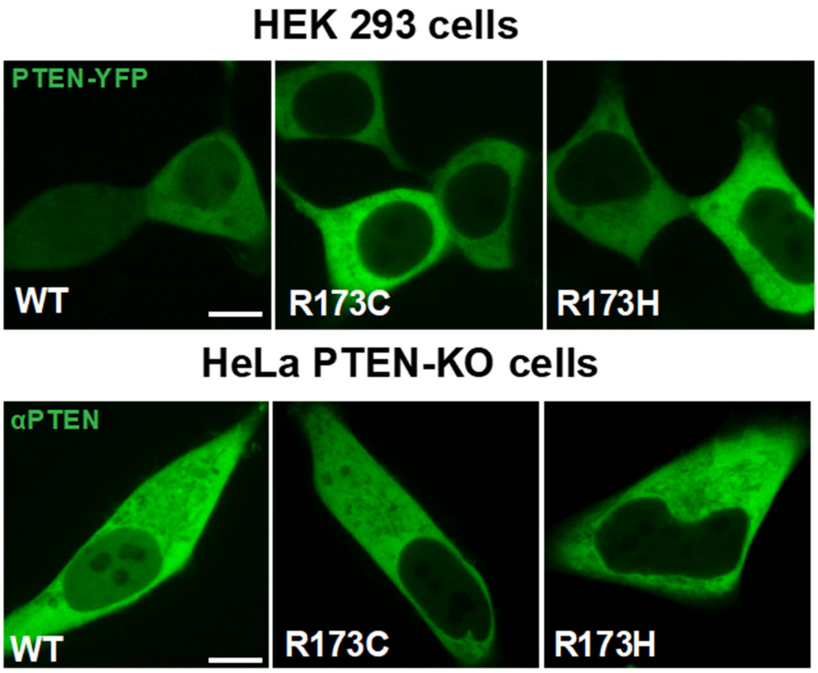


**Figure S5. Sub-cellular distribution of PTEN.** Live HEK-293T cells or PTEN-KO HeLa cells expressing Rluc-PTEN-WT-YFP, Rluc-PTEN-R173C-YFP or Rluc-PTEN-R173H-YFP were imaged in 4-well µ slides (IBIDI) by confocal microscopy to assess the subcellular localisation of PTEN-YFP. Figure shows representative images from n=3. Scale bar: 10 µm.

### Figure S6

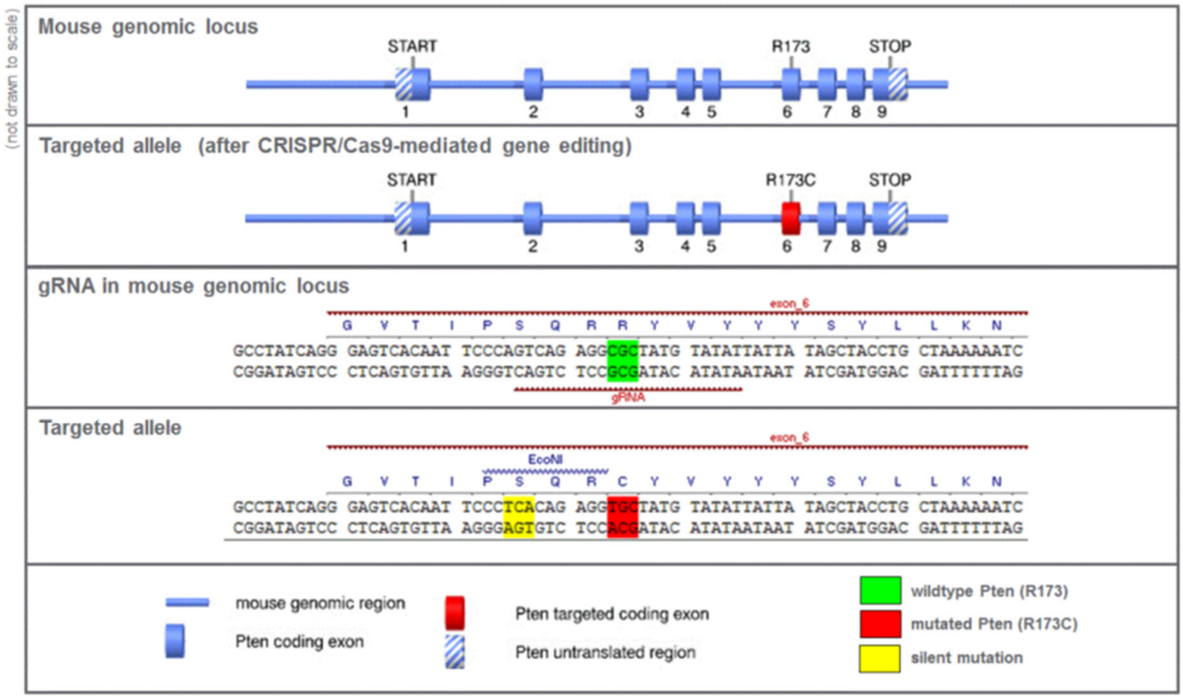


**Figure S6. Generation and characterization of *Pten*$^{+/R173C}$ mice.** The R173C mutation was introduced into the mouse *Pten* locus using a CRISPR-Cas9 approach. gRNA was designed to contain the R173C mutation (c.517C>T) and a silent mutation containing the *Eco*NI restriction site upstream of the R173 site, to allow discrimination of wild-type and knock-in allele by PCR analysis. The figure above shows the gRNA sequence and the targeting strategy. gRNA was microinjected into C57BL/6J mouse zygotes along with recombinant Cas9. The zygotes were then transferred to pseudo-pregnant mothers. The litters born (founder animals) were sequenced to check for the incorporation of the R173C mutation and percentage of mosaicism. They were then used as a breeder to generate F1 animals. F1 males positive for the R173C mutation and negative for any off-target mutations were used to generate further animals by IVF using C57BL/6J females.

Figure S7

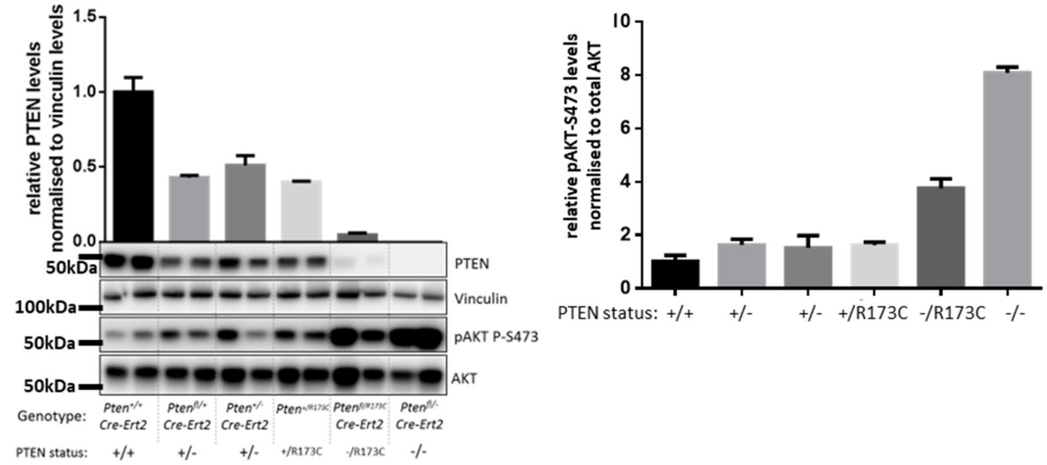


**Figure S7. *In vivo* characterisation of PTEN-R173C.** (**A**) Immunoblots of PTEN protein (quantification in graph on the left) and pAKT-473 (quantification in graph on the right) from MEFs of the indicated genotypes. Quantification shown as mean ± SEM from 2 MEF lines for each genotype. The MEFs were generated from mice on a C57BL/6J background.

Figure S8

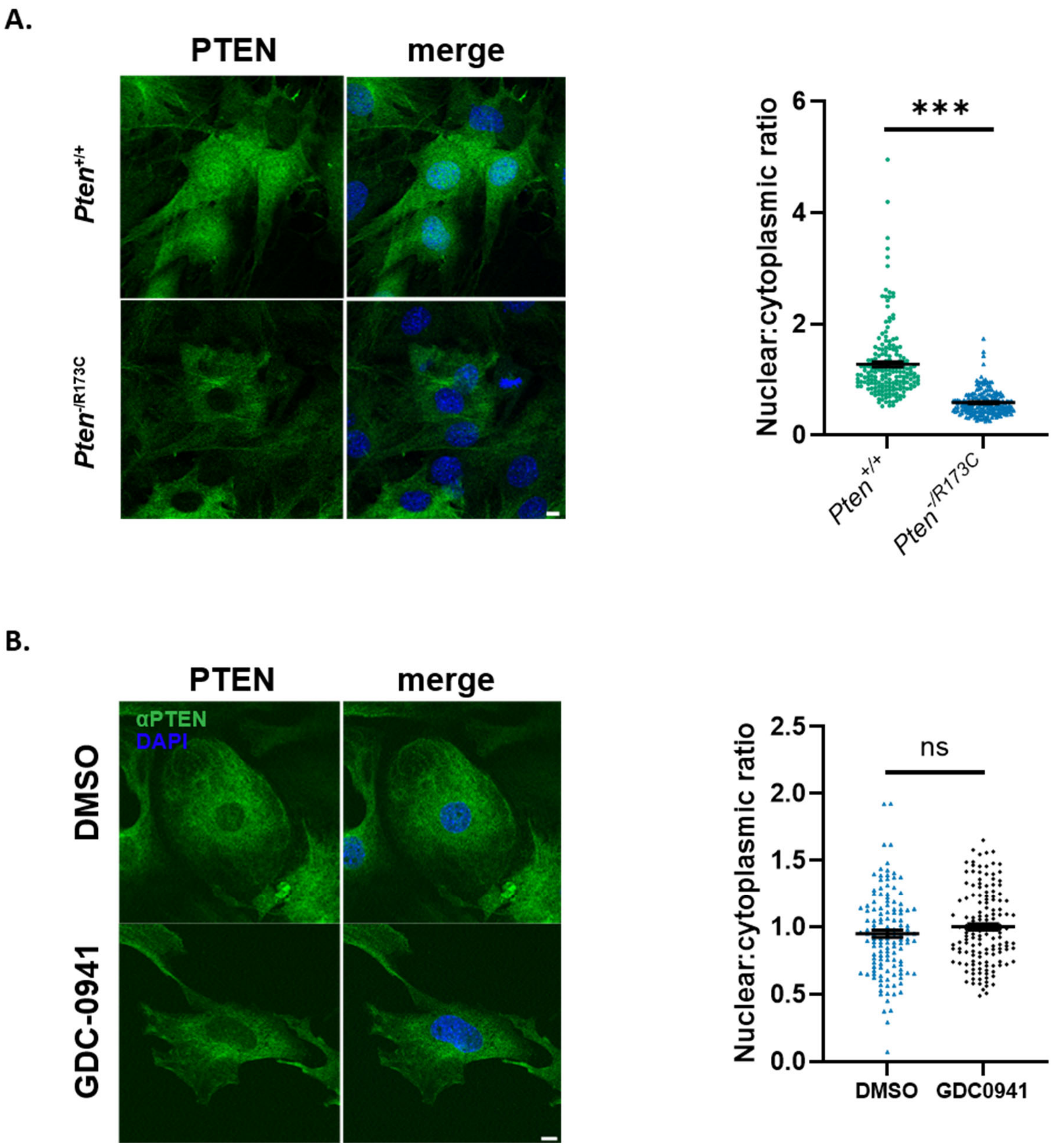


**Figure S8. Subcellular localisation of PTEN-R173C.** (**A-B**) MEFs of the indicated genotype were immunostained with antibodies to PTEN (green) and DAPI (blue) and imaged by confocal microscopy. Scale bar: 10 µm. Cells in (B) were pre-treated with or without GDC-0941 (1 µM) for 1 h, as indicated. Graphs show nuclear:cytoplasmic ratio of the GFP MFI (mean ± SEM from 3 independent experiments). Statistical analysis was performed using Unpaired Student's t-test. p-values for all statistical analysis: *, p< 0.05; **, p<0.01; ***, p<0.001; ****, p<0.0001. The MEFs were generated from mice on a C57BL/6J background.

### Figure S9

#### A. Uterus

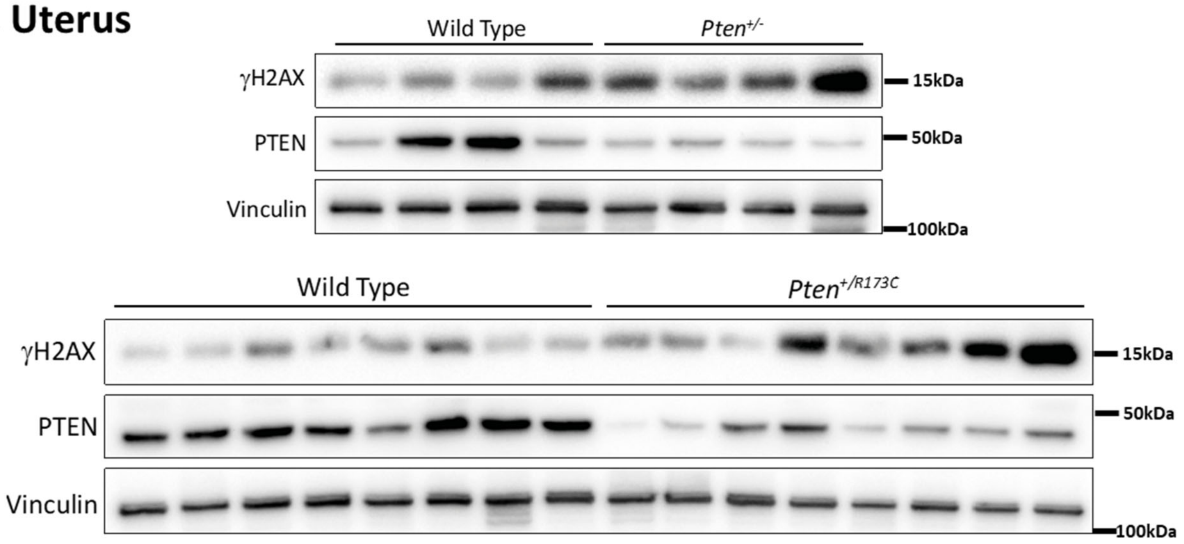


#### B. Kidney

Males

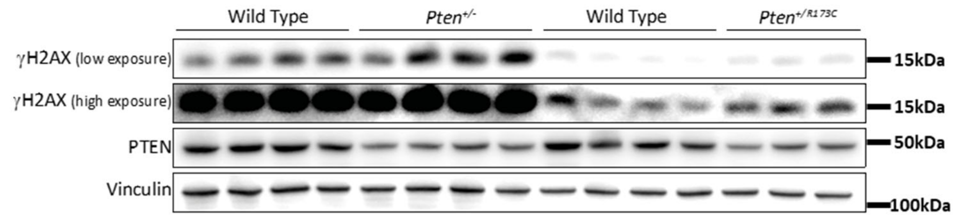


Females

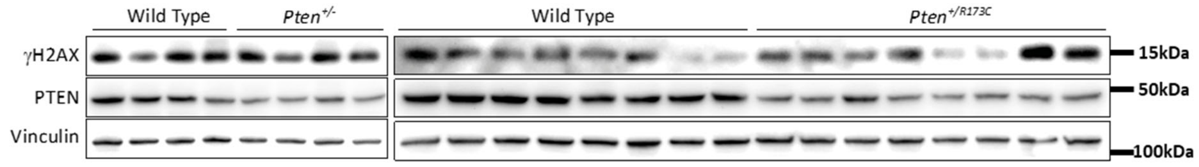


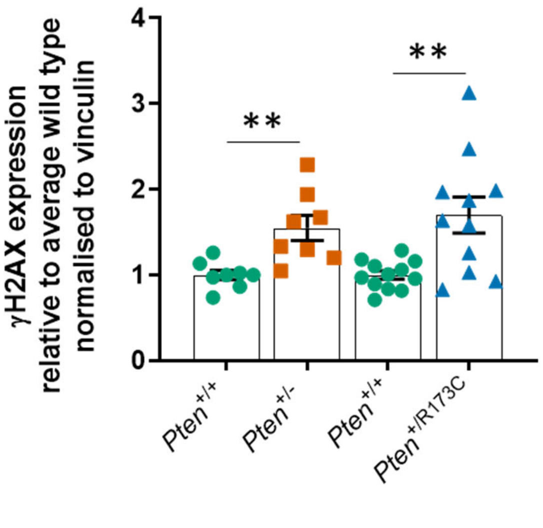

#### C. Liver

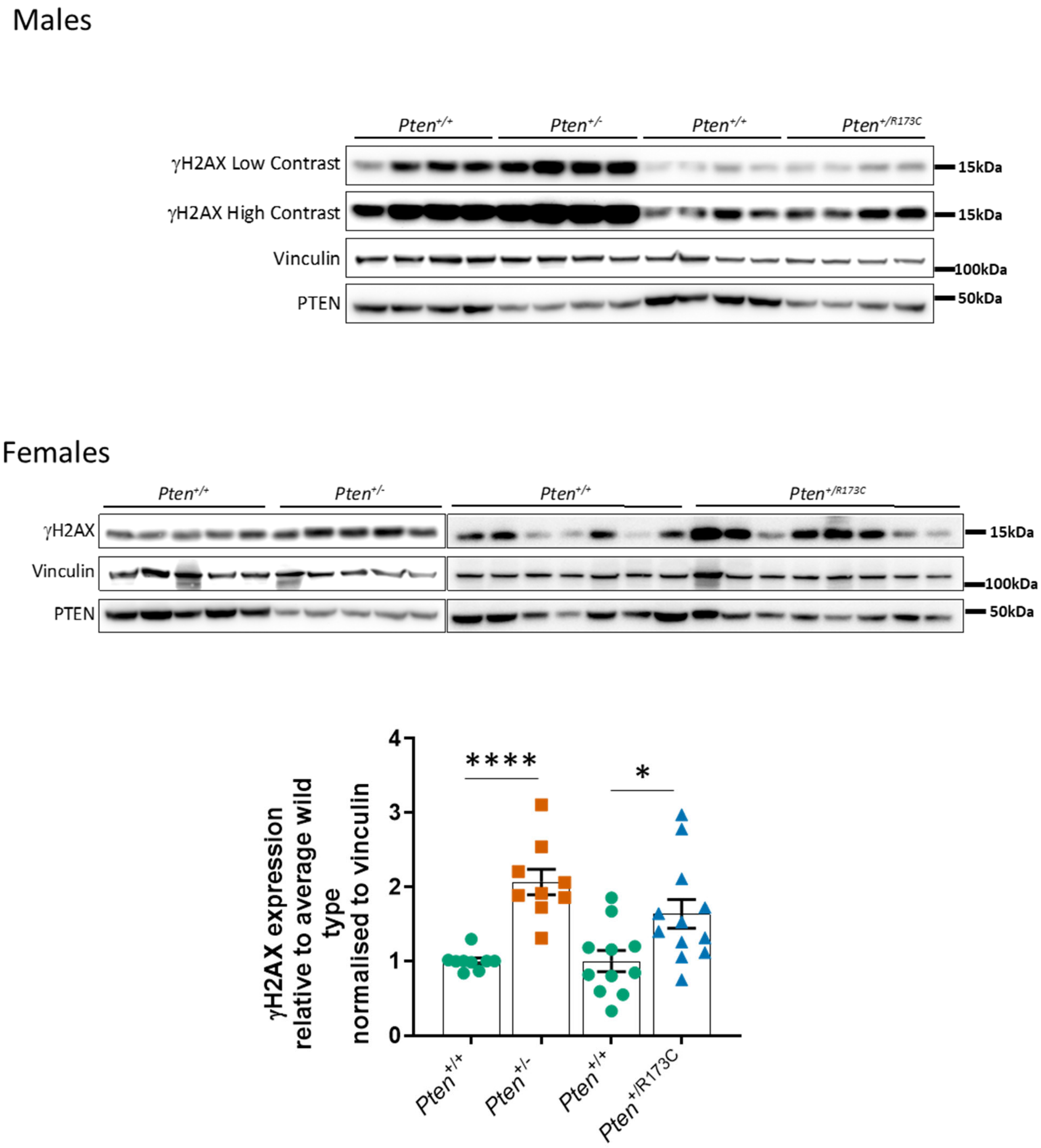


**Figure S9. *In vivo* characterisation of PTEN-R173C mice.** 6-week-old mice on a C57BL/6J background of the indicated genotypes were treated with 7Gy γ-radiation and protein extracts from tissues indicated were used for immunoblotting. Immunoblots showing levels of γH2AX in (**A**) uterus (quantification in Fig. 2G), (**B**) kidney and (**C**) liver. Graphs show quantification of immunoblots. Bars show mean ± SEM. Pairwise comparisons were performed using Mann–Whitney U test. p-values: *, $p < 0.05$; **, $p<0.01$; ***, $p<0.001$; ****.

### Figure S10

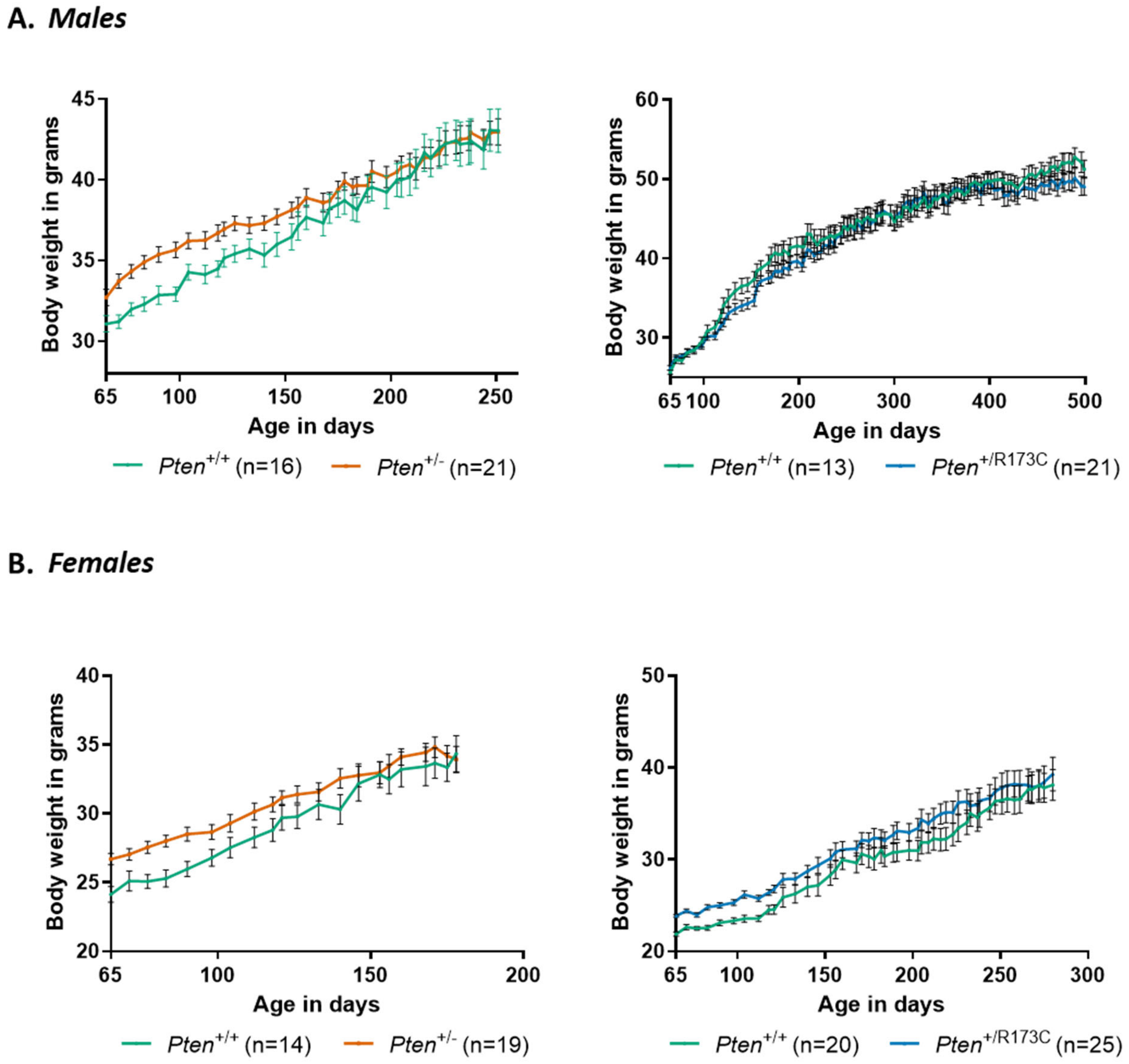


**Figure S10. Age-dependent body weight of *Pten* mutant mice.** *Pten*$^{+/R173C}$ and *Pten*$^{+/-}$ mice, and their littermate controls on a mixed C57BL/6J x Sv/129 background were allowed to age. The mice weighed once a week. Data show mean ± SEM.

### Figure S11

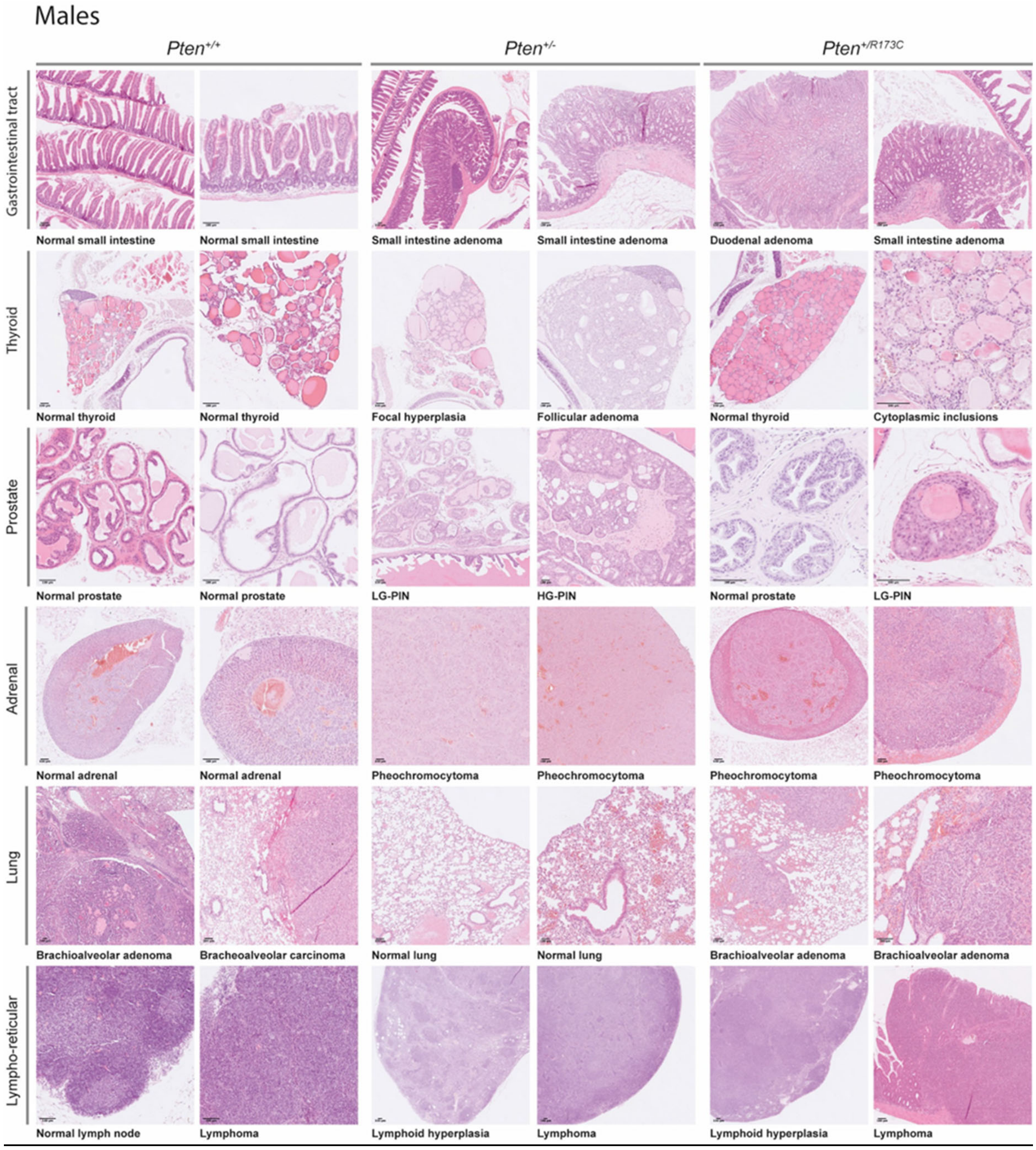


**Figure S11. Histopathological analysis of male *Pten*$^{+/R173C}$ mice.** *Pten*$^{+/R173C}$ and *Pten*$^{+/-}$ mice, and their littermate control *Pten*$^{+/+}$ mice, on a mixed C57BL/6J x Sv129 background were allowed to age. Mice were euthanised for welfare reasons (ill health or masses with a combined size of ≥ 1.4 $cm^2$ surface area) or at a specified age. Representative photomicrographs of H&E-stained sections of indicated tissues from male mice. Scale bar: 100 µm.

### Figure S12

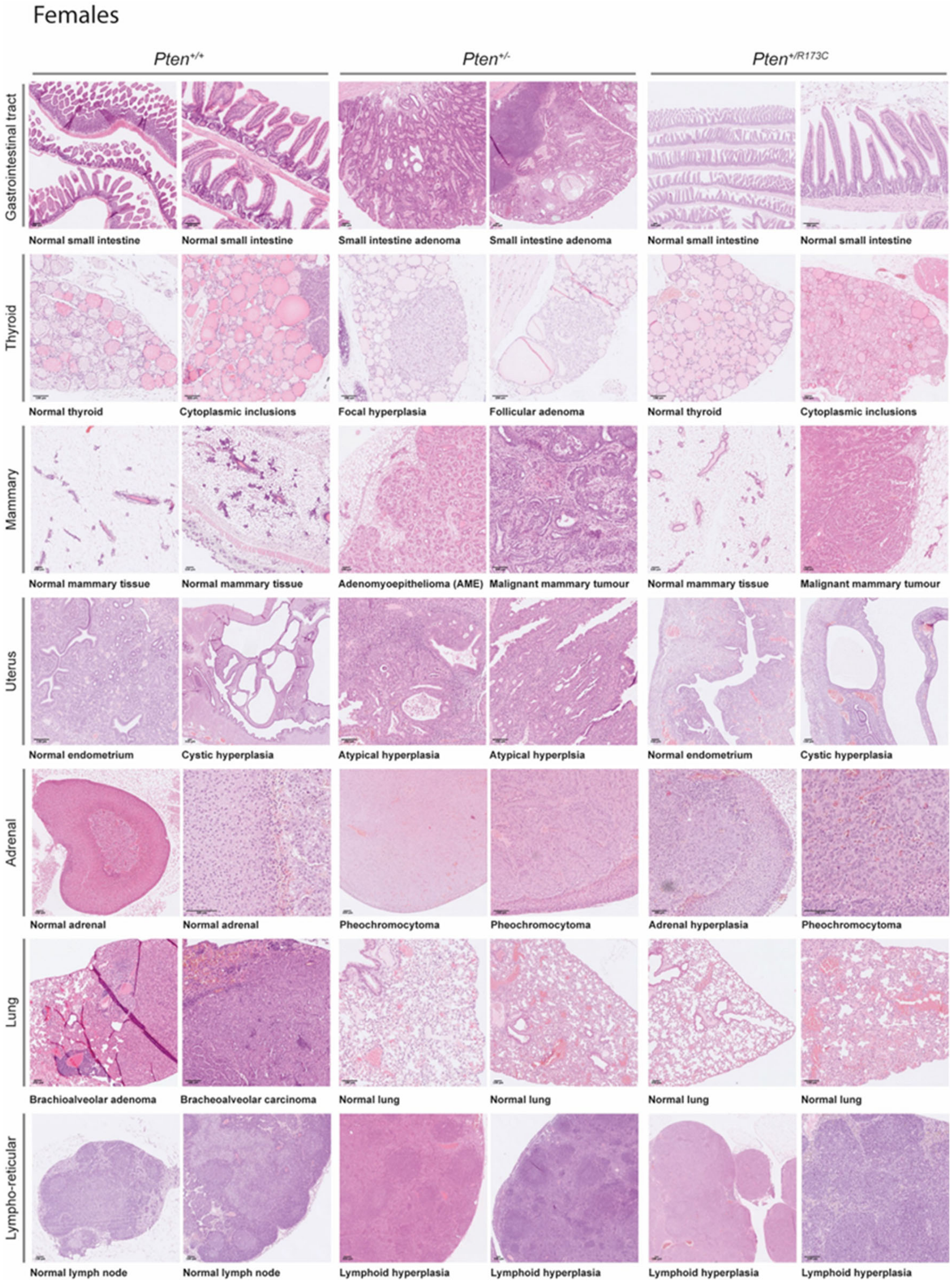


**Figure S12. Histopathological analysis of female *Pten*$^{+/R173C}$ mice.** *Pten*$^{+/R173C}$ and *Pten*$^{+/-}$ mice, and their littermate control *Pten*$^{+/+}$ mice, on a mixed C57BL/6J x Sv129 background were allowed to age. Mice were euthanised for welfare reasons (ill health or masses with a combined size of ≥ 1.4 cm$^2$ surface area) or at a specified age. Representative photomicrographs of H&E-stained sections of indicated tissues from female mice. Scale bar: 100 µm.

### Figure S13

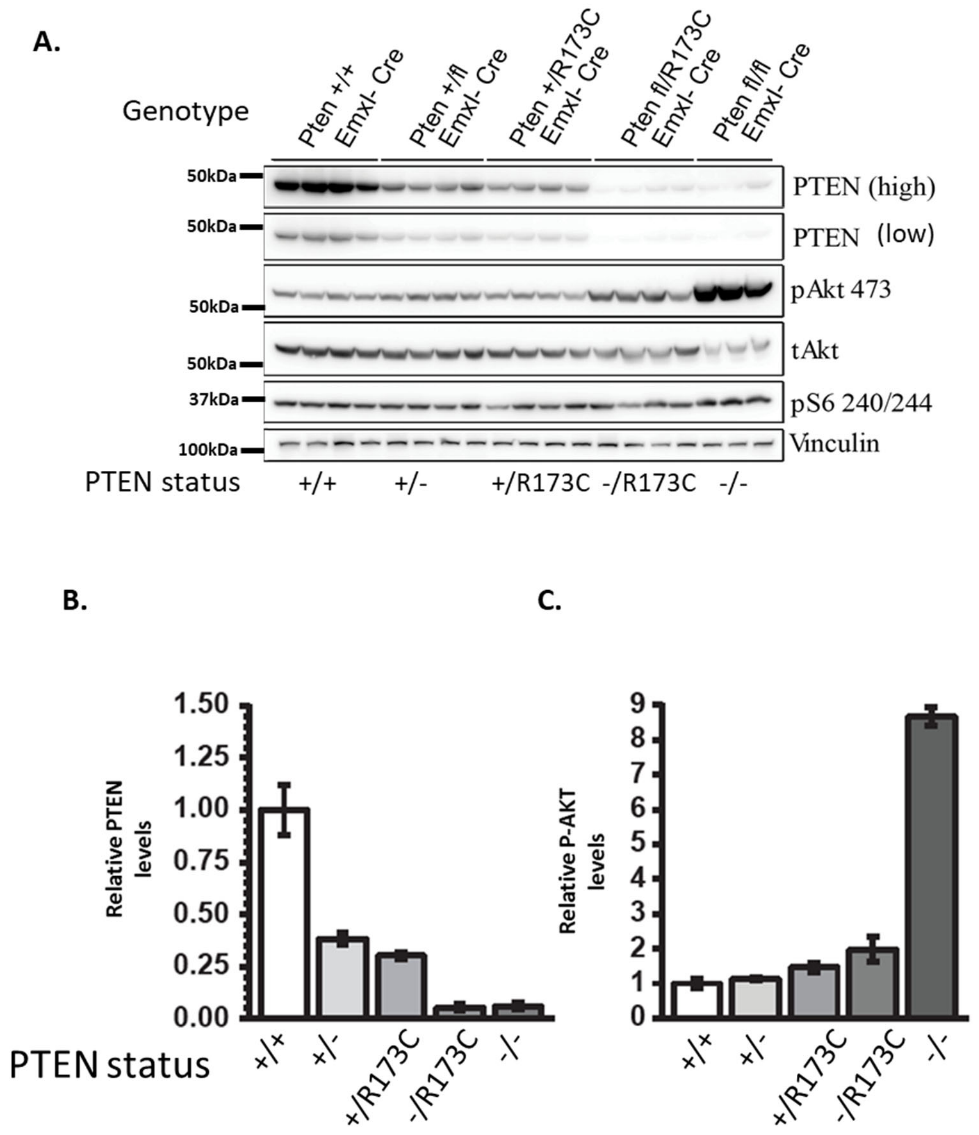


**Figure S13. Expression of PTEN and downstream pathway component in brain cortical lysates.** (**A**) Immunoblots of PTEN protein and downstream PI3K pathway components in cortico-hippocampal lysates of postnatal pups (P1/P2) from the indicated genotypes. (**B**) Quantification of PTEN protein, normalised to the vinculin loading control. (**C**) Quantification of pAKT normalised to total AKT. n=4 for all mutants except for *Pten*$^{fl/fl}$;*Emx1-Cre* where n=3. All mice were on a C57BL/6J background.

### Figure S14

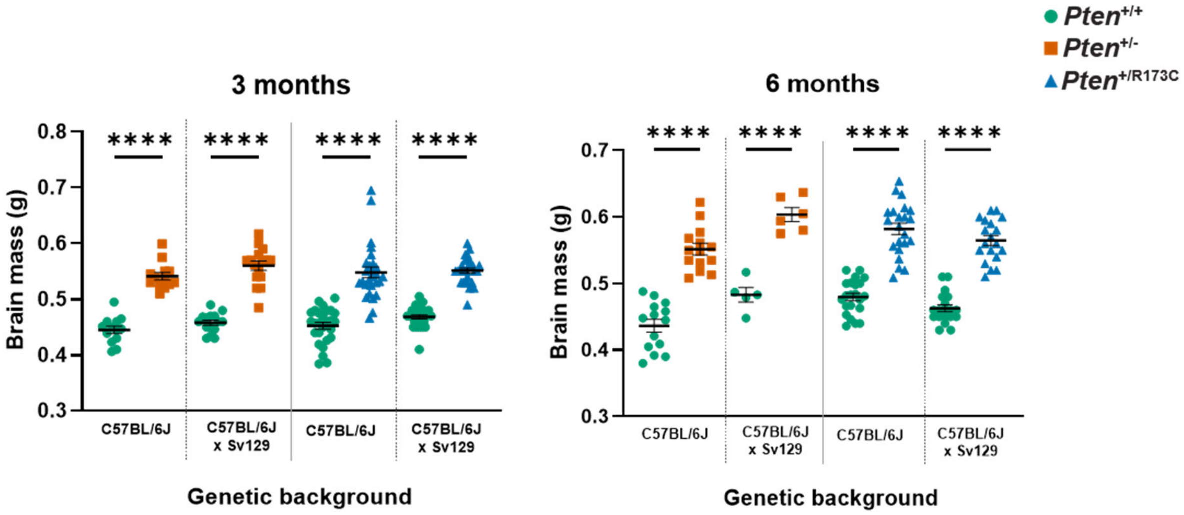


**Figure S14. Analysis of brain mass in mice.** Brain mass of *Pten*$^{+/-}$, *Pten*$^{+/R173C}$ and *Pten*$^{+/+}$ littermate controls of both sexes at 3 and 6 month of age on the indicated genetic backgrounds. Data show mean ± SEM, statistical analysis done using One-way ANOVA. p values: ****p<0.0001, ***p<0.0005, **p<0.01, *p<0.05.

### Figure S15

**A.**

**B.**

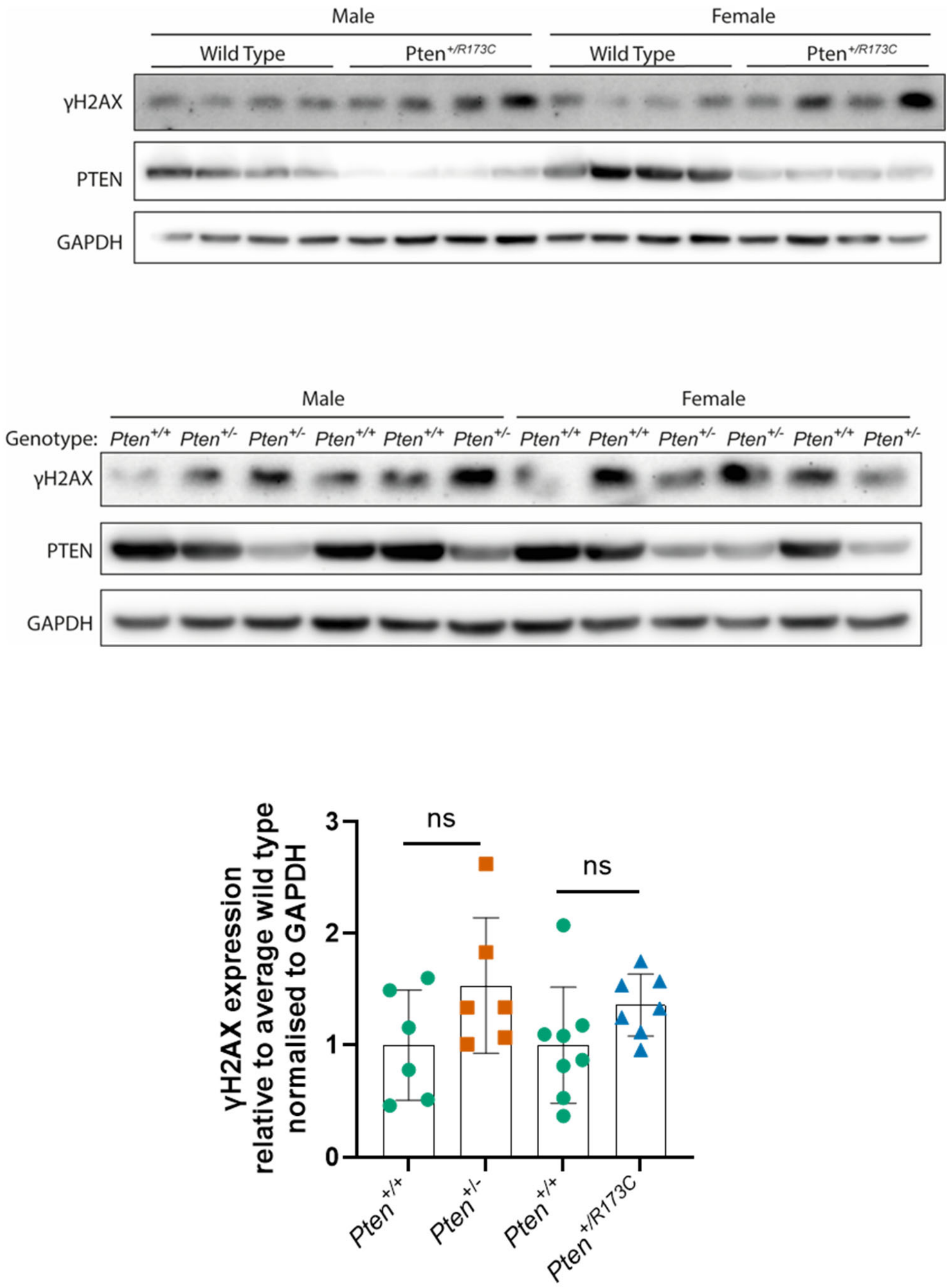


**Figure S15.** ***In vivo*** **characterisation of PTEN-R173C mice.** (**A**) 6-week-old mice on a C57BL/6J background of the indicated genotypes were treated with 7Gy γ-radiation and protein extracts from brain were used for immunoblotting. Immunoblots showing levels of γH2AX. (**B**) Graphs show quantification of immunoblots. Bars show mean ± SEM. Pairwise comparisons were performed using Mann–Whitney U test. ns = non-significant.

#### Figure S16

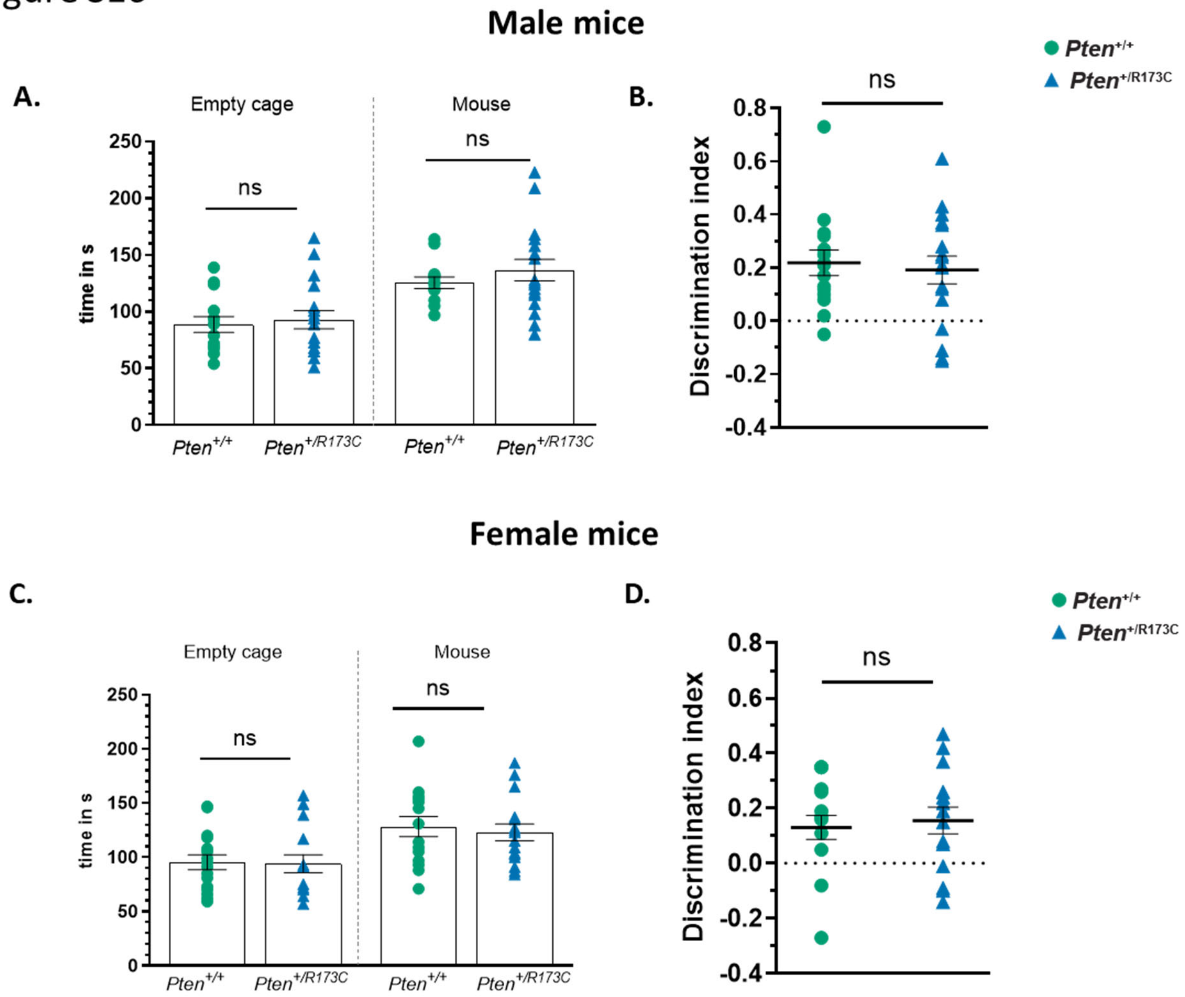


**Figure S16. Lack of sociability deficits in *Pten*$^{+/R173C}$ mice in Crawley's sociability test.** Analysis of sociability of male (**A**) and female (**C**) *Pten*$^{+/R173C}$ mice and littermate controls. In Crawley's three chamber test, mice were analysed for their preference of a mouse *versus* an empty cage. Data show mean ± SEM. Ordinary one way ANOVA. p values: ***$p<0.0005$, **$p<0.01$, *$p<0.05$. (**B and D**) Discrimination indexes for male and female WT and *Pten* mutant animals analysed in A and C. Males: n=14 for *Pten*$^{+/+}$ and n=17 for *Pten*$^{+/R173C}$ mice. Females: n=16 mice of each genotype. Data show mean ± SEM, statistical analysis done using Unpaired Student's t-test. ns; non-significant.

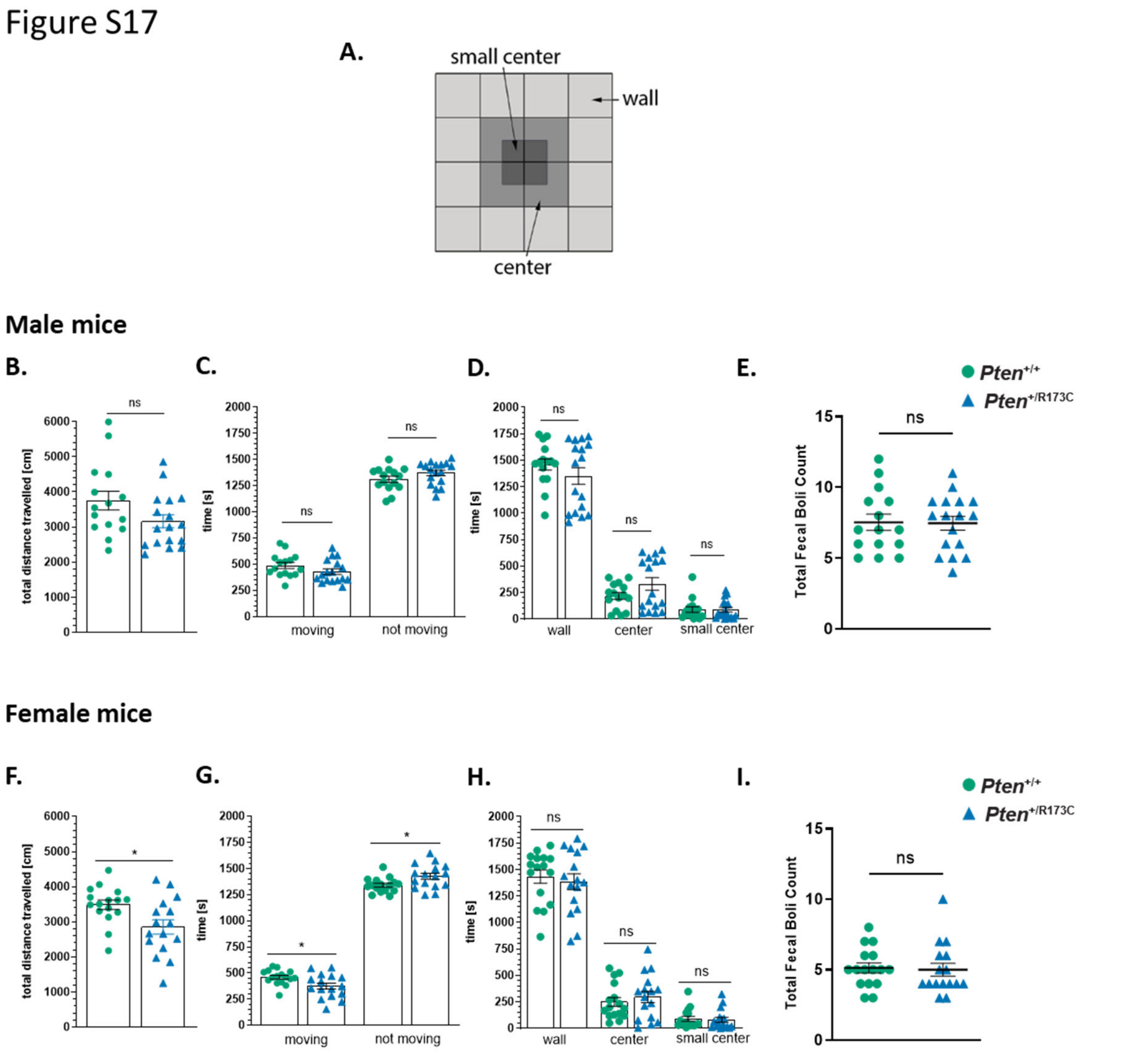


**Figure S17. Female *Pten*$^{+/R173C}$ mice show reduction in exploratory locomotion in the open field test.** **A.** Scheme of the open field arena. Parameters measured in the open field test for male (**B, C, D and E**), and female (**F, G, H, I**) *Pten*$^{+/R173C}$ mice and their littermate WT controls. Total distance travelled (**B and F**), time spent moving and not moving (**C and G**), time spent in each of the three zones as indicate in A (**D and H**) and defecation during the test (**E and I**). There were no differences between *Pten*$^{+/R173C}$ and control male mice. Female *Pten*$^{+/R173C}$ mice spent less time moving and travel smaller distances compared to their WT littermate controls indicating possible exploratory locomotion deficits. Males: n=15 for WT and n=17 *Pten*$^{+/R173C}$ mice. Females: n=16 mice of each genotype. Data show mean ± SEM. Two-tailed Mann-Whitney U test. p values: *p<0.05. ns = non-significant.

#### Figure S18

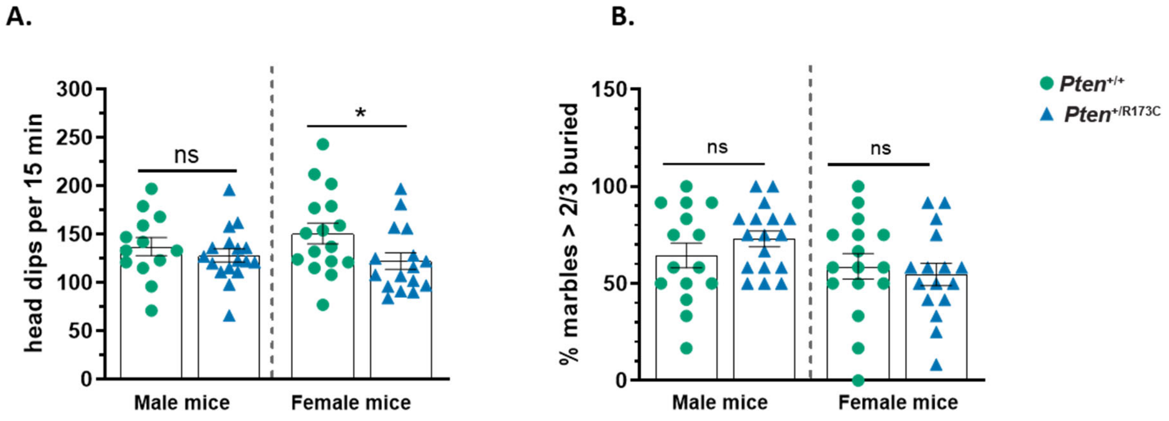


**Figure S18. Female *Pten*$^{+/R173C}$ mice show reduction in exploratory locomotion in the hole board test.** **A.** Hole board test: female *Pten*$^{+/R173C}$ mice made significantly fewer head dips than *Pten*$^{+/+}$ littermate controls. Males: n=13 *Pten*$^{+/+}$ and 17 *Pten*$^{+/R173C}$ mice. Females n=16 mice per genotype. Data show mean ± SEM. Two-tailed Mann-Whitney U test. *p<0.05. **B.** Marble Burying test: there was no significant difference between *Pten*$^{+/R173C}$ and *Pten*$^{+/+}$ controls of either sex in this test. Males: n=15 for *Pten*$^{+/+}$ and n=17 for *Pten*$^{+/R173C}$ mice. Females n=16 mice per genotype. Data show mean ± SEM, statistical analysis done using Two-tailed Mann-Whitney U test. ns = non-significant.
