## Supplementary tables for "A mouse model of PTEN Hamartoma Tumour Syndrome reveals that loss of the nuclear function of PTEN drives macrocephaly, lymphoid overgrowth, and late-onset cancer"

[15]Exepathology, Exmouth, UK

*Corresponding authors

**Index**

| | |
|---|---|
| **Supplementary table 1** | Clinical findings of PHTS patients with PTEN-R173 mutations |
| **Supplementary table 2** | Clinical findings of PHTS patients with nuclear excluded but catalytically active PTEN variants |
| **Supplementary table 3** | Reason for euthanasia of *Pten* $^{+/+}$, *Pten* $^{+/-}$ and *Pten* $^{+/R173C}$ mice |
| **Supplementary table 4** | Frequency and grade of lesions in male mice |
| **Supplementary table 5** | Frequency and grade of lesions in female mice |
| **Supplementary table 6** | Frequency and grade of genetic background specific and immune findings in male mice |
| **Supplementary table 7** | Frequency and grade of genetic background specific and immune findings in female mice |
| **Supplementary table 8** | DNA constructs |
| **Supplementary table 9** | Primers for site-directed mutagenesis |
| **Supplementary table 10** | Primers used for genotyping |
| **Supplementary table 11** | Antibodies used |
| **Supplementary table 12** | qRT PCR primers |

**Supplementary table 1: Clinical findings of PHTS patients with PTEN-R173 mutations**

| *PTEN* variant | | | | | PHTS patient data | | | | | | References |
|---|---|---|---|---|---|---|---|---|---|---|---|
| Amino acid change | Nucleotide change | HGVS nomenclature | ClinVar ID | ClinVar classification | Patient ID assigned for this study only | Gender | Age | Clinical phenotypes | Other genetic alterations | Additional notes | |
| *p.Arg173Cys* (R173C) | c.517C>T | NM_000314.8(PTEN):c.517C>T (p.Arg173Cys) | VCV000189500 | Pathogenic | P001 | Female | 8 years | Macrocephaly, DD, ASD | | | 1, clinical phenotypes updated for this study by Dr Katherine Lachlan |
| | | | | | P002 | Male | 13 years | Macrocephaly, MR/DD and penile freckling | | | 2 |
| | | | | | P003 | Female | 29 years | Macrocephaly, Endometrial polyp, hyperpigmentation, ovarian cysts, skin Tag | | | 3 |
| | | | | | P004 | Female | 59 years | Macrocephaly, atypical ductal breast hyperplasia, benign ovarian neoplasm, breast fibrocystic disease, ductal breast carcinoma, fibroadenoma of breast, follicular thyroid carcinoma, GI Polyp, goiter, noninfiltrating intraductal carcinoma, ovarian cysts, uterine fibroids, visceral hemangioma | | | 3 |
| | | | | | P005 | Male | 19 years | Macrocephaly, DD, MR, acral keratoses, benign neoplasm of skin, Tan Macules on glans penis and penile shaft | | | 3 |
| | | | | | P006 | Female | 63 years | Macrocephaly, breast disease benign, endometrial polyp, hemangioma of skin, malignant neoplasm of ovary, malignant neoplasm of skin, polyp hyperplastic benign, skin tag, thyroid nodule, uterine fibroids | | | 3 |
| | | | | | P007 | Male | 40 years | Macrocephaly, skin tag, Tan Macules on glans penis and penile shaft | | | 3 |
| | | | | | P008 | Male | 15 years | Macrocephaly, ASD, DD, Tan Macules on glans penis and penile shaft | | | 3 |
| | | | | | P009 | Male | 4 years | Macrocephaly, ASD, dysmorphic features, DD, Tan Macules on glans penis and penile shaft | | | 3 |
| | | | | | P010 | Female | 2 years | Macrocephaly, DD, ASD, Cafe-au-Lait spots | | | 3 |
| | | | | | P011 | Female | 12 years | Macrocephaly, DD | | | 4 |
| | | | | | P012 | Male | 43 years | Macrocephaly, colon adenoma, left ventricular noncompaction cardiomyopathy | | | 5 |
| | | | | | P013 | Male | NR | Macrocephaly | *GATA4 c.778 C>T,p.Arg260Trp JPH2, c.572 C>G, p.Pro191Arg* | Son of P012 | 5 |
| | | | | | P014 | Male | 1 year | Macrocephaly, Gorham–Stout phenomenon, fatal chylothorax, lymphatic malformation in thorax, venous malformation of liver and spleen years | *FLT4 c.2180C>T ,p.Ala727Val* | patient died of respiratory insufficiency at 2 | 6 |
| | | | | | P015 | Female | NR | Macrocephaly, fibrocystic disease of the breast, thyroid adenomas, subcutaneous hamartomas | *FLT4 c.2180C>T ,p.Ala727Val* | Mother of patient P014 | 6 |
| | | | | | P016 | Female | 7 years | Macrocephaly | *FLT4 c.2180C>T ,p.Ala727Val* | Sibling of patient P014 | 6 |
| | | | | | P017 | Male | 7 months | Macrocephaly | *FLT4 c.2180C>T ,p.Ala727Val* | Sibling of patient P014 | 6 |
| | | | | | P018 | Male | 39 years | DD, GI polyps | | | this study, PHTS Patient Registry UK (https://www.phts.org.uk) |
| | | | | | P019 | Male | 11 years | Macrocephaly, thyroid cyst, GI polyps | | | this study, PHTS Patient Registry UK (https://www.phts.org.uk) |
| | | | | | P020 | Male | 22 years | Macrocephaly, DD, hypotonia, thyroid cysts, oral papillomas, multiple skin lesions | | | this study, patient information provided by Dr Katherine Lachlan |
| | | | | | P021 | Female | 38 years | Macrocephaly, severe learning difficulties, acral keratosis, hypothyroidism, thyroid colloid nodules. Patient is asthmatic and on inhalers. Patient has had tonsillectomy and bilateral mastectomy | | | this study, patient information provided by Dr Isra Mohammed |
| | | | | | Additional 3 patients reported with this mutation , no phenotypic data avaialble | | | | | | 7-9 |
| *p.Arg173His* (R173H) | c.518G>A | NM_000314.8(PTEN):c.518G>A (p.Arg173His) | VCV000376032 | Pathogenic/likely pathogenic | P022 | Male | 9 years | Macrocephaly, DD, ASD, multiple cutaneous features | | | 1, clinical phenotypes updated for this study by Dr Katherine Lachlan |
| | | | | | P023 | Male | 43 years | Macrocephaly, tongue lesions, penile macules, GI polyps | | | 1 |
| | | | | | P024 | NR | NR | Macrocephaly, MR/DD | | | 10 |
| | | | | | P025 | Male | 10 | Macrocephaly, ASD, DD | | | 4,11 |
| | | | | | P026 | Male | NR | Macrocephaly | | Father of patient PO25 | 11 |
| | | | | | P027 | Male | 8 years | Macrocephaly, DD, ASD | | | 4,11 |
| | | | | | P028 | Female | 10 years | Macrocephaly, DD, Cafe-au-Lait spots, diffuse goiter | | Sibling of patient PO27 | 2 |
| | | | | | P029 | Female | 54 years | Invasive ductal carcinoma and ductal carcinoma in-situ of the breast | | only breast cancer data available | 12 |
| | | | | | P030 | Male | 3 years | Macrocephaly, DD, overgrowth, hypotonia | | | 13 |
| | | | | | P031 | Male | 4 years | Macrocephaly, DD, abnormal adenoids and tonsils, tongue papules, venous haemangioma | | | 14 |
| | | | | | P032 | Female | 35 years | Macrocephaly, ovarian cysts, joint hypermobility | | | 15 |
| | | | | | One additional patient reported with this mutation , no phenotypic data avaialble | | | | | | 16 |
| *p.Arg173Pro* (R173P) | c.518G>C | NM_000314.8(PTEN):c.518G>C (p.Arg173Pro) | VCV000185195 | Pathogenic/likely pathogenic | P033 | Female | 42 years | Macrocephaly, LDD, MR, DD, benign thyroid tumour, mucocutaneous lesions, facial papules, acral keratosis, palmoplantar keratoses, fibromas | | | 17 |
| | | | | | P034 | Female | 6 years | Macrocephaly, alopecia, nephritis, oral ulcers, submandibular lymphadenopathy, hepatosplenomegaly, haemagiomas, monogenic systemic lupus erythromatosis | *TALDO1 c.793C del: p. Gln265f* | | 18 |

DD: developmental delay
MR: Mental retardation
ASD: autism spectrum discorder
LDD: Lhermitte Duclos Disease
GI: gastrointestinal

**Supplementary table 2: Clinical findings of PHTS patients with nuclear excluded but catalytically active PTEN variants**

| *PTEN* variant | | | | | | PHTS patient data | | | | | | |
|---|---|---|---|---|---|---|---|---|---|---|---|---|
| **Amino acid change** | **Nucleotide change** | **HGVS nomenclature** | **ClinVar ID** | **ClinVar classification** | **Reference for nuclear exclusion** | **Patient ID assigned for this study only** | **Gender** | **Age** | **Clinical phenotypes** | **Other genetic alterations** | **Additional notes** | **Reference for patient data** |
| *p. Lys 13Glu* (K13E) | c.37A>G | NM_000314.8(PTEN):c.37A>G (p.Lys13Glu) | VCV000988004 | Likely pathogenic | 1 | P035 | NR | NR | Macrocephaly, mucocutaneous lesions, facial papules, oral papillomas, acral keratoses, GI polyps, genitourinary lesions, benign thyroid lesions | | | 7 |
| | | | | | | P036 | NR | NR | Macrocephaly, mucocutaneous lesions, facial papules, oral papillomas, acral keratoses, benign thyroid lesions, lung cancer | | | 7 |
| | | | | | | P037 | Male | NR | Macrocephaly, Mucocutaneous lesions, oral papillomas, penile freckling, vascular lesions/lipomas | | | 7 |
| | | | | | | P038 | Female | 8 years | Macrocephaly, DD, ASD | | | 8 |
| *p. Thre26Ile* (T26I) | c.77C>T | NM_000314.8(PTEN):c.77C>T (p.Thr26Ile) | VCV000189399 | Pathogenic/likely pathogenic | 2 | P039 | Male | 11years | Macrocephaly, ASD, MR, macular pigmentation of penis, bilateral gynecomastia, musculoskeletal alteration, obesity | | | 9 |
| | | | | | | P040 | Female | 1year | Macrocephaly, DD, capillary malformations on the limbs, facial AVM, | | | 10 |
| *p.Ile101Thr* (I101T) | c.302T>C | NM_000314.8(PTEN):c.302T>C (p.Ile101Thr) | VCV000822660 | Pathogenic/likely pathogenic | 3 | P041 | Female | 6 years | DD, tricholemmoma | | | 11 |
| | | | | | | P042 | Male | 6 years | Macrocephaly, DD, frequent upper respiratory tract infections, hypogammaglobulinemia | | | 12 |
| | | | | | | P043 | Male | 3 years | Macrocephaly, DD | | | 13 |
| | | | | | | P044 | Male | 10 years | Macrocephaly, DD, ASD,overgrowth | | | 3 |
| | | | | | | P045 | Female | 3 years | Macrocephlay, DD, ASD, dysmorphic features, generalized overgrowth, hemangioma of skin | | | 14 |
| | | | | | | P046 | Female | 3 years | Macrocephaly, enlarged perivascular spaces as seen on MRI, lumbar hypochromic spot | | | 15 |
| | | | | | | One additional patient reported with this mutation , no phenotypic data avaialble | | | | | | 16 |
| *p.Tyr177Asn* (Y177N) | c.529T>A | NM_000314.8(PTEN):c.529T>A (p.Tyr177Asn) | VCV000873327 | Variant of unknown significance | 2 | P047 | Male | 5 years | Macrocephaly, motor delay | | | 9 |
| *p.Phe241Ser* (F241S) | c.722T>C | NM_000314.8(PTEN):c.722T>C (p.Phe241Ser) | VCV000007850 | Likely pathogenic | 4 | P048 | Male | 2 years | DD, ASD, overgrowth, penile freckling | | | 17 |
| | | | | | | P049 | Male | 15 years | Macrocephaly, joint hypermobility, DD, thyroid nodule, cutaneous features | | | this study, PHTS Patient Registry UK (https://www.phts.org.uk) |
| *p.Asp252Gly* (D252G) | c.755A>G | NM_000314.8(PTEN):c.755A>G (p.Asp252Gly) | VCV000007849 | Pathogenic | 4 | P050 | Male | 3 years | Macrocephaly, DD, ASD | | | 17 |
| | | | | | | P051 | NR | NR | Macrocephaly, mucocutaneous lesions, facial papules, oral papillomas | | | 7 |
| *p.Gln261Glu* (Q261E) | c.781C>G | NM_000314.8(PTEN):c.781C>G (p.Gln261Glu) | VCV000873325 | Variant of unknown significance | 2 | P052 | Male | 34 years | Macrocephaly, lipomas, penile freckling, colorectal polyps, general overgrowth. | | | 9 |
| *p.Trp274Leu* (W274L) | c.821G>T | NM_000314.8(PTEN):c.821G>T (p.Trp274Leu) | VCV000427600 | Pathogenic/likely pathogenic | 4 | P053 | Female | 6 years | Macrocephaly, DD, scoliosis | | | 18,19 |
| | | | | | | P054 | Female | NR | Macrocephaly, DD | | mother of P046 | 19 |
| *p.Asn276Ser* (N276S) | c.827A>G | NM_000314.8(PTEN):c.827A>G (p.Asn276Ser) | VCV002502345 | Pathogenic | 4 | P055 | NR | NR | Macrocephaly, ASD | | | 20 |
| *p.Thr277Ala* (T277A) | c.829A>G | NM_000314.8(PTEN):c.829A>G (p.Thr277Ala) | VCV000571869 | Pathogenic | 5 | P056 | Male | 9 years | Macrocephaly, general developmental disorder, speech delay, general overgrowth | | | 9 |
| *p.Lys289Glu* (K289E) | c.865A>G | NM_000314.8(PTEN):c.865A>G (p.Lys289Glu) | VCV000142912 | Variant of unknown significance | 1 | P057 | Female | 40 years | Facial papillomas, fibrocystic disease in the breast, lipoma, hamartomatous polyps in the stomach, GI polyps and GI adenoma | | | 21 |
| | | | | | | P058 | Female | 14 years | Facial papillomas, lipoma, hamartomatous polyps in the stomach, GI polyps | | | 21 |
| *p.Asp310Gly* (D310G) | c.929A>G | NM_000314.8(PTEN):c.929A>G (p.Asp310Gly) | VCV000566673 | Likely pathogenic | 2 | P059 | Male | 29 years | Macrocephaly, papules, palmoplantar keratoses, thyroid adenomas, papillary-follicular thyroid cancer, GI polyps , testicular cancer. | | | 9 |
| *p.Phe341Val* (F341V) | c.1021T>G | NM_000314.8(PTEN):c.1021T>G (p.Phe341Val) | VCV000495806 | Conflicting classifications of pathogenicity | 6 | One patient reported with this mutation, no patient phenotypic data available | | | | | | 22 |

DD: developmental delay
MR: Mental retardation
ASD: autism spectrum discorder
LDD: Lhermitte Duclos Disease
GI: gastrointestinal

**Supplementary table 3: Reason for euthanasia of *Pten* $^{+/+}$, *Pten* $^{+/-}$ and *Pten* $^{+/R173C}$ mice**

| | *Pten* $^{+/+}$ | | | | | *Pten* $^{+/-}$ | | | | | *Pten* $^{+/R173C}$ | | | | |
|---|---|---|---|---|---|---|---|---|---|---|---|---|---|---|---|
| **Reason for euthanasia** | **Palpable masses** | **Ill health** | **Both masses & ill health** | **Found dead** | **End of study** | **Palpable masses** | **Ill health** | **Both masses & ill health** | **Found dead** | **End of study** | **Palpable masses** | **Ill health** | **Both masses & ill health** | **Found dead** | **End of study** |
| Female | 0/32<br>0% | 12/32<br>37% | 0/32<br>0% | 0/32<br>0% | 20/32<br>63% | 15/19<br>79% | 4/19<br>21% | 0/19<br>0% | 0/19<br>0% | 0/19<br>0% | 13/26<br>50% | 12/26<br>46% | 1/26<br>4% | 0/26<br>0% | 0/26<br>0% |
| Male | 0/25<br>0% | 15/25<br>60% | 0/25<br>0% | 0/25<br>0% | 10/25<br>40% | 14/19<br>74% | 5/19<br>26% | 0/19<br>0% | 0/19<br>0% | 0/19<br>0% | 4/19<br>21% | 11/19<br>58% | 0/19<br>0% | 0/19<br>0% | 4/19<br>21% |

**Supplementary table 4: Frequency and grade of lesions in male mice**

| | | | Frequency and grade of lesions in male mice | | | | | |
|---|---|---|---|---|---|---|---|---|
| | | | *Pten* $^{+/+}$ | | *Pten* $^{+/-}$ | | *Pten* $^{+/R173C}$ | |
| | | **Median survival age** | **681 days** | | **310 days** | | **585 days** | |
| **Tissue** | **Finding** | **Site** | **Incidence** | **%** | **Incidence** | **%** | **Incidence** | **%** |
| **Small intestine** | *Hyperplasia* | *Focal and crypt* | 1/16 | 6.2 | 5/18 | 27.7 | 2/14 | 14.2 |
| | *Adenoma* | | 0/16 | 0 | 3/18 | 16.6 | 0/14 | 0 |
| **Stomach** | *Hyperplasia* | *Glandular* | 9/16 | 56.2 | 5/18 | 27.7 | 6/14 | 42.8 |
| **Thyroid** | *Hyperplasia, focal* | | 0/13 | 0 | 11/18 | 61.1 | 0/13 | 0 |
| | *Adenoma* | | 0/13 | 0 | 8/18 | 44.4 | 0/13 | 0 |
| | *cytoplasmic inclusions* | | 0/13 | 0 | 0/18 | 0 | 1/13 | 7.6 |
| **Adrenal gland** | *Hyperplasia* | *Medulla* | 2/16 | 15.3 | 2/17 | 11.7 | 2/13 | 15.3 |
| | *Hyperplasia* | *Subcapsular* | 1/16 | 7.6 | 0/17 | 0 | 0/13 | 0 |
| | *Phaeochromocytoma* | | 0/16 | 0 | 14/17 | 82.3 | 9/13 | 69.2 |
| **Lung** | *Bronchioalveolar adenoma* | | 2/16 | 12.5 | 0/18 | 0 | 5/14 | 35.7 |
| | *Bronchoalevolar carcinoma* | | 3/16 | 18.7 | 0/18 | 0 | 0/14 | 0 |
| **Prostate** | *LG-PINs* | | 1/16 | 0 | 4/17 | 23.5 | 4/14 | 28.5 |
| | *HG-PINs* | | 0/16 | 0 | 6/17 | 35.2 | 0/14 | 0 |
| | *Adencocarcinoma* | | 0/16 | 0 | 1/17 | 5.8 | 0/14 | 0 |

**Supplementary table 5: Frequency and grade of lesions in female mice**

| | | | Frequency and grade of lesions in female mice | | | | | |
|---|---|---|---|---|---|---|---|---|
| | | | *Pten* $^{+/+}$ | | *Pten* $^{+/-}$ | | *Pten* $^{+/R173C}$ | |
| | | **Median survival age** | **Undefined** | | **179 days** | | **302 days** | |
| **Tissue** | **Finding** | **Site** | **Incidence** | **%** | **Incidence** | **%** | **Incidence** | **%** |
| **Small intestine** | *Hyperplasia* | *Focal and crypt* | 1/23 | 4.3 | 6/19 | 31.5 | 0/22 | 0 |
| | *Adenoma* | | 0/23 | 0 | 4/19 | 21 | 0/22 | 0 |
| **Stomach** | *Hyperplasia* | *Glandular* | 12/23 | 52.1 | 3/19 | 15.7 | 7/22 | 31.8 |
| **Thyroid** | *Hyperplasia, focal* | | 0/18 | 0 | 4/15 | 26.6 | 0/19 | 0 |
| | *Adenoma* | | 0/18 | 0 | 3/15 | 20 | 0/19 | 0 |
| | *cytoplasmic inclusions* | | 6/18 | 33.3 | 0/15 | 0 | 9/19 | 47.3 |
| **Adrenal gland** | *Hyperplasia* | *Medulla* | 0/17 | 0 | 3/18 | 16.6 | 1/19 | 5.2 |
| | *Hyperplasia* | *Subcapsular* | 6/17 | 35.2 | 1/18 | 5.5 | 4/19 | 21 |
| | *Phaeochromocytoma* | | 0/17 | 0 | 12/18 | 66.6 | 5/19 | 26.3 |
| **Lung** | *Bronchioalveolar adenoma* | | 3/23 | 13 | 0/18 | 0 | 0/22 | 0 |
| | *Bronchioalevolar carcinoma* | | 1/23 | 4.3 | 0/18 | 0 | 0/22 | 0 |
| **Mammary** | *Lymphoid infiltrate* | | 2/23 | 8.6 | 0/19 | 0 | 0/22 | 0 |
| | *Adenomyoepithelioma (AME)* | | 0/23 | 0 | 2/19 | 10.4 | 0/22 | 0 |
| | *Malignant tumour* | | 0/23 | 0 | 1/19 | 5.2 | 1/22 | 4.5 |
| **Uterus** | *Cystic hyperplasia* | *Endometrium* | 10/23 | 43.4 | 1/19 | 5.2 | 3/22 | 13.6 |
| | *Atypical hyperplasia* | *Endometrium* | 1/23* | 4.3 | 19/19 | 100 | 1/28** | 3.5 |
| | *Carcinoma* | *Endometrium* | 0/23 | 0 | 0/19 | 0 | 0/28 | 0 |
| | *Decidualisation* | *Focal* | 0/23 | 0 | 6/19 | 31.5 | 0/28 | 0 |
| **Ovary** | *Haemorrhagic cyst* | | 2/21 | 9.5 | 0/19 | 0 | 0/28 | 0 |
| | *Cyst* | | 2/21 | 9.5 | 1/19 | 5.2 | 0/22 | 0 |

* age of mouse was 647 days
**age of mouse was 300 days

**Supplementary table 6: Frequency and grade of genetic background specific and immune findings in male mice**

| | | | Frequency and grade of immune findings in male mice | | | | | |
|---|---|---|---|---|---|---|---|---|
| | | | *Pten* $^{+/+}$ | | *Pten* $^{+/-}$ | | *Pten* $^{+/R173C}$ | |
| | | **Median survival age** | **681 days** | | **310 days** | | **585 days** | |
| **Tissue** | **Finding** | **Site** | **Incidence** | **%** | **Incidence** | **%** | **Incidence** | **%** |
| **Small intestine** | *Lymphoma* | | 1/16 | 6.2 | 1/18 | 5.5 | 2/14 | 14.2 |
| | *Lymphoid Hyperplasia* | | 0/16 | 0 | 0/18 | 0 | 1/14 | 7.1 |
| **Large Intestine** | *Lymphoma* | | 1/16 | 6.2 | 0/18 | 0 | 0/14 | 0 |
| | *Lymphoid Hyperplasia* | | 0/16 | 0 | 0/18 | 0 | 0/14 | 0 |
| **Pancreas** | *Lymphoma* | | 0/16 | 0 | 0/18 | 0 | 1/14 | 7.1 |
| | *Lymphoid infiltrates* | | 4/16 | 25 | 0/18 | 0 | 11/14 | 78.5 |
| **Liver** | *Inflammatory cell infiltrate* | *Focal* | 4/16 | 25 | 2/18 | 11.1 | 3/14 | 21.4 |
| | *Inflammatory cell infiltrate* | *Periportal* | 0/16 | 0 | 0/18 | 0 | 0/14 | 0 |
| | *Lymphoid infiltrates* | *Perivascular* | 0/16 | 0 | 0/18 | 0 | 0/14 | 0 |
| | *Lymphoma* | | 0/16 | 0 | 0/18 | 0 | 0/14 | 0 |
| **Kidney** | *Inflammatory cell infiltrate* | *Pelvis* | 12/16 | 75 | 15/18 | 83.3 | 14/14 | 100 |
| | *Lymphoid infiltrates* | | 0/16 | 0 | 0/18 | 0 | 0/14 | 0 |
| | *Lymphoma* | | 0/16 | 0 | 0/18 | 0 | 0/14 | 0 |
| **Lung** | *Lymphoid infiltrate* | *Peribronchiolar* | 7/16 | 43.7 | 4/18 | 22.2 | 9/14 | 64.2 |
| | *Lymphoma* | | 0/16 | 0 | 0/18 | 0 | 1/14 | 7.1 |
| **Spleen** | *Haematopoiesis, extramedullary* | | 16/16 | 100 | 18/18 | 100 | 13/14 | 92.8 |
| | *Myelopoiesis* | | 1/16 | 6.25 | 2/18 | 11.1 | 1/14 | 7.1 |
| | *Lymphoid hyperplasia* | | 0/16 | 0 | 1/18 | 5.5 | 1/14 | 7.1 |
| | *Lymphoma* | | 0/16 | 0 | 0/18 | 0 | 5/14 | 35.7 |
| **Thymus** | *Lymphoid hyperplasia* | | 2/13 | 15.3 | 5/15 | 33.3 | 8/9 | 88.8 |
| | *Lymphoma* | | 0/13 | 0 | 0/15 | 0 | 0/9 | 0 |
| **Prostate** | *Lymphoid infiltrate* | | 1/16 | 6.25 | 0/18 | 0 | 3/14 | 21.4 |
| | *Lymphoma* | | 0/16 | 0 | 0/18 | 0 | 0/14 | 0 |
| **Lymph nodes** | *Lymphoid hyperplasia* | *all tissues* | 1/16 | 6.2 | 16/18 | 88.8 | 9/14 | 64.2 |
| | *Lymphoma* | | 3/16 | 18.7 | 3/18 | 16.6 | 2/14 | 14.2 |

**Supplementary table 7: Frequency and grade genetic background specific and immune findings in female mice**

| | | | Frequency and grade of immune findings in female mice | | | | | |
|---|---|---|---|---|---|---|---|---|
| | | | ***Pten*** $^{+/+}$ | | ***Pten*** $^{+/-}$ | | ***Pten*** $^{+/R173C}$ | |
| | | **Median survival age** | **Undefined** | | **179 days** | | **302 days** | |
| **Tissue** | **Finding** | **Site** | **Incidence** | **%** | **Incidence** | **%** | **Incidence** | **%** |
| **Small intestine** | *Lymphoma* | | 0/23 | 0 | 0/19 | 0 | 0/22 | 0 |
| | *Lymphoid Hyperplasia* | | 0/23 | 0 | 0/19 | 0 | 3/22 | 13.6 |
| **Large intestine** | *Lymphoma* | | 0/23 | 0 | 0/19 | 0 | 0/22 | 0 |
| | *Lymphoid Hyperplasia* | | 0/23 | 0 | 0/19 | 0 | 2/22 | 9 |
| **Pancreas** | *Lymphoma* | | 0/23 | 0 | 0/19 | 0 | 0/22 | 0 |
| | *Lymphoid infiltrate* | | 14/23 | 60.8 | 1/19 | 5.2 | 15/22 | 68.1 |
| **Liver** | *Inflammatory cell infiltrate* | *Focal* | 8/23 | 34.7 | 5/19 | 26.3 | 1/22 | 4.5 |
| | *Inflammatory cell infiltrate* | *Periportal* | 0/23 | 0 | 0/19 | 0 | 0/22 | 0 |
| | *Lymphoid infiltrates* | *Perivascular* | 2/23 | 8.6 | 0/19 | 0 | 7/22 | 31.8 |
| | *Lymphoma* | | 1/23 | 4.3 | 0/19 | 0 | 0/22 | 0 |
| **Kidney** | *Inflammatory cell infiltrate* | *Pelvis* | 20/23 | 86.9 | 12/15 | 80 | 18/22 | 81.8 |
| | *Lymphoid infiltrates* | | 0/23 | 0 | 0/15 | 0 | 6/22 | 27.2 |
| | *Lymphoma* | | 1/23 | 4.3 | 0/15 | 0 | 0/22 | 0 |
| **Lung** | *Lymphoid infiltrate* | *Peribronchiolar* | 17/23 | 73.9 | 3/18 | 16.6 | 21/22 | 95.4 |
| | *Lymphoma* | | 2/23 | 8.6 | 0/18 | 0 | 0/22 | 0 |
| **Spleen** | *Haematopoiesis, extramedullary* | | 23/23 | 100 | 19/19 | 100 | 22/22 | 100 |
| | *Myelopoiesis* | | 1/23 | 4.3 | 0/19 | 0 | 0/22 | 0 |
| | *Lymphoid hyperplasia* | | 0/23 | 0 | 0/19 | 0 | 9/22 | 40.9 |
| | *Lymphoma* | | 3/23 | 13 | 0/19 | 0 | 0/22 | 0 |
| **Thymus** | *Lymphoid hyperplasia* | | 7/11 | 63.6 | 3/15 | 20 | 12/28 | 66.6 |
| | *Lymphoma* | | 2/11 | 18.1 | 0/15 | 0 | 0/18 | 0 |
| **Lymph nodes** | *Lymphoid hyperplasia* | *all tissues* | 3/23 | 13 | 18/19 | 94.7 | 17/22 | 77.2 |
| | *Lymphoma* | | 2/23 | 8.6 | 0/19 | 0 | 0/22 | 0 |

**Supplementary table 8: DNA constructs**

| Plasmid construct | Purpose |
|---|---|
| pHR SIN IRES Puro PTEN wild-type, C124S, R173C, R173H, D252G, K66R, R173C K66R, R14G, D24V, G36R, H61R, I67R, M134I, C136R, W274L | Lentiviral expression of PTEN wild-type and mutants in U-87 MG cells |
| pHR SIN GFP | Lentiviral expression of GFP in U-87 MG cells |
| pHR SIN GFP-ITK PH-GRP1 PH | Lentiviral expression of GFP-ITK PH-GRP1 PH domains in U-87 MG cells |
| p-CMV-VSV-G, pCMV-dR8.91 and pCMV-dR8.2 | Packaging vectors for lentiviral production |
| pGEX-6P-1 PTEN wild-type, C124S, D252G, R173C | Expression of GST tagged PTEN wild-type and mutants in *E. coli* for recombinant protein purification |
| pEBG2T empty, PTEN wild-type, C124S, R173C | Expression of GST tagged PTEN wild-type and mutants in mammalian cells for recombinant protein purification |
| EGFPN3 PTEN wild-type, R173C | Expression of PTEN wild-type and R173C with a C-terminal GFP Tag in mammalian cells |
| Rluc-PTEN-YFP wild-type, Rluc-PTENR173C-YFP, Rluc-PTENR173H-YFP | Expression of intramolecular BRET-based fusions for conformational readouts and live cell imaging, in HEK and Hela PTEN KO cells |

**Supplementary table 9: Primers for site-directed mutagenesis**

| PTEN mutation | Primers for site directed mutagenesis | |
|---|---|---|
| | **Forward Primer** | **Reverse Primer** |
| **R173C** | 5’TTCCCAGTCAGAGGtgcTATGTGTATTATTATAGC 3’ | 5’GCTATAATAATACACATAGCACCTCTGACTGGGAA3’ |
| **R173H** | 5’ CAGTCAGAGGcacTATGTGTATTATTATAG 3’ | 5’ GGAATAGTTACTCCCTTTTTG 3’ |

**Supplementary table 10: Primers used for genotyping**

| Mouse line | Primers for genotyping | Expected size of PCR product |
|---|---|---|
| ***Pten*** $^{+/-}$ | *Pten* common: 5’ TTGCACAGTATCCTTTTGAAG 3’ | Wild-type: 240bp |
| | *Pten* WT: 5’ GTCTCTGGTCCTTACTTCC 3’ | Mutant: 320bp |
| | *Pten* Neo: 5’ ACGAGACTAGTGAGACGTGC 3’ | |
| ***Pten*** $^{+/R173C}$ | *Pten* R173C F: 5’ GTAAACAGGTTTGTGAAGACTTGC 3’ | iCre 250S F: 5’ GAG GGA CTA CCT CCT GTA CC 3’<br>Wild-type: 564bp |
| | *Pten* R173C R: 5’ AAGTTTTCCGACACACAGACAG 3’ | Mutant: 397bp and 167bp (seen after EcoNI digestion) |
| ***Pten*** $^{flox/flox}$ | *Pten* flox F: 5’ CAAGCACTCTGCGAACTGAG 3’ | Wild-type: 156bp |
| | *Pten* flox R: 5’ AAGTTTTTGAAGGCAAGATGC 3’ | Mutant: 328bp |
| ***Cag-Cre-ER*** $^{T2}$ | CreERT2 F: 5’ GAATGTGCCTGGCTAGAGATC 3’ | PCR product of transgene at 190bp |
| | CreERT2 R: 5’ GCAGATTCATCATGCGGA 3’ | |
| ***Emx1-Cre*** | iCre 250S F: 5’ GAGGGACTACCTCCTGTACC 3’ | PCR product of transgene at 630bp |
| | iCre 880AS R: 5’ TGCCCAGAGTCATCCTTGGC 3’ | |

**Supplementary table 11: Antibodies used**

**A. Antibodies used for western blotting**

| **Antibody** | **Supplier** | **Catalogue number** | **Species reactivity** | **Dilution** |
|---|---|---|---|---|
| PTEN | Cell Signaling Technology | 9552 | Rabbit | 1:1000 |
| AKT P-S473 | Cell Signaling Technology | 9271 | Rabbit | 1:1000 |
| AKT P-T308 | Cell Signaling Technology | 9275 | Rabbit | 1:500 |
| AKT | Cell Signaling Technology | 9272 | Rabbit | 1:1000 |
| S6 P-S240/244 | Cell Signaling Technology | 2215 | Rabbit | 1:1000 |
| S6 P-S235/236 | Cell Signaling Technology | 2211 | Rabbit | 1:1000 |
| S6 | Cell Signaling Technology | 2217 | Rabbit | 1:1000 |
| p70 S6K P-T389 | Cell Signaling Technology | 9205 | Rabbit | 1:1000 |
| p70 S6K | Cell Signaling Technology | 9202 | Rabbit | 1:1000 |
| PTEN P-380/82/83 | Cell Signaling Technology | 9554 | Rabbit | 1:1000 |
| PRAS40 P-T246 | Cell Signaling Technology | 2640 | Rabbit | 1:1000 |
| PRAS40 | Cell Signaling Technology | 2610 | Rabbit | 1:1000 |
| FoxO1 P-T24/FoxO3a P-T32 | Cell Signaling Technology | 9464 | Rabbit | 1:1000 |
| FoxO3a | Cell Signaling Technology | 2497 | Rabbit | 1:1000 |
| γH2AX | Sigma-Aldrich | 05-636 | Mouse | 1:1000 |
| Vinculin | Sigma-Aldrich | V-9131 | Mouse | 1:10,000 |
| GAPDH | Cell Signaling Technology | 2218 | Rabbit | 1:10,000 |
| GFP | Cell Signaling Technology | 2956 | Rabbit | 1:2000 |

**B. Antibodies used for IHC**

| **Antibody** | **Supplier** | **Catalogue number** | **Species reactivity** | **Dilution** |
|---|---|---|---|---|
| PTEN | Cell Signaling Technology | 9559 | Rabbit | 1:100 |
| pAKT Ser473 | Cell Signaling Technology | 4060 | Rabbit | 1:50 |
| NeuN | Millipore | MAB377 | Mouse | 1:300 |
| GFAP | Sigma Aldrich | C9205 | Mouse | 1:1000 |
| Olig2 | Merck-Millipore | AB9610 | Rabbit | 1:500 |

**C. Antibodies used for flow cytometry**

| **Antibody** | **Fluorochrome** | **Clone** | **Surface/intracellular** | **Supplier** | **Catalogue number** | **Dilution** |
|---|---|---|---|---|---|---|
| B220 | AF700 | RA36B2 | Surface | Biolegend | 103231 | 1:100 |
| TCRβ | BV711 | H57-597 | Surface | Biolegend | 109243 | 1:100 |

**Supplementary table 12: qRT PCR primers**

| Gene | Primers for qRT PCR |
|---|---|
| ***PTEN*** | *PTEN* EXON 5 F: 5’ CTGCAGAGTTGCACAGTATCC 3’ |
| | *PTEN* EXON 5 R: 5’ GAATTGTGACTCCCTTTTTGTCTCTGG 3’ |
| ***GAPDH*** | *GAPDH* F: 5’ GTGAAGGTCGGAGTCAACGG 3’ |
| | *GAPDH* R: 3’ GAGGGATCTCGCTCCTGGAA 3’ |
